# MR-LDP: a two-sample Mendelian randomization for GWAS summary statistics accounting for linkage disequilibrium and horizontal pleiotropy

**DOI:** 10.1101/684746

**Authors:** Qing Cheng, Yi Yang, Xingjie Shi, Kar-Fu Yeung, Can Yang, Heng Peng, Jin Liu

## Abstract

The proliferation of genome-wide association studies (GWAS) has prompted the use of two-sample Mendelian randomization (MR) with genetic variants as instrumental variables (IV) for drawing reliable causal relationships between health risk factors and disease outcomes. However, the unique features of GWAS demand that MR methods account for both linkage disequilibrium (LD) and ubiquitously existing horizontal pleiotropy among complex traits, which is the phenomenon wherein a variant affects the outcome through mechanisms other than exclusively through the exposure. Therefore, statistical methods that fail to consider LD and horizontal pleiotropy can lead to biased estimates and false-positive causal relationships. To overcome these limitations, we propose a probabilistic model for MR analysis to identify the casual effects between risk factors and disease outcomes using GWAS summary statistics in the presence of LD and to properly account for horizontal pleiotropy among genetic variants (MR-LDP). MR-LDP utilizes a computationally efficient parameter-expanded variational Bayes expectation-maximization (PX-VBEM) algorithm to estimate the parameter of interest and further calibrates the evidence lower bound (ELBO) for a likelihood ratio test. We then conducted comprehensive simulation studies to demonstrate the advantages of MR-LDP over the existing methods in terms of both type-I error control and point estimates. Moreover, we used two real exposure-outcome pairs (CAD-CAD and Height-Height; CAD for coronary artery disease) to validate the results from MR-LDP compared with alternative methods, showing that our method is more efficient in using all instrumental variants in LD. By further applying MR-LDP to lipid traits and body mass index (BMI) as risk factors for complex diseases, we identified multiple pairs of significant causal relationships, including a protective effect of high-density lipoprotein cholesterol (HDL-C) on peripheral vascular disease (PVD), and a positive causal effect of body mass index (BMI) on hemorrhoids.

## 1 Introduction

Epidemiological studies have contributed tremendously to improving our understanding of the primary causes of complex diseases. However, numerous cases of significant associations from observational studies have been subsequently contradicted by large clinical trials [1, 2]. Drawing causal inferences from observational studies is particularly challenging because of unmeasured confounding, reverse causation and selection bias [3, 4]. Although the randomized controlled trial (RCT) is considered a gold standard to evaluate causality in an exposure-outcome pair, RCTs have certain limitations including impracticality (no intervention may exist), high expense, and ethical issues [5]. Fortunately, as germline genetic variants (single nucleotide polymorphisms, SNPs) are fixed after random mating and cannot be modified by subsequent factors, e.g., environment factors and living styles, Mendelian randomization (MR) uses genetic variants as instruments to examine the causal effects between health risk factors and disease outcomes, largely excluding the influence from unobserved confounding factors [3]. In the past decade, a large number of genome-wide association studies (GWAS) have been successfully used to identify genetic variants associated with complex traits at the genome-wide significance level, including both health factors and diseases, e.g., lipids, BMI, and type-2 diabetes, and most of completed GWAS are simply observational studies instead of RCTs. The results from completed GWAS are mostly publicly accessible, e.g., GWAS Catalog outlines a list of sources for summary statistics (https://www.ebi.ac.uk/gwas/downloads/summary-statistics). This large amount of publicly available GWAS summary statistics has prompted the widespread use of two-sample MR as an efficient and cost-effective method to interrogate the causal relationships among health risk factors and disease outcomes.

MR is closely related to the instrumental variable (IV) methods, which have a long history of use in econometrics [6]. Classically, an inverse-variance weighted (IVW) and a likelihood-based approach have been used for two-sample MR analysis with summary-level data [7]. These methods must strictly obey assumptions for MR, including two most fundamental ones:

1. IVs affect the outcome exclusively through the risk exposures.
2. IVs are independent from each other, or in a GWAS context, instrumental variants are not in LD.

The first assumption is also referred to as exclusion restriction assumption or no horizontal pleiotropy. The violation of this assumption can distort the statistical inference for MR analysis, leading to biased estimates and false-positive causal relationships. Recent comprehensive surveys reported persuasive pleiotropy among complex traits [8, 9], such as autoimmune diseases [10] and psychiatric disorders [11]. Consequently, methods that do not account for pleiotropy can substantially reduce the power and inflate the false-positive discoveries. To address this issue, sisVIVE was proposed in the presence of individual-level data [12]. To further relax this assumption for two-sample MR analysis using summary-level data, various statistical methods have been proposed and we divide them into two categories. The first group consists of step-wise methods to correct the impact of horizontal pleiotropy. These methods first detect and remove SNPs with horizontal pleiotropy, and MR analysis is performed in the subsequent step, including Q test [13], Cook’s distance [14], Studentized residuals [14], GSMR [15], and MR-PRESSO [16]. The drawback of this type of methods is that the number of SNPs after removal is limited given that abundant pleiotropy exists among complex traits, which can substantially reduce the statistical power to detect the causal relationships. In contrast, the second group of methods jointly estimate causal effects by taking into account the horizontal pleiotropy, e.g., MR-Egger [17], MRMix [18] and RAPS [19]. Compared to MR-Egger, RAPS further addressed the measurement error issues, where most of existing methods applicable to GWAS summary statistics assume that sampling error from SNP-exposure is negligible [20].

On the other hand, minimal literature is available regarding the relaxation of the second assumption above. Among the methods mentioned above, only GSMR is capable of accounting weak or moderate linkage disequilibrium (LD) for SNPs, while others demand all instrumental SNPs to be independent, which is typically achieved by conducting SNP pruning and thus reducing the number of instrumental variants for follow-up MR analysis. As SNPs within proximity tend to be highly correlated, MR methods not accounting for LD structure may substantially lose statistical power due to the pruning process to obtain independent SNPs. Moreover, GSMR is a step-wise method to remove instrumental variants with horizontal pleiotropy, making it less powerful due to the removal of invalid variants.

In this paper, we propose a statistically unified and efficient two-sample <u>MR</u> method to utilize all weak instruments within <u>LD</u> (MR-LD), and further consider a MR-LD accounting for horizontal <u>P</u>leiotropy (MR-LDP). Similar to RAPS, MR-LDP does not require the no measurement error assumption. The key idea is to build a joint probabilistic model for GWAS summary statistics from both exposure and outcome using a reference panel to reconstruct LD among instrumental variants and to conduct a formal hypothesis testing to make inferences about the causal effect that links the exposure and the outcome through a linear relationship. We also develop an efficient variational Bayesian expectation-maximization algorithm accelerated by using the parameter expansion (PX-VBEM) to estimate the causal effect for MR-LD and MR-LDP. Moreover, we calibrate the evidence lower bound (ELBO) to construct the likelihood ratio test for the evaluation of statistical significance of the estimated effect. Simulation studies show that MR-LDP outperforms competing methods in terms of type-I error control and point estimates for making causal inference. Additionally, we used two real exposure-outcome pairs to validate results from MR-LD and MR-LDP compared with alternative methods, particularly showing our methods more efficiently use all SNPs in LD. By further applying MR-LDP to summary statistics from GWAS, we identified multiple pairs of significant causal relationships, including a protective effect of high-density lipoprotein cholesterol (HDL-C) on peripheral vascular disease (PVD), and a positive causal effect of BMI on hemorrhoids.

## 2 Material and Methods

### 2.1 Reference panel data

As MR-LD and MR-LDP use the marginal effect sizes and their standard errors from GWAS summary statistics to build a probabilistic model for making causal inference, information regarding correlations among SNPs is missing, i.e., LD denoted as **R** is missing. Thus, we need to use a reference panel dataset to assist with reconstructing LD. In this study, given that we primarily focus on European populations, we choose to use samples from the following resource as the external reference panel: UK10K Project (Avon Longitudinal Study of Parents and Children, ALSPAC; TwinsUK) merged with 1000 Genome Project Phase 3 (*N* = 4, 284), which is denoted UK10K thereafter. As SNPs from HapMap Project Phase 3 (HapMap3) are more reliable, we choose to limit our analysis using SNPs from HapMap3 (*p* = 1, 189, 556).

As samples from ALSPAC and TwinsUK include populations other than European, we conducted strict quality control for UK10K data using PLINK [21]. First, SNPs were excluded from the analysis if their calling rates were less than 95%, minor allele frequencies were less than 0.01, or *p*-values were less than 1 × 10^*-*6^ in the Hardy-Weinberg equilibrium test. We then removed the individuals with genotype missing rates greater than 5%. To further remove individuals with high relatedness in all samples, we used GCTA [22] to first identify those individual pairs with estimated genetic relatedness greater than 0.05 and then randomly remove one from such a pair. Additionally, we carried out the principal components analysis (PCA) on the individuals to identify the population stratification [23]. In this way, we extracted the clustering subgroup representing the major European ancestry using hierarchical clustering on principal components(HCPC) approach [24]. Finally, there were 3,764 individuals remained with 989,932 SNPs.

### 2.2 Choice of LD matrix

Since the LD between two SNPs decays exponentially with respect to their distance, we use LDetect [25] to partition the whole genome into *L* blocks first and then calculate the estimated correlation matrix in each block. For each block, we adopt a shrinkage method to guarantee the sparsity and positive definiteness of the estimated correlation matrix [26]. In particular, the correlation matrix estimator 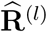 in each block is obtained by optimizing as follows

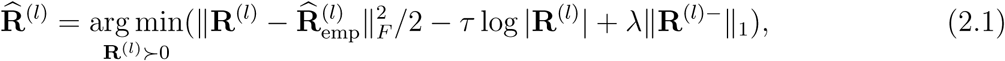

where 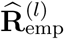 is the empirical correlation matrix in the *l*-th block, *λ* ≥ 0 is the shrinkage tuning parameter, and the lasso-type penalty ensures a sparse solution. In addition, *τ* > 0 is fixed at a small value and the logarithmic barrier term is used to enforce a positive-definite solution. More details can be found in [26]. A corresponding R package named *PDSCE* is available to complete the estimation process. In addition, we fix the shrinkage parameter *λ* to be 0.055 in simulation studies and vary *λ* ∈ {0.1, 0.15} in real data analysis.

### 2.3 Likelihood for summary statistics

Before elaborating on our method, we first review the following multiple linear regression model that links a trait to genotype data:

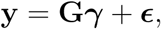

where **y** is an *n* × 1 vector for trait among *n* individuals, **G** is an *n* × *p* matrix for genotypes, ***γ*** is a *p* × 1 vector for effect sizes, and ***ϵ*** is the vector for random noises. Suppose that the individual-level data *{***G, y***}* are not accessible, but the summary statistics 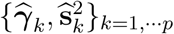 from univariate linear regression are available:

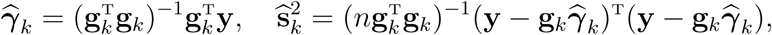

where **g**_*k*_ is the *k*-th column of 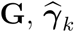 and 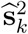 are estimated effect sizes and its variance for SNP *k*, respectively. 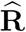 denotes the correlation among all genotyped SNPs and 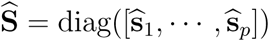, which is a diagonal matrix for corresponding standard errors. Provided that sample size *n* is large enough and the trait is highly polygenic (i.e., the squared correlation coefficient between the trait and each genetic variant is close to zero), we can use the following formula to approximate the distribution of ***γ*** based on the summary statistics in a similar fashion as [27, 28, 29, 30]:

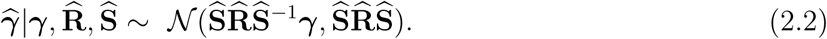

Analogously, we apply this distribution to the two-sample MR analysis. The summary statistics for SNP-exposure and SNP-outcome are denoted by 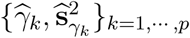 and 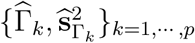 respectively. Therefore, the likelihood for two-sample summary statistics can be written as:

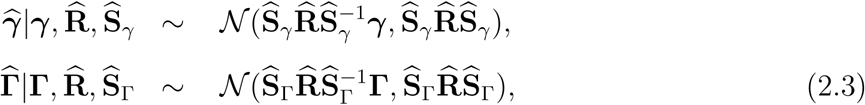

where 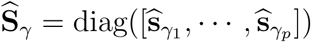 and 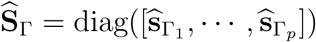 are both diagonal matrices. In this formulation, the correlations among all *p* SNPs, 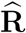, are not estimable from summary statistics itself. Zhu and Stephens [29] showed that 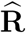 could be replaced with 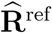 that is estimated from independent samples, where the difference in log-likelihood between individual-level data and summary statistics is a constant that does not depend on the effect size assuming that polygenicity holds and the sample size of individual-level data is large. Thus, distributions for summary statistics (2.3) will produce approximately the same inferential results as its counterpart for individual-level data. Hereafter, we use 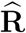 implicitly for 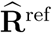 and details on estimating 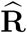 can be found in Section 2.2.

### 2.4 MR-LDP model overview

The fundamental assumptions for two-sample MR analysis include the independence among instrumental variables, and three IV assumptions for a genetic instrument: (1) associated with health risk factors (*γ* ≠ 0); (2) independent of unobserved confounding factors between the risk factors and the disease outcomes; (3) independent of **Y** given risk factors and confounders. Given strong LD structure among SNPs and abundant horizontal pleiotropy in GWAS, these unique features invalidate the independence assumption for genetic variants and two IV assumptions (2) and (3). Our proposed MR-LDP aims to make causal inference of the risk factors on a disease outcome using a probabilistic model by accounting for both the LD structure and the influence of horizontal pleiotropy as depicted in Figure 1. We first utilize an approximated likelihood to depict the distribution of correlated SNPs from GWAS summary statistics for the risk exposure and the disease outcome, respectively, as shown Equation (2.3). Given *p* instrumental variants, the inputs for MR-LDP are GWAS summary statistics for SNP-exposure and SNP-outcome, respectively, and a genotype reference panel (Figure 1A). By introducing an additional random effect ***α***, we would further eliminate the variance in the disease outcome due to pervasive horizontal pleiotropy. Since MR-LDP uses an approximated likelihood to jointly delineate the distribution for summary statistics (i.e., estimated effect sizes and their standard errors) from GWAS, it is free of the assumption for no measurement errors, requiring that sample sizes used to generate GWAS summary statistics are large [31, 20]. Figure 1B depicts MR-LDP as a probabilistic graphical model, where the observed variables of our model include GWAS summary statistics from both the SNP-exposure and the SNP-outcome, and an external reference panel for genotype data. We assume that *α*_*k*_ and *γ*_*k*_ follow two independent Gaussian distributions. The latent variable *γ*_*k*_ and parameter *β*_0_ jointly assist with formulating the distribution for SNP-outcome. Then, we can formalize the hypothesis testing for *β*_0_ as shown in Figure 1B. The scatter plots of estimated effect sizes for SNP-exposure against SNP-outcome together with the MR-LDP analysis results (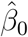 and *p*-value) are shown in Figure 1C. In both BMI-T2D and BMI-VV, there is a dominant proportion of instrumental variants in the center that is largely due to LD, and methods that do not account for LD tend to inflate findings.

**Figure 1:**
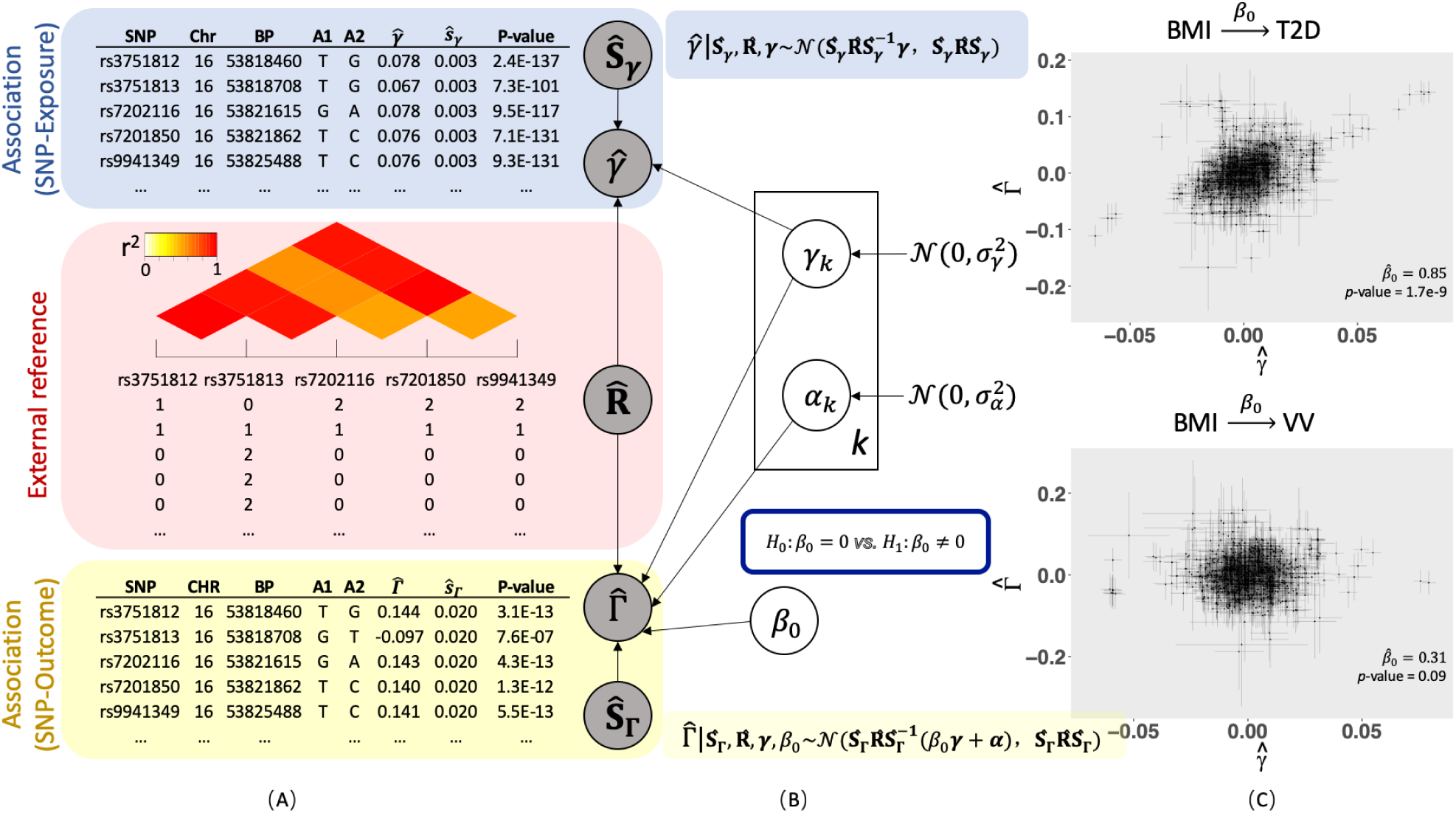
MR-LDP model overview: (A) Inputs for MR-LDP include GWAS summary statistics from both the risk factor (Blue) and the disease outcome (Yellow), and an external reference panel data (Red). (B) A probabilistic graphical model representation of MR-LDP. The box is the “plate” representing SNPs, *k* = 1, *…*, *p*. The circles are either variables or parameters. The circles at the root are parameters. The variables in shaded circles are observed (i.e., GWAS summary statistics 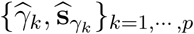 and 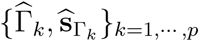, and the estimated 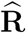 for these *p* SNPs from a reference panel) and variables in unshaded circles are latent variables (i.e., *γ*_*k*_ and *α*_*k*_, *k* = 1, *…*, *p*). The primary goal is to conduct a formal hypothesis testing for ℋ_0_: *β*_0_ = 0 vs ℋ_1_: *β*_0_ ≠ 0. (C) Scatter plots of effect sizes with their standard errors for two exposure-outcome pairs: BMI-T2D and BMI-VV; T2D for type-2 diabetes and VV for varicose veins. Dots represent the effect sizes from SNP-exposure against these from SNP-outcome, and horizontal and vertical bars represent the standard errors from SNP-exposure and SNP-outcome, respectively. The estimated *β*_0_ and its *p*-value from MR-LDP are not shown in each subfigure.

### 2.5 Details of MR-LDP

#### Parameterization for causal relationship

The relationship between ***γ*** and **Γ** can be constructed using linear structural models as follows:

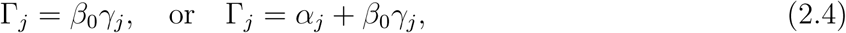

where *j* = 1, *…*, *p*, considering without/with horizontal pleiotropy, respectively [12, 32]. Note that *β*_0_ is the effect size of the exposure on the outcome and ***α*** = [*α*_1_, *…*, *α*_*p*_]^*T*^ is the vector of effects of genetic variants on the outcome due to horizontal pleiotropy. Importantly, *β*_0_ can be interpreted as the causal effect between exposure and outcome in the study [32]. More details regarding linear structural models incorporating the relationship (2.4) are available in the supplementary document. As MR-LD can be taken as a special case of MR-LDP by taking all ***α*** to be zero, we focus on deriving MR-LDP in the main text and provide the supplementary document for details on MR-LD.

#### Empirical Bayes model

By assuming that ***γ*** and ***α*** are two latent variables coming from two independent Gaussian distributions, the complete-data likelihood can be written as follows:

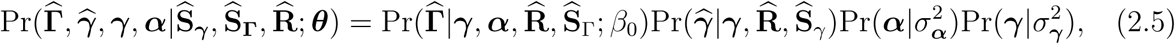

where 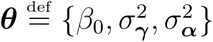 denotes the collection of model parameters. Integrating out the latent variables ***γ*** and ***α***, the marginal likelihood can be written as:

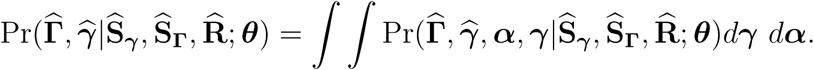

#### Algorithm

The standard expectation-maximization (EM) algorithm is a common choice to find the maximum likelihood for probabilistic models in the presence of latent variables [33]. However, it may cause instability or numerical failure as 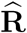 can be non-positive definite due to the relative small sample size in the reference panel. To address these issues, we develop an accelerated variational Bayes (VB) EM algorithm in light of [34], namely, PX-VBEM. Starting with the algorithm, we expand the original MR-LD/MR-LDP model (2.5) as follows:

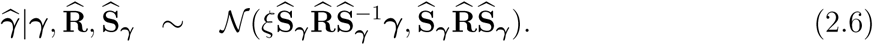

Next, we sketch the VBEM algorithm using the parameter expanded in Equation (2.6) for MR-LDP and algorithmic details for MR-LD can be found in the supplementary document. The model parameters for MR-LDP after parameter expansion become 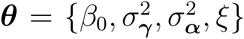. Given variational posterior distribution *q*(***γ, α***), it is straightforward to evaluate the marginal log-likelihood by decomposing it into two parts, the evidence lower bound (ELBO) and the Kullback-Leibler (KL) divergence, which is denoted as follows:

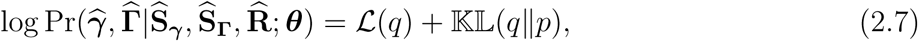

where

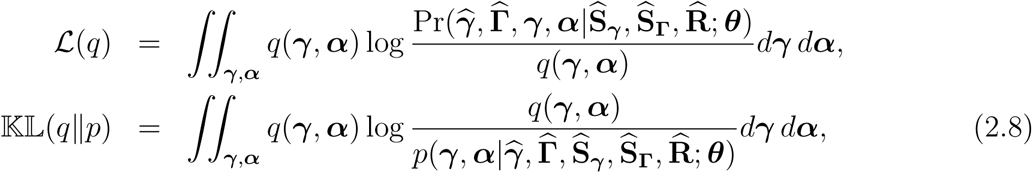

where ℒ(*q*) is the ELBO of the marginal log-likelihood, and 𝕂𝕃 (*q*‖*p*) is the KL divergence between two distributions. Moreover, 𝕂𝕃 (*q*‖*p*) ≥ 0 with equality holding if and only if the variational posterior distribution (*q*) is equal to the true posterior distribution (*p*). As a consequence, minimizing the KL divergence is equivalent to maximizing ELBO. Compared with the standard EM algorithm, the crux of VBEM is to optimize *q* within a factorizable family of distributions by the mean-field assumption [35], which assumes that *q*(***γ, α***) can be factorized as

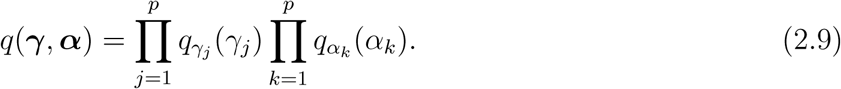

This only assumption in variational inference promotes computational efficiency and scalability in large-scale computational problems given that a coordinate descent algorithm is commonly used to identify the optimal distribution *q*\*. To briefly show this, we first note that this factor-ization (2.9) is used as an approximation for the posterior distribution 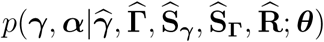. In the VB E-step, given the latent variables ***γ***_*-k*_ and ***α***, the terms with *γ*_*k*_ have a quadratic form, where ***γ***_*-k*_ is the ***γ*** vector removing the *k*-th element. Similarly, when all other latent variables fixed, we can show that the terms with *α*_*k*_ also take a quadratic form. Thus, the variational posterior distribution for *γ*_*k*_ and *α*_*k*_ are both from Gaussian distributions, 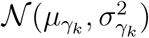 and 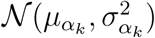, respectively, where we call 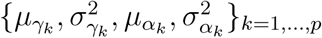 variational parameters. The details of derivations for updating these variational parameters, and the ELBO ℒ(*q*) in the marginal log-likelihood (2.7) at the old parameter ***θ***^*old*^ can be found in the supplementary document. After updating variational parameters in the VB E-step, model parameters (***θ***) can be updated by setting the derivative of the ELBO to zero. Derivation details can be found in supplementary document, where we summarize the PX-VBEM algorithms for MR-LD and MR-LDP in Algorithms 1 and 2, respectively.

#### Inference for causality

We can easily formulate the problem (2.5) as a statistical test for the null hypothesis that the health risk factor is not associated with the disease of interest, or ℋ_0_: *β*_0_ = 0. Testing this hypothesis requires evaluating the marginal log-likelihood of observed data in MR-LD or MR-LDP similar to what has been done previously in [36, 37]; details are given in supplementary document. As VB searches within a factorizable family for the posterior distribution, one can only obtain an approximation for the posterior distribution of latent variables. Earlier works showed that VBEM provides useful and accurate posterior mean estimates [38]. Despite its computational efficiency and accuracy for estimating posterior mean, VB suffers from under-estimating the variance of the target distribution [25, 39, 40]. Thus, the evidence lower bound (ELBO) from VB-type algorithm cannot be directly used to conduct a likelihood-based test. In this paper, we follow Yang et al. [37] and adopt the similar strategy to calibrate ELBO as well as mitigate the bias of variance. Details for the PX-VBEM algorithm and the calibration of ELBO can be found in the supplementary document.

#### Relationship between MR-LD and TWAS

Using transcriptome data as risk factors, MR-LD can be viewed as a TWAS-type analysis using summary-level data from both expression quantitative trait loci (eQTL) and GWAS, where eQTL and GWAS summary statistics are used for SNP-exposure and SNP-outcome, respectively. Since TWAS-type analysis only seeks genes that are significantly associated with the outcome of interest at the genome-wide level, one cannot infer causality without excluding other potential associations, e.g., horizontal pleiotropy. We note that PMR-Egger [41] was recently proposed to calibrate the type-I error control by using a burden test assumption to infer causal relationship. However, this assumption depends heavily on the fact that all effect sizes from horizontal pleiotropy are the same. Therefore, MR-LDP can also be viewed as a relaxation of the burden assumption that is more powerful to account for horizontal pleiotropy with more general patterns.

## 3 Results

### 3.1 Simulations

#### Methods for comparison

We compared the performance of five methods in the main text: (1) our MR-LD and MR-LDP implemented in the R package *MR.LDP*; (2) GSMR implemented in the R package *gsmr*; (3) RAPS implemented in the R package *mr.raps*; (4) IVW implemented in the R package *MendelianRandomization*; (5) MR-Egger implemented in the R package *MendelianRandomization*. All methods were used with default settings. We conducted comprehensive simulation studies to better gauge the performance of each method in simulation studies in terms of type-I error control and point estimates.

In simulation studies, we considered genetic instruments both without and with horizontal pleiotropy. In the scenario that genetic instruments have horizontal pleiotropy, we further considered two cases: sparse and dense horizontal pleiotropy, i.e., sparse horizontal pleiotropy indicates that only a proportion of genetic instruments have direct effects on the outcome while dense horizontal pleiotropy indicates that all genetic instruments have direct effects. As GSMR is a step-wise method that first removes invalid instruments, dense horizontal pleiotropy theoretically implies that all genetic instruments are invalid. To make fair comparisons, we considered sparse horizontal pleiotropy with sparsity at 0.2 or 0.4. In addition, as RAPS, IVW, and MR-Egger tend to inflate type-I error in the presence of LD, we conducted SNP pruning for a fair comparison of point estimates.

#### Simulation settings

To make our simulations as realistic as possible, we started by generating the individual-level two-sample data as follows

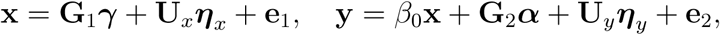

where 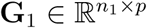 and 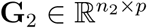 were both genotype matrices, 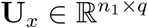 and 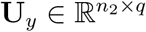 were matrices for confounding variables, *n*_1_ and *n*_2_ were the corresponding sample sizes, *p* was the number of genetic variants, 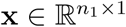 was the exposure vector, 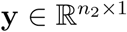 was the outcome vector, and the error terms **e**_1_ and **e**_2_ were obtained from 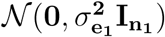 and 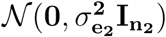, respectively. In this generative model, *β*_0_ was the true causal effect while ***α*** exhibited the direct effects on the disease. We considered two cases: dense and sparse horizontal pleiotropy. For the dense case, we assumed that *α*_*k*_s was independent and identically distributed as 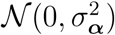. However, for the sparse case, we assumed that only a fraction of *α*_*k*_s was from a Gaussian distribution and remainders were zero. In simulations, we considered sparsity both at 0.2 and 0.4. Note that 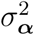 was set by controlling the heritability due to horizontal pleiotropy. Moreover, to mimic the real applications where an external reference panel was applied to estimate the correlation among SNPs, another genotype matrix 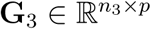 was generated as the reference panel data to estimate the correlation matrix, where *n*_3_ was the sample size in the reference panel. We fixed *n*_1_ = *n*_2_ = 20, 000 but varied *n*_3_ ∈ {500, 2, 500, 4, 000}. In details, we first generated a data matrix from multivariate normal distribution 𝒩(**0, ∑**(*ρ*)), where **∑**(*ρ*) is a block autoregressive (AR) with *ρ* = 0, 0.4, or 0.8 representing weak, moderate or strong LD, respectively. We then generated minor allele frequencies from a uniform distribution 𝕌(0.05, 0.5) and categorized the data matrix into dosage values {0, 1, 2} according to Hardy-Weinberg equilibrium under the generated minor allele frequencies. The number of blocks was *M* = 10 or 20 and the number of SNPs within each block was 50. Correspondingly, *p* = 500 or 1, 000. For confounding variables, we sampled each column of **U**_*x*_ and **U**_*y*_ from a standard normal distribution with fixed *q* = 50 while ***η***_*x*_ ∈ ℝ^*q*×1^ and ***η***_*y*_ ∈ ℝ^*q*×1^ were the corresponding coefficients of confounding factors. Each row of (***η***_*x*_, ***η***_*y*_) was generated from a multivariate normal distribution 𝒩(**0, ∑**_*η*_), and **∑**_*η*_ is a two-by-two matrix with diagonal elements set as 1 and off-diagonal elements set as 0.8.

We then conducted single-variant analysis to obtain the summary statistics for SNP-exposure and SNP-outcome, 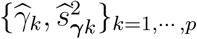 and 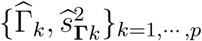, respectively. In simulations, we controlled the signal magnitude for both ***γ*** and ***α*** using their corresponding heritability, 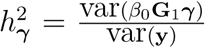 and 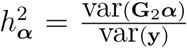, respectively. Thus, we could control 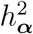 and 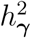 at any value by controlling confounding variables, and the error terms, 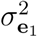 and 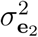. In all settings, we fixed 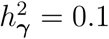 and varied 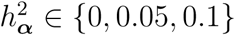.

#### Simulation results: Type-I error control and point estimates

We conducted various simulation studies to make comparisons of MR-LD and MR-LDP with other four commonly used alternative methods: (1) IVW; (2) MR-Egger; (3) GSMR; (4) RAPS. We first compared the type-I error rate for MR-LD and MR-LDP together with other alternative methods based on 1,000 replications. The simulation results for dense pleiotropy and sparse pleiotropy with sparsity at 0.2 and 0.4 are shown in Figures 2, and S2 - S8, respectively with *n*_3_ = 500; 2, 500; 4, 000, respectively. Note that when 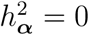, there was no difference between dense and sparse pleiotropy. As shown in the left column of Figure 2A, in the case of no horizontal pleiotropy 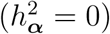, all methods could control type-I error at the nominal level 0.05 generally well when genetic variants were independent (*ρ* = 0). However, as LD become stronger (*ρ* = 0.4 or 0.8), alternative methods failed to control type-I error without SNP pruning. In this setting 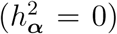, MR-LD and MR-LDP performed equally well in type-I error control. In the presence of horizontal pleiotropy (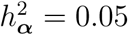 or 0.1), as shown in the middle and right columns of Figure 2A, MR-LD failed to control type-I error for all *ρ* values while type-I error rates of alternative methods without SNP pruning were not controlled in the case of moderate or strong LD. However, MR-LDP could still control type-I error at its nominal level. The similar patterns could be observed for settings under sparse horizontal pleiotropy with sparsity at 0.2 and 0.4 as shown in Figures 2C, and S4 - S8, where the settings was not in favor of MR-LDP. Note that after SNP pruning, genetic variants that remained could be taken as independent. Thus, alternative methods after SNP pruning could control type-I error in all settings. However, this is achieved at the expense of losing weak instruments in LD.

**Figure 2:**
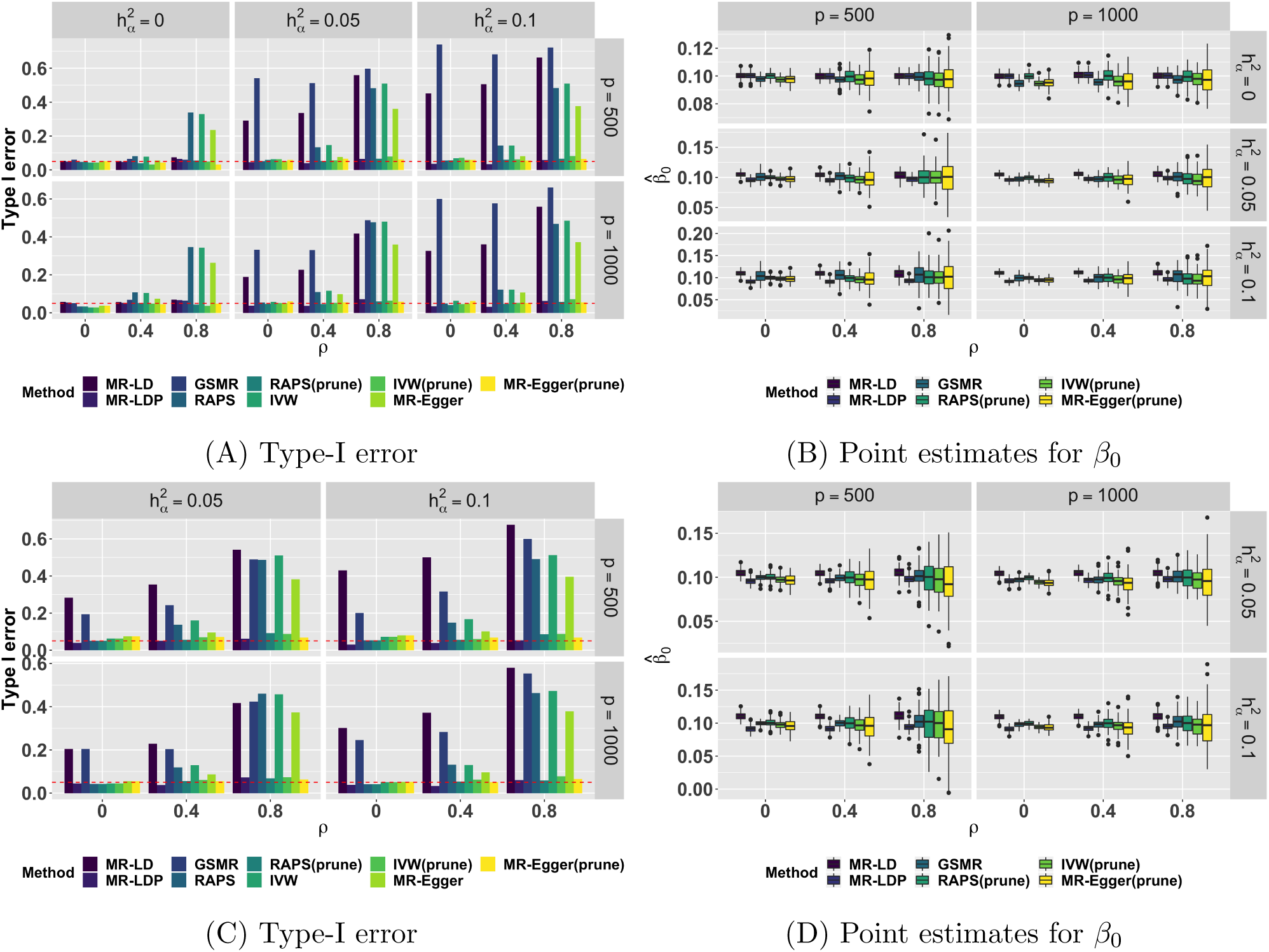
Simulation of type-I error control and point estimates under the dense horizontal pleiotropy (A, B) and the sparse (0.2) horizontal pleiotropy (C, D). *n*_1_ = *n*_2_ = 20, 000, *n*_3_ = 500.

Next, we made comparisons of point estimates for MR-LD and MR-LDP together with alternative methods, where SNP pruning was performed for analysis using alternative methods. In this simulation, *β*_0_ = 0.1 and results were based on 100 replications. Clearly, as shown in Figure 2B, the proposed methods, MR-LD and MR-LDP, had narrower standard errors than alternative methods when LD was moderate or strong (*ρ* = 0.4 or 0.8) as the number of valid instruments were less after SNP pruning for alternative methods. MR-LD and MR-LDP performed equally well in the case of no horizontal pleiotropy, while MR-LD that did not account for horizontal pleiotropy was biased. Similar patterns could be observed for dense and sparse pleiotropy both at sparsity equaling 0.2 and 0.4, as shown in Figures 2D, and S4 - S8.

### 3.2 CAD-CAD and Height-Height studies

In addition, we used real datasets, i.e., CAD-CAD and Height-Height pairs, to compare the estimates from MR-LD and MR-LDP with those from other four alternative methods, where the causal effect *β*_0_ can be taken as known, i.e., *β*_0_ = 1. In these two examples, we used GWAS summary statistics for the same traits (i.e., CAD and BMI, respectively) from three datasets – selection, exposure and outcome [42]. The first two datasets are non-overlapping GWAS for the same trait. The exposure dataset and outcome dataset are non-overlapping individuals from European ancestry. Since IVW, MR-Egger, and RAPS are designed for independent or weak-LD SNPs and GSMR only works for SNPs with moderate LD, we conducted the LD-based clumping to obtain the near-independent SNPs based on PLINK [43]. Individual-level genotype data from UK10K projects was served as the reference panel in this study.

For CAD-CAD analysis, the selection dataset is myocardial infarction (MI) data from UK Biobank (UKB), the exposure data is obtained from the C4D Genetics Consortium [44], and the outcome data is obtained from the transatlantic Coronary ARtery DIsease Genome wide Replication and Meta-analysis (CARDIoGRAM) [45]. We first selected instrumental variants using MI from UKB under different *p*-value thresholds and then conducted MR analysis between the exposure and the outcome using MR-LD, MR-LDP, least squares (LS), IVW, MR-Egger, Raps and GSMR. First, the scatter plots of 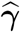 (C4D) against 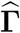 (CAD1) are shown in Figure S9 in the supplementary document, where we found that when a large threshold, e.g., *p*-value=0.001, is applied to select more genetic variants, the points in the center make the inference for causality difficult. We reported the point estimates with its 95% corresponding confidence intervals for all methods in Figures 3 and S10 for *λ* = 0.1 and 0.15, respectively. Clearly, MR-LD and MR-LDP were superior to other methods in terms of smaller bias and shorter confidence interval when the number of instrumental variants is large. Moreover, the estimates from MR-LD and MR-LDP also exhibited statistical significance consistently, while the coverage of *β*_0_ = 1 from other methods was incorrect under small thresholds except for RAPS with larger standard errors due to the SNP pruning. Additionally, estimates from GSMR, IVW, and MR-Egger were always biased when the threshold was small.

**Figure 3:**
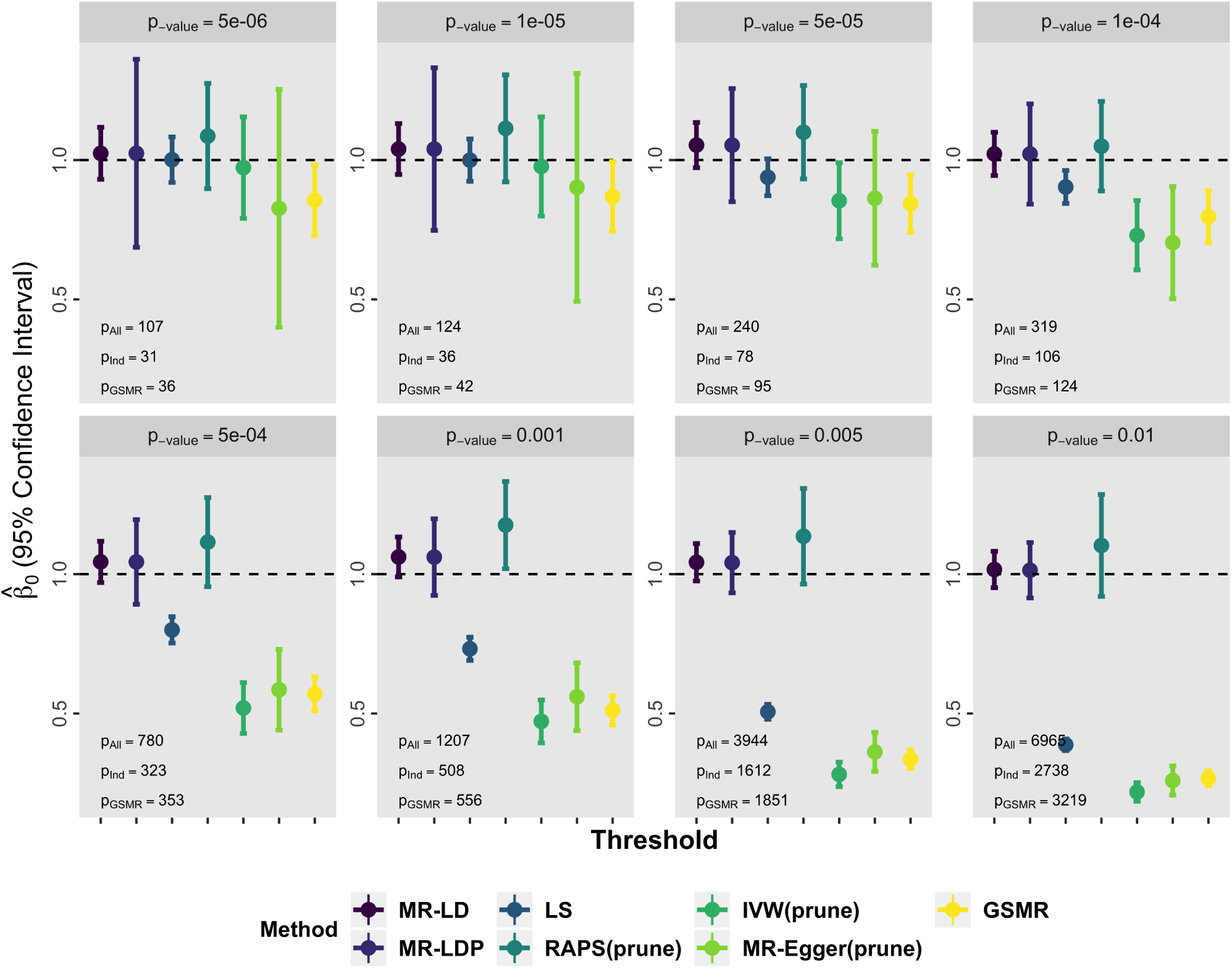
The result of estimates and confidence intervals for CAD-CAD using UK10K as the reference panel with shrinkage parameter *λ* = 0.1 under different *p*-value thresholds to choose genetic variants. MR-LD, MR-LDP and LS methods use all SNPs selected by the screening dataset. Default value is used to choose *r*^2^ in GSMR and the other three methods is 0.001.

Next, we investigated the case that both the exposure and outcome were from human height. In particular, we treated the height in UK Biobank [46] as the screening dataset. The exposure data is from the height for males in a European population-based study and the outcome data is from the height for females in EUR population [47]. First, the scatter plot of 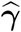 (height for males) against 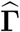 (height for females) are shown in Figure S11 in the supplementary document. Since height is highly polygenic and sample size is very large in [47] (270,000 individuals), the points are crowded in the middle even with a very small threshold (*p*-value = 5 × 10^*-*6^). The results of point estimates with their 95% confidence intervals were illustrated in Figure S12 - S13 for *λ* = 0.1 and 0.15, respectively. Similar patterns were observed in all cases. In particular, RAPS only offered a better performance with larger instrumental variants but did not work for some small thresholds, GSMR failed to estimate the causal effect for this validation study, and other methods underestimated the causal effect with relatively larger standard errors. MR-LD and MR-LDP used all SNPs passing a certain thresholding value and thus provided more accurate estimates of *β*_0_ = 1.

### 3.3 The causal effects of lipids and BMI on common human diseases

We further applied our method, MR-LDP, to estimate the causal effects of lipids and BMI on complex diseases including coronary artery disease (CAD1 and CAD2 from CARDIoGRAM and UKB, respectively), asthma, allergic rhinitis (AR), cancer, major depression disorder (MDD), type 2 diabetes (T2D), dyslipidemia (Dyslid), hypertensive disease (Hyper), hemorrhoids, hernia abdominopelvic cavity hernia, insomnia, iron deficiency anemias (IDA), irritable bowel syndrome (IBS), macular degeneration, osteoarthritis, osteoporosis, peripheral vascular disease (PVD), peptic ulcer (PU), psychiatric disorder, acute reaction to stress (Stress), varicose veins (VV), and disease count (DC). The summary statistics for risk factors include lipoprotein cholesterol(HDL-C), low-density lipoprotein cholesterol (LDL-C), total cholesterol (TC), and body mass index (BMI). Tables S6 and S7 in the supplementary document summarize the total number of SNPs and sample sizes for each trait in each health risk factor or disease outcome and the details for the sources of these GWAS summary statistics.

First, we applied MR-LDP together with alternative methods to analyze to the exposure-outcome pairs using lipids as the exposure, i.e., HDL-C, LDL-C, and TC. Specifically, the selection and exposure datasets were obtained from [48] and [49], respectively, where the threshold for selecting instrumental variants in the selection dataset is set to 1 × 10^*-*4^. The association results from the analysis are summarized in Table 1. Note that we did SNP pruning for RAPS, IVW, and MR-Egger and used the default settings in all alternative methods. As GSMR removes SNPs by providing an LD threshold, we chose to use *r*^2^ = 0.05 as suggested by its paper [15].

**Table 1:** Causal associations of lipids with common diseases using UK10K as the reference panel with shrinkage parameter *λ* = 0.1. MR-LDP uses all SNPs selected by the screening dataset. The thresholds of *r*^2^ for GSMR and the other three methods are 0.05 and 0.001, respectively. Statistically significant results are indicated in blue.

| Lipids | Outcome | #$SNP_{ALL}$ | MR-LDP | #$SNP_{GSMR}$ | GSMR(prune) | #$SNP_{LD}$ | RAPS(prune) | IVW(prune) | MR-Egger(prune) |
| --- | --- | --- | --- | --- | --- | --- | --- | --- | --- |
| HDL-C | CAD1 | 2104 | -0.09(0.027) | 269 | -0.26(0.038) | 203 | -0.38(0.07) | -0.36(0.07) | -0.28(0.157) |
|  | CAD2 | 2071 | -0.08(0.02) | 277 | -0.07(0.03) | 206 | -0.15(0.047) | -0.15(0.047) | -0.08(0.098) |
|  | T2D | 2071 | -0.09(0.031) | 272 | -0.16(0.044) | 206 | -0.33(0.081) | -0.35(0.082) | 0.03(0.17) |
|  | Dyslid | 2071 | -0.14(0.023) | 255 | -0.1(0.03) | 206 | -0.23(0.08) | -0.26(0.076) | -0.17(0.158) |
|  | Hyper | 2071 | -0.05(0.017) | 270 | -0.14(0.022) | 206 | -0.2(0.037) | -0.21(0.038) | -0.09(0.079) |
|  | PVD | 2071 | -0.11(0.048) | 277 | -0.12(0.077) | 206 | -0.19(0.109) | -0.19(0.105) | 0.12(0.222) |
|  | DC | 2071 | -0.04(0.01) | 270 | -0.08(0.013) | 206 | -0.09(0.025) | -0.1(0.025) | -0.03(0.052) |
| LDL-C | CAD1 | 1867 | 0.27(0.029) | 257 | 0.42(0.037) | 193 | 0.34(0.065) | 0.32(0.062) | 0.33(0.133) |
|  | CAD2 | 1820 | 0.11(0.021) | 266 | 0.16(0.027) | 199 | 0.15(0.043) | 0.14(0.043) | 0.26(0.085) |
|  | Dyslid | 1820 | 0.56(0.03) | 258 | 0.94(0.027) | 199 | 0.9(0.053) | 0.86(0.051) | 0.93(0.1) |
|  | DC | 1820 | 0.08(0.01) | 267 | 0.13(0.012) | 199 | 0.13(0.019) | 0.13(0.019) | 0.17(0.037) |
| TC | CAD1 | 2546 | 0.24(0.028) | 309 | 0.46(0.036) | 215 | 0.41(0.061) | 0.39(0.062) | 0.35(0.146) |
|  | CAD2 | 2484 | 0.08(0.02) | 314 | 0.16(0.029) | 218 | 0.15(0.043) | 0.14(0.043) | 0.22(0.094) |
|  | Dyslid | 2484 | 0.54(0.03) | 303 | 1.08(0.029) | 218 | 0.93(0.055) | 0.9(0.051) | 0.97(0.111) |
|  | DC | 2484 | 0.06(0.01) | 314 | 0.13(0.012) | 218 | 0.12(0.019) | 0.12(0.019) | 0.14(0.041) |

In practice, HDL-C and LDL-C are often referred as “good” and “bad” cholesterol, respectively. HDL-C is known to be inversely correlated with heart and vascular diseases. We found several significant protective effects of HDL-C against CAD1 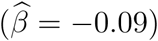, CAD2 disease 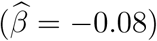, T2D 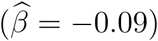, Dyslid 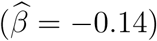, Hyper 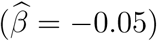, PVD 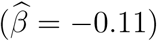 and DC 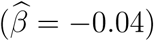, which is consistent with known epidemiological associations in the same direction [50, 51, 52]. In particular, although HDL-C was found to be associated with CAD in multiple observational studies [53, 54, 55], the role of HDL-C in CAD was overturned by later studies [56, 57]. Recently, Zhao et al. [42] showed that the effect of HDL-C in CAD is heterogeneous using different instruments. Moreover, MR-LDP identified the significant negative causality between HDL-C and PVD, which is consistent with previous studies [58, 59]. On the other hand, MR-LDA identified the significant positive causality between LDL-C and CAD which is consistent with the fact that LDL-C narrows the arteries and increases the chance of developing heart diseases. Regarding TC, MR-LDP identified the significant risk effects for cardiovascular disease as confirmed by RCTs.

To better understand of the impact of different thresholds, we re-performed the analysis for HDL-C on CAD1, CAD2, and PVD, separately, using a sequence of thresholds as shown in Figures 4, and S14 - S18. Several patterns can be observed: 1. Methods taking into account LD have small standard errors; 2. Using more SNPs under larger thresholds, the standard errors become smaller; 3. As thresholds become relatively large, e.g., 0.005, the point estimates tend to be biased. The first two patterns are expected. Generally, MR-LDP is robust under different thresholds but shows biasedness when the threshold is too liberal, which is primarily due to the inclusion of invalid variants. As the threshold is relatively large, more genetic variants with no associations to the exposure are included in the analysis, which induce biasedness either upward or downward depending on the directions of effects for invalid instrumental variants.

**Figure 4:**
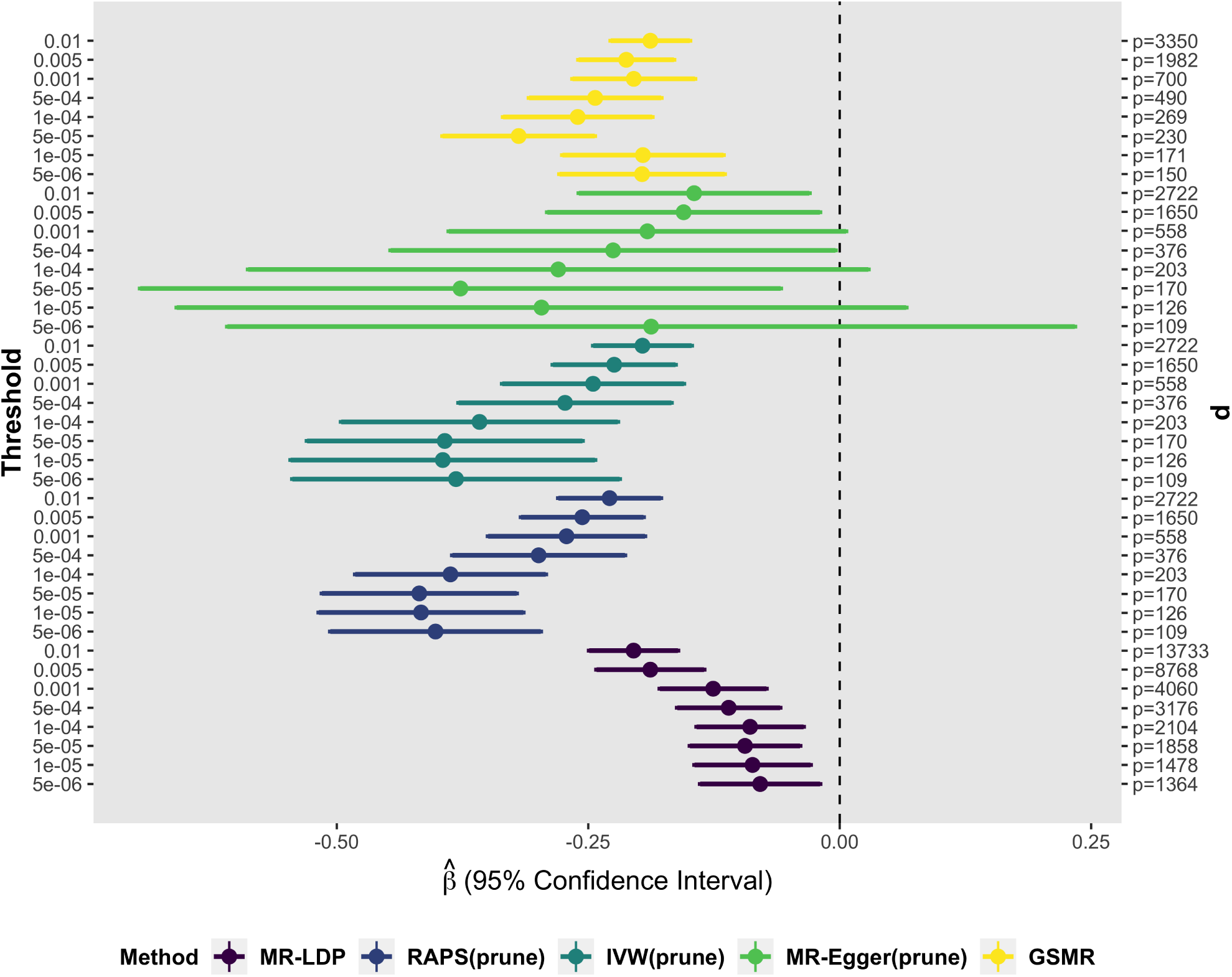
The causal associations of HDL-C on CAD1 under different thresholds using UK10K as the reference panel with *λ* = 0.1.

Second, we examine the associations between BMI and common diseases where the exposure and the selection datasets were obtained from GIANT [60] and [61], respectively. We chose threshold to be 1 × 10^*-*4^ for selecting the instrumental variants from the selection dataset. The association results from the analysis are summarized in Table 2. Overall, our MR-LDP detected relatively more significant causality between BMI and complex diseases in this study. The extent of obesity increase the risk of certain diseases, such as heart disease, type 2 diabetes and hypertensive disease identified by RCT [62].

**Table 2:** Causal associations of BMI with common diseases using UK10K as the reference panel with shrinkage parameter *λ* = 0.1. MR-LDP uses all SNPs selected by the screening dataset. The thresholds of *r*^2^ for GSMR and the other three methods are 0.05 and 0.001, respectively. Statistically significant results are indicated in blue.

| Outcome | #$SNP_{ALL}$ | MR-LDP | #$SNP_{GSMR}$ | GSMR(prune) | #$SNP_{LD}$ | RAPS(prune) | IVW(prune) | MR-Egger(prune) |
| --- | --- | --- | --- | --- | --- | --- | --- | --- |
| CAD1 | 4405 | 0.2(0.084) | 701 | 0.33(0.07) | 563 | 0.2(0.121) | 0.17(0.091) | 0.2(0.129) |
| Asthma | 4428 | 0.28(0.073) | 707 | 0.23(0.061) | 563 | 0.24(0.107) | 0.19(0.08) | 0.18(0.115) |
| CAD2 | 4428 | 0.23(0.066) | 708 | 0.21(0.062) | 563 | 0.26(0.105) | 0.2(0.079) | 0.22(0.113) |
| T2D | 4428 | 0.85(0.141) | 708 | 0.84(0.091) | 563 | 1.22(0.16) | 0.93(0.124) | 1.46(0.175) |
| Dyslip | 4428 | 0.22(0.076) | 704 | 0.29(0.059) | 563 | 0.18(0.133) | 0.16(0.086) | 0.29(0.124) |
| Hemorrhoids | 4428 | 0.3(0.135) | 709 | 0.2(0.111) | 563 | 0.15(0.17) | 0.11(0.129) | -0.1(0.184) |
| Hyper | 4428 | 0.47(0.066) | 703 | 0.5(0.047) | 563 | 0.58(0.095) | 0.46(0.067) | 0.54(0.097) |
| Insomnia | 4428 | 0.77(0.235) | 708 | 0.85(0.215) | 563 | 1.24(0.325) | 0.96(0.246) | 0.6(0.353) |
| Osteoa | 4428 | 0.27(0.078) | 709 | 0.27(0.068) | 563 | 0.26(0.114) | 0.2(0.084) | 0.39(0.119) |
| Osteop | 4428 | -0.44(0.178) | 709 | -0.36(0.15) | 563 | -0.62(0.238) | -0.48(0.178) | -0.73(0.254) |
| PVD | 4428 | 0.35(0.167) | 709 | 0.41(0.159) | 563 | 0.32(0.242) | 0.24(0.183) | 0.41(0.263) |
| DC | 4428 | 0.27(0.035) | 700 | 0.3(0.027) | 563 | 0.3(0.051) | 0.23(0.037) | 0.26(0.053) |

We also estimated some causal effects that are rarely involved in the previous MR analysis but reported in the epidemiological studies. For instance, BMI is an important risk factor for hemorrhoids [63]. BMI is positively associated with knee osteoarthritis and sleep duration reported by [64] and [65], respectively. We also confirmed a protective effect of BMI on osteoporosis reported by [66] and [67]. Moreover, the increased BMI is also considered to be one of the contributing factors for peripheral vascular disease [68].

In addition, MR-Egger is too conservative to identify the causal relationship between BMI and common diseases, and the same conclusion can be found in [18]. Similar to lipids studies, we re-performed the analysis for BMI on hemorrhoids and PVD, respectively, using a sequence of thresholds as shown in Figures S19 - S22. The patterns are similar to those in Figures 4, and S14 - S18

## 4 Discussion

Here, we proposed a statistically rigorous and efficient approach to perform a two-sample MR analysis that accounts for both LD structure and horizontal pleiotropy using GWAS summary statistics and a genotype reference panel. We implemented our method in the R package *MR.LDP*, which is available for download at Github. MR-LDP jointly estimated the causal effect through an approximated likelihood of GWAS summary statistics from both the risk factor and disease outcome using an additional variance component to eliminate the impact of horizontal pleiotropy. Thus, MR-LDP controls for type-I error in the presence of LD structure among instrumental variants and horizontal pleiotropy and is statistically more powerful in identifying causal effects.

MR-LDP is particularly suited to analyze complex traits that have multiple instrumental variants within LD. The key is to jointly model the distributions for summary statistics and the causal relationship between the risk factor and disease outcome. The only approximation here uses the fact of polygenicity in complex traits for the distributions of summary statistics. Moreover, we model the causality as Equation (2.4) as the average of ‘local’ causal effect [32]. The linear model (2.4) holds in very general situations, beyond the linear structural model presented in the supplementary document; see Appendix A in [32] for details. To further eliminate the impact of horizontal pleiotropy, we used a random effect to control the variation in disease outcome. As horizontal pleiotropy is not an estimate of interest, a Gaussian distribution with a mean of zero and a variance parameter is generally robust although the underlying horizontal pleiotropy is sparse. Therefore, the complete-data likelihood for MR-LDP can be written as Equation (2.5). As the iteration for the standard EM algorithm involves inverting 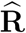, which may cause numerical failure, we developed a PX-VBEM algorithm by expanding parameters. Our previous works have shown that the parameter expansion step is crucial in speeding up the algorithm, and we refer to the supplementary document in [36] for details. To further conduct hypothesis testing for causal effects, we calibrated the EBLO from the PX-VBEM algorithm. In our numerical studies, we demonstrated that MR-LDP is more effective in controlling type-I error in the presence of LD and either sparse or dense horizontal pleiotropy. These merits enable us to apply MR-LDP using GWAS summary statistics, likely yielding more fruitful and meaningful causal discovery in the future.

We used two pairs (CAD-CAD and Height-Height) of real data to partially validate the proposed method. As the risk factor and the outcome are the same, we can take true causal effect as known (*β*_0_ = 1). By applying MR-LD and MR-LDP with alternative methods, we found that estimates from the proposed methods can effectively cover the true *β*_0_ with 95% confidence intervals with instrument variants chosen under a wide ranging of thresholds. When more instrumental variants come into the model under a less stringent threshold, the estimates for the causality have narrower confidence intervals or smaller standard errors. We also note that MR-LDP has wider confidence interval. This is because MR-LDP makes additional efforts to model the horizontal pleiotropy.

In this article, we primarily focus on modeling the lipids and BMI as the exposures and complex diseases as the outcomes. Using a threshold of 1 × 10^*-*4^ in the selection dataset, we identified multiple pairs of significant causal relationships, including a protective effect of high-density lipoprotein cholesterol (HDL-C) on peripheral vascular disease (PVD), and a positive causal effect of body mass index (BMI) on hemorrhoids. We further demonstrated the robustness of MR-LDP using a sequence of threshold values to select instrumental variants. The empirical results show that the threshold of 0.001 is optimal to balance the standard error and biasedness. However, MR-LDP is not without limitations. First, MR-LDP cannot be utilized for overlapped samples in SNP-exposure and SNP-outcome. Furthermore, MR-LDP cannot address the selection bias explicitly but uses an extra SNP-exposure summary statistics to select instrumental variants.

## Web Resources

*MR.LDP* is available at Github (https://github.com/QingCheng0218/MR.LDP). BMI(Jap): ftp://ftp.ebi.ac.uk/pub/databases/gwas/summary_statistics/AkiyamaM_28892062_GCST004904. Other BMI datasets: https://portals.broadinstitute.org/collaboration/giant/index.php/GIANT_consortium_data_files#2018_GIANT_and_UK_BioBank_Meta_Analysis_for_Public_Release lipids(screen datasets): http://csg.sph.umich.edu/willer/public/lipids2010/. lipids(exposure datasets): http://csg.sph.umich.edu/willer/public/lipids2013/. CAD datasets: http://www.cardiogramplusc4d.org/data-downloads/ Common human disease datasets: http://cnsgenomics.com/data.html. UK10K datasets: https://www.uk10k.org/data_access.html.

## Acknowledgements

This work was supported by grant R-913-200-098-263 from the Duke-NUS Medical School, AcRF Tier 2 (MOE2016-T2-2-029, MOE2018-T2-1-046 and MOE2018-T2-2-006) from the Ministry of Education, Singapore, grant No. 71501089, No. 11501579 and No. 71472023 from the National Natural Science Foundation of China; and grant Nos. 22302815, No. 12316116 and No. 12301417 from the Hong Kong Research Grant Council. The computational work for this article was partially performed using resources from the National Supercomputing Centre, Singapore (https://www.nscc.sg).

