## supplementary document for "MR-LDP: a two-sample Mendelian randomization for GWAS summary statistics accounting for linkage disequilibrium and horizontal pleiotropy"

**5 The detail information of GWAS Datasets** **39**

### 1 Statistical Model for MR-LD and MR-LDP

#### 1.1 Validity of instrumental variables

Denote $\mathbf{G} \in \mathbb{R}^{n\times p}$ the matrix for $p$ SNPs among $n$ independent individuals, $\mathbf{x} \in \mathbb{R}^{n\times 1}$ and $\mathbf{y} \in \mathbb{R}^{n\times 1}$ the corresponding exposure and outcome variable, respectively, and $\mathbf{U} \in \mathbb{R}^{n\times q}$ the matrix for $q$ confounding factors among $n$ samples. The SNP-exposure and SNP-outcome true effects are denoted as $\boldsymbol{\gamma} \in \mathbb{R}^{p\times 1}$ and $\boldsymbol{\Gamma} \in \mathbb{R}^{p\times 1}$, respectively, for all $p$ SNPs. Then, we can depict the causal model through a directed acyclic graph in Figure S1, where the dashed line represents the horizontal pleiotropy of genetic variants on outcome, denoted as $\boldsymbol{\alpha} \in \mathbb{R}^{p\times 1}$. We start from the classical assumption that all IVs are valid (namely, core assumptions include: all $\boldsymbol{\gamma}$s are non-zero, SNPs are independent of the confounding factors, and no horizontal pleiotropy holds), where the associations of the exposure and the outcome can be framed as the following linear structural model [1]:

$$\mathbf{x} = \sum_{j=1}^{p} \mathbf{g}_j \gamma_j + \mathbf{U}\boldsymbol{\eta}_x + \boldsymbol{\epsilon}_x, \quad \mathbf{y} = \beta_0 \mathbf{x} + \mathbf{U}\boldsymbol{\eta}_y + \boldsymbol{\epsilon}_y, \tag{S1}$$

where $\beta_0$ is the effect size of the exposure on the outcome, $\mathbf{g}_j$ is the $j$-th column of $\mathbf{G}$, $\boldsymbol{\eta}_x$ and $\boldsymbol{\eta}_y$ are effects on confounding factors for exposure and outcome, respectively, and $\boldsymbol{\epsilon}_x$ and $\boldsymbol{\epsilon}_y$ are independent random noises. Importantly, in this model (S1, $\beta_0$ can be interpreted as the causal effect of the exposure on the outcome as long as the core assumptions for IV are satisfied [2, 3, 1]. Clearly, the effect size of genetic variant $\mathbf{g}_j$ on the outcome $\mathbf{y}$ can be explicitly expressed as $\Gamma_j = \beta_0 \gamma_j$. Thus $\Gamma_j$ is linear to $\gamma_j$, and $\beta_0$ can be interpreted as the causal effect between exposure and outcome in the study [4]. In practice, horizontal pleiotropy is abundant in complex traits and violation of such an assumption can induce severe bias in MR analysis. To investigate the impact of horizontal pleiotropy, one may consider a modified linear structural model [4]:

$$\mathbf{x} = \sum_{j=1}^{p} \mathbf{g}_j \gamma_j + \mathbf{U}\boldsymbol{\eta}_x + \epsilon_x, \quad \mathbf{y} = \sum_{j=1}^{p} \mathbf{g}_j \alpha_j + \beta_0 \mathbf{x} + \mathbf{U}\boldsymbol{\eta}_y + \epsilon_y, \tag{S2}$$

where genetic variants have direct effects on the outcome, denoted as $\boldsymbol{\alpha} = [\alpha_1, \ldots, \alpha_p]^T$. Implicitly, model (S2) also assumes a linear relationship between $\gamma_j$ and $\Gamma_j$ but with a intercept deviating from origin: $\Gamma_j = \alpha_j + \beta_0 \gamma_j$, which is essential to model causality in the presence of horizontal pleiotropy. The linear structural models (S1) and (S2) are not useful in practice as confounding factors are usually not observed in observational studies. However, these two models shed a fundamental insight into modeling causality using summary statistics, namely two-sample MR analysis.

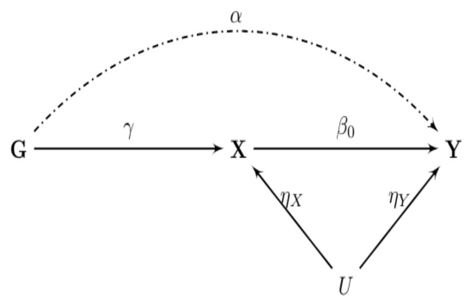

Figure S1: Causal diagram representing three IV assumptions.

#### 1.2 Model for MR-LD

Suppose that all genetic variants are satisfied with the core assumptions for an IV model. Invoking by [2, 4], the linear structural model (S1) suggests that we can model the causal effect in the absence of pleiotropy as

$$\Gamma_k = \beta_0 \gamma_k, \quad \text{for } k = 1, \cdots, p, \tag{S3}$$

where $\beta_0$ is the causal effect of interest. We assign a Gaussian prior on each $\gamma_k$, that is $\boldsymbol{\gamma} \sim \mathcal{N}(\mathbf{0}, \sigma_{\gamma}^{\mathbf{2}}\mathbf{I_p})$. The Gaussian prior is widely used in genetics studies due to polygenicity [5, 6, 7], which offers a great computational advantage over the complicated ones. Combining Equations (X) and (S3), the likelihood for summary statistics from SNP-outcome can be writen as

$$\widehat{\boldsymbol{\Gamma}}|\boldsymbol{\Gamma}, \widehat{\mathbf{R}}, \widehat{\mathbf{S}}_{\Gamma} \sim \mathcal{N}(\widehat{\mathbf{S}}_{\Gamma}\widehat{\mathbf{R}}\widehat{\mathbf{S}}_{\Gamma}^{-1}\beta_0\boldsymbol{\gamma}, \widehat{\mathbf{S}}_{\Gamma}\widehat{\mathbf{R}}\widehat{\mathbf{S}}_{\Gamma}). \tag{S4}$$

Taking $\boldsymbol{\gamma}$ as the latent variables, the completely-data likelihood can be written as follows

$$\Pr(\widehat{\boldsymbol{\Gamma}}, \widehat{\boldsymbol{\gamma}}, \boldsymbol{\gamma}|\widehat{\mathbf{S}}_{\gamma}, \widehat{\mathbf{S}}_{\boldsymbol{\Gamma}}, \widehat{\mathbf{R}}; \boldsymbol{\theta}) = \mathcal{N}(\widehat{\mathbf{S}}_{\boldsymbol{\Gamma}}\widehat{\mathbf{R}}\widehat{\mathbf{S}}_{\boldsymbol{\Gamma}}^{-1}\beta_0\boldsymbol{\gamma}, \widehat{\mathbf{S}}_{\boldsymbol{\Gamma}}\widehat{\mathbf{R}}\widehat{\mathbf{S}}_{\boldsymbol{\Gamma}})\ \mathcal{N}(\widehat{\mathbf{S}}_{\gamma}\widehat{\mathbf{R}}\widehat{\mathbf{S}}_{\gamma}^{-1}\boldsymbol{\gamma}, \widehat{\mathbf{S}}_{\gamma}\widehat{\mathbf{R}}\widehat{\mathbf{S}}_{\gamma})\ \mathcal{N}(\mathbf{0}, \sigma_{\gamma}^{\mathbf{2}}\mathbf{I_p}), \tag{S5}$$

where $\boldsymbol{\theta} = \{\beta_0, \sigma_{\boldsymbol{\gamma}}^2\}$ is the collection of model parameters. The marginal likelihood can be obtained through integrating out the latent variable $\boldsymbol{\gamma}$, that is

$$\Pr(\widehat{\boldsymbol{\Gamma}}, \widehat{\boldsymbol{\gamma}}|\widehat{\mathbf{S}}_{\gamma}, \widehat{\mathbf{S}}_{\boldsymbol{\Gamma}}, \widehat{\mathbf{R}}; \boldsymbol{\theta}) = \int_{\boldsymbol{\gamma}} \Pr(\widehat{\boldsymbol{\Gamma}}, \widehat{\boldsymbol{\gamma}}, \boldsymbol{\gamma}|\widehat{\mathbf{S}}_{\gamma}, \widehat{\mathbf{S}}_{\boldsymbol{\Gamma}}, \widehat{\mathbf{R}}; \boldsymbol{\theta}) d\boldsymbol{\gamma}.$$

Note that we further expand MR-LD (S5) as the following equation.

$$\Pr(\widehat{\boldsymbol{\Gamma}}, \widehat{\boldsymbol{\gamma}}, \boldsymbol{\gamma}|\widehat{\mathbf{S}}_{\gamma}, \widehat{\mathbf{S}}_{\boldsymbol{\Gamma}}, \widehat{\mathbf{R}}; \boldsymbol{\theta}) = \mathcal{N}(\widehat{\mathbf{S}}_{\boldsymbol{\Gamma}}\widehat{\mathbf{R}}\widehat{\mathbf{S}}_{\boldsymbol{\Gamma}}^{-1}\beta_0\boldsymbol{\gamma}, \widehat{\mathbf{S}}_{\boldsymbol{\Gamma}}\widehat{\mathbf{R}}\widehat{\mathbf{S}}_{\boldsymbol{\Gamma}})\ \mathcal{N}(\xi\widehat{\mathbf{S}}_{\gamma}\widehat{\mathbf{R}}\widehat{\mathbf{S}}_{\gamma}^{-1}\boldsymbol{\gamma}, \widehat{\mathbf{S}}_{\gamma}\widehat{\mathbf{R}}\widehat{\mathbf{S}}_{\gamma})\ \mathcal{N}(\mathbf{0}, \sigma_{\gamma}^{\mathbf{2}}\mathbf{I_p}),$$

where $\boldsymbol{\theta} \overset{\text{def}}{=} \{\beta_0, \sigma_{\boldsymbol{\gamma}}, \xi\}$ is the collection of model parameters.

### 2 The derivation of PX-VBEM algorithm

In this section, we present the details on deriving PX-VBEM algorithm for both MR-LD and MR-LDP.

#### 2.1 PX-VBEM for MR-LD

##### 2.1.1 Variational E-step:

Let $\boldsymbol{\theta} \stackrel{\text{def}}{=} \{\beta_0, \sigma_{\boldsymbol{\gamma}}, \xi\}$ be the collection of model parameters, the log-likelihood of complete-data can be written as

$$
\begin{aligned}
& \log \Pr(\widehat{\boldsymbol{\Gamma}}, \widehat{\boldsymbol{\gamma}}, \boldsymbol{\gamma} | \widehat{\mathbf{S}}_{\gamma}, \widehat{\mathbf{S}}_{\boldsymbol{\Gamma}}, \widehat{\mathbf{R}}; \boldsymbol{\theta}) \\
= \; & \log \; \mathcal{N}(\widehat{\mathbf{S}}_{\boldsymbol{\Gamma}} \widehat{\mathbf{R}} \widehat{\mathbf{S}}_{\boldsymbol{\Gamma}}^{-1} \beta_0 \boldsymbol{\gamma}, \widehat{\mathbf{S}}_{\boldsymbol{\Gamma}} \widehat{\mathbf{R}} \widehat{\mathbf{S}}_{\boldsymbol{\Gamma}}) N(\xi \widehat{\mathbf{S}}_{\gamma} \widehat{\mathbf{R}} \widehat{\mathbf{S}}_{\gamma}^{-1} \boldsymbol{\gamma}, \widehat{\mathbf{S}}_{\gamma} \widehat{\mathbf{R}} \widehat{\mathbf{S}}_{\gamma}) \; \mathcal{N}(0, \sigma_{\boldsymbol{\gamma}}^2 \mathbf{I}_p) \\
= \; & \frac{p}{2} \log(2\pi) - \frac{1}{2} \log |\widehat{\mathbf{S}}_{\boldsymbol{\Gamma}} \widehat{\mathbf{R}} \widehat{\mathbf{S}}_{\boldsymbol{\Gamma}}| - \frac{1}{2} (\widehat{\boldsymbol{\Gamma}} - \widehat{\mathbf{S}}_{\boldsymbol{\Gamma}} \widehat{\mathbf{R}} \widehat{\mathbf{S}}_{\boldsymbol{\Gamma}}^{-1} \beta_0 \boldsymbol{\gamma})^{\mathrm{T}} (\widehat{\mathbf{S}}_{\boldsymbol{\Gamma}} \widehat{\mathbf{R}} \widehat{\mathbf{S}}_{\boldsymbol{\Gamma}})^{-1} (\widehat{\boldsymbol{\Gamma}} - \widehat{\mathbf{S}}_{\boldsymbol{\Gamma}} \widehat{\mathbf{R}} \widehat{\mathbf{S}}_{\boldsymbol{\Gamma}}^{-1} \beta_0 \boldsymbol{\gamma}) \\
- \; & \frac{p}{2} \log(2\pi) - \frac{1}{2} \log |\widehat{\mathbf{S}}_{\gamma} \widehat{\mathbf{R}} \widehat{\mathbf{S}}_{\gamma}| - \frac{1}{2} (\widehat{\boldsymbol{\gamma}} - \xi \widehat{\mathbf{S}}_{\gamma} \widehat{\mathbf{R}} \widehat{\mathbf{S}}_{\gamma}^{-1} \boldsymbol{\gamma})^{\mathrm{T}} (\widehat{\mathbf{S}}_{\gamma} \widehat{\mathbf{R}} \widehat{\mathbf{S}}_{\gamma})^{-1} (\widehat{\boldsymbol{\gamma}} - \xi \widehat{\mathbf{S}}_{\gamma} \widehat{\mathbf{R}} \widehat{\mathbf{S}}_{\gamma}^{-1} \boldsymbol{\gamma}) \\
- \; & \frac{p}{2} \log(2\pi) - \frac{p}{2} \log \sigma_{\gamma}^2 - \frac{1}{2\sigma_{\gamma}^2} \sum_{k=1}^{p} \gamma_k^2 \qquad \text{(S1)}
\end{aligned}
$$

We could obtain the expression of $\log q(\gamma_j)$ as follows with the help of equation (10.9) in [8]

$$
\begin{aligned}
\log q(\gamma_j) = \; & E_{q_{-j}} [\log \Pr(\widehat{\boldsymbol{\Gamma}}, \widehat{\boldsymbol{\gamma}}, \boldsymbol{\gamma} | \widehat{\mathbf{S}}_{\gamma}, \widehat{\mathbf{S}}_{\boldsymbol{\Gamma}}, \widehat{\mathbf{R}}; \boldsymbol{\theta})] + const(\gamma_k) \\
= \; & E_{q_{-j}} \left[ \left( \frac{\xi}{\widehat{\gamma}_j} \widehat{\mathbf{s}}_{\gamma j}^2 - \frac{\xi^2}{\widehat{\mathbf{s}}_{\gamma j}} \sum_{k \neq j} \frac{\gamma_k \widehat{\mathbf{R}}_{jk}}{\widehat{\mathbf{s}}_{\gamma k}} + \frac{\beta_0 \widehat{\boldsymbol{\Gamma}}_j}{\widehat{\mathbf{s}}_{\boldsymbol{\Gamma} j}^2} - \frac{\beta_0^2}{\widehat{\mathbf{s}}_{\boldsymbol{\Gamma} j}} \sum_{k \neq j} \frac{\widehat{\mathbf{R}}_{jk} \gamma_k}{\widehat{\mathbf{s}}_{\boldsymbol{\Gamma} k}} \right) \gamma_j \right. \\
- \; & \left. \frac{1}{2} \left( \frac{\xi^2 \widehat{\mathbf{R}}_{jj}}{\widehat{\mathbf{s}}_{\gamma j}^2} + \frac{\beta_0^2 \widehat{\mathbf{R}}_{jj}}{\widehat{\mathbf{s}}_{\boldsymbol{\Gamma} j}^2} + \frac{1}{\sigma_{\gamma}^2} \right) \gamma_j^2 \right] + const(\gamma_k),
\end{aligned}
$$

where the notation $E_{q_{-j}}[\cdot]$ denotes an expectation with respect to the $q$ distributions over all variables $\gamma_i$ , for $i \neq j$. Hence the posterior distribution $\log(\gamma_j)$ should be Gaussian $\mathcal{N}(\mu_j, \sigma_j^2)$

with parameters

$$
\begin{aligned}
\frac{1}{\sigma_j^2} &= \frac{\xi^2 \widehat{\mathbf{R}}_{jj}}{\widehat{\mathbf{s}}_{\gamma j}^2} + \frac{\beta_0^2 \widehat{\mathbf{R}}_{jj}}{\widehat{\mathbf{s}}_{\boldsymbol{\Gamma} j}^2} + \frac{1}{\sigma_\gamma^2}, \\
\mu_j &= \sigma_j^2 \left( \frac{\xi \widehat{\gamma}_j}{\widehat{\mathbf{s}}_{\gamma j}^2} - \frac{\xi^2}{\widehat{\mathbf{s}}_{\gamma j}} \sum_{k \neq j} \frac{\langle \gamma_k \rangle \widehat{\mathbf{R}}_{jk}}{\widehat{\mathbf{s}}_{\gamma k}} + \frac{\beta_0 \widehat{\Gamma}_j}{\widehat{\mathbf{s}}_{\boldsymbol{\Gamma} j}^2} - \frac{\beta_0^2}{\widehat{\mathbf{s}}_{\boldsymbol{\Gamma} j}} \sum_{k \neq j} \frac{\widehat{\mathbf{R}}_{jk} \langle \gamma_k \rangle}{\widehat{\mathbf{s}}_{\boldsymbol{\Gamma} k}} \right),
\end{aligned}
\tag{S2}
$$

where $\langle \gamma_k \rangle \stackrel{\text{def}}{=} E_q(\gamma_k)$.

##### 2.1.2 Variational M-step:

As illustrated before, the ELBO of the marginal log-likelihood can be written as follows given the current estimates of all parameters $\boldsymbol{\theta}$.

$$
\mathcal{L}(\boldsymbol{\theta}) = E_{q(\boldsymbol{\gamma})} \left\{ \log \Pr(\widehat{\boldsymbol{\Gamma}}, \widehat{\boldsymbol{\gamma}}, \boldsymbol{\gamma} | \widehat{\mathbf{S}}_\gamma, \widehat{\mathbf{S}}_{\boldsymbol{\Gamma}}, \widehat{\mathbf{R}}; \boldsymbol{\theta}) \right\} - E_{q(\boldsymbol{\gamma})} \left\{ \log q(\boldsymbol{\gamma}) \right\}.
$$

The second term is independent with the parameters $\boldsymbol{\theta}$, the first term is the expectation of log-complete-data likelihood evaluated under the variational distribution, which can be written as

$$
\begin{aligned}
& E_{q(\boldsymbol{\gamma})} \left\{ \log \Pr(\widehat{\boldsymbol{\Gamma}}, \widehat{\boldsymbol{\gamma}}, \boldsymbol{\gamma} | \widehat{\mathbf{S}}_\gamma, \widehat{\mathbf{S}}_{\boldsymbol{\Gamma}}, \widehat{\mathbf{R}}; \boldsymbol{\theta}) \right\} \\
= \; & E_{q(\boldsymbol{\gamma})} \left\{ \left( \xi \widehat{\boldsymbol{\gamma}}^{\mathrm{T}} \widehat{\mathbf{S}}_\gamma^{-2} + \beta_0 \widehat{\Gamma}^{\mathrm{T}} \widehat{\mathbf{S}}_{\boldsymbol{\Gamma}}^{-2} \right) \boldsymbol{\gamma} - \frac{1}{2} \boldsymbol{\gamma}^{\mathrm{T}} \left( \xi^2 \widehat{\mathbf{S}}_\gamma^{-1} \widehat{\mathbf{R}} \widehat{\mathbf{S}}_\gamma^{-1} + \beta_0^2 \widehat{\mathbf{S}}_{\boldsymbol{\Gamma}}^{-1} \widehat{\mathbf{R}} \widehat{\mathbf{S}}_{\boldsymbol{\Gamma}}^{-1} + \sigma_\gamma^{-2} \mathbf{I}_p \right) \boldsymbol{\gamma}. \right\} \\
- \; & \left. \frac{p}{2} \log \sigma_\gamma^2 \right\} + const(\boldsymbol{\theta}) \\
= \; & \left( \xi \widehat{\boldsymbol{\gamma}} \widehat{\mathbf{S}}_\gamma^{-2} + \beta_0 \widehat{\boldsymbol{\Gamma}}^{\mathrm{T}} \widehat{\mathbf{S}}_{\boldsymbol{\Gamma}}^{-2} \right) \boldsymbol{\mu}_\gamma - \frac{1}{2} \boldsymbol{\mu}_\gamma^{\mathrm{T}} \left( \xi^2 \widehat{\mathbf{S}}_\gamma^{-1} \widehat{\mathbf{R}} \widehat{\mathbf{S}}_\gamma^{-1} + \beta_0^2 \widehat{\mathbf{S}}_{\boldsymbol{\Gamma}}^{-1} \widehat{\mathbf{R}} \widehat{\mathbf{S}}_{\boldsymbol{\Gamma}}^{-1} + \sigma_\gamma^{-2} \mathbf{I}_p \right) \boldsymbol{\mu}_\gamma \\
- \; & \frac{1}{2} \xi^2 Tr(\widehat{\mathbf{S}}_\gamma^{-1} \widehat{\mathbf{R}} \widehat{\mathbf{S}}_\gamma^{-1} \boldsymbol{\Sigma}_\gamma) - \frac{1}{2} \beta_0^2 Tr(\widehat{\mathbf{S}}_{\boldsymbol{\Gamma}}^{-1} \widehat{\mathbf{R}} \widehat{\mathbf{S}}_{\boldsymbol{\Gamma}}^{-1} \boldsymbol{\Sigma}_\gamma) - \frac{p}{2} \log \sigma_\gamma^2 - \frac{1}{2\sigma_\gamma^2} Tr(\boldsymbol{\Sigma}_\gamma) + const(\boldsymbol{\theta}).
\end{aligned}
$$

The second equation is due to the fact: if $\mathbf{x} \sim \mathcal{N}(\mathbf{x} | \boldsymbol{\mu}, \boldsymbol{\Sigma})$, then $E(\mathbf{x}^{\mathrm{T}} \mathbf{A} \mathbf{x}) = \boldsymbol{\mu}^{\mathrm{T}} \mathbf{A} \boldsymbol{\mu} + Tr(\mathbf{A} \boldsymbol{\Sigma})$ for any symmetric matrix $\mathbf{A}$.

By setting the derivative of $E_{q(\boldsymbol{\gamma})} \left\{ \log \Pr(\widehat{\boldsymbol{\Gamma}}, \widehat{\boldsymbol{\gamma}}, \boldsymbol{\gamma} | \widehat{\mathbf{S}}_\gamma, \widehat{\mathbf{S}}_{\boldsymbol{\Gamma}}, \widehat{\mathbf{R}}; \boldsymbol{\theta}) \right\}$ to zero, we obtain the new updates for all parameters

$$\begin{aligned}\beta_0 &= \left\{\boldsymbol{\mu}_{\boldsymbol{\gamma}}^{\mathrm{T}}\widehat{\mathbf{S}}_{\boldsymbol{\Gamma}}^{-1}\widehat{\mathbf{R}}\widehat{\mathbf{S}}_{\boldsymbol{\Gamma}}^{-1}\boldsymbol{\mu}_{\boldsymbol{\gamma}} + Tr(\widehat{\mathbf{S}}_{\boldsymbol{\Gamma}}^{-1}\widehat{\mathbf{R}}\widehat{\mathbf{S}}_{\boldsymbol{\Gamma}}^{-1}\boldsymbol{\Sigma}_{\boldsymbol{\gamma}})\right\}^{-1}\left(\widehat{\boldsymbol{\Gamma}}^{\mathrm{T}}\widehat{\mathbf{S}}_{\boldsymbol{\Gamma}}^{-2}\boldsymbol{\mu}_{\boldsymbol{\gamma}}\right)\\ \sigma_{\boldsymbol{\gamma}}^2 &= \left\{\boldsymbol{\mu}_{\boldsymbol{\gamma}}^{\mathrm{T}}\boldsymbol{\mu}_{\boldsymbol{\gamma}} + Tr(\boldsymbol{\Sigma}_{\boldsymbol{\gamma}})\right\}/p\\ \xi &= \left\{\boldsymbol{\mu}_{\boldsymbol{\gamma}}^{\mathrm{T}}\widehat{\mathbf{S}}_{\boldsymbol{\gamma}}^{-1}\widehat{\mathbf{R}}\widehat{\mathbf{S}}_{\boldsymbol{\gamma}}^{-1}\boldsymbol{\mu}_{\boldsymbol{\gamma}} + Tr(\widehat{\mathbf{S}}_{\boldsymbol{\gamma}}^{-1}\widehat{\mathbf{R}}\widehat{\mathbf{S}}_{\boldsymbol{\gamma}}^{-1}\boldsymbol{\Sigma}_{\boldsymbol{\gamma}})\right\}^{-1}\left(\widehat{\boldsymbol{\gamma}}^{\mathrm{T}}\widehat{\mathbf{S}}_{\boldsymbol{\gamma}}^{-2}\boldsymbol{\mu}_{\boldsymbol{\gamma}}\right)\end{aligned} \tag{S3}$$

where $\boldsymbol{\mu}_{\boldsymbol{\gamma}} = (\mu_1, \cdots, \mu_p)^{\mathrm{T}}, \boldsymbol{\Sigma}_{\boldsymbol{\gamma}} = \mathrm{diag}((\sigma_1^2, \cdots, \sigma_p^2)^{\mathrm{T}})$.

Since the entropy of $q(\boldsymbol{\gamma})$ equals $\frac{p}{2}(\log(2\pi)+1) + \frac{1}{2}\sum_{j=1}^{p}\log\sigma_j^2$, the analytic form of ELBO can be obtained as follows

$$\begin{aligned}&\mathcal{L}(q)\\ =\ & \xi\widehat{\boldsymbol{\gamma}}\widehat{\mathbf{S}}_{\boldsymbol{\gamma}}^{-2}\boldsymbol{\mu}_{\boldsymbol{\gamma}} - \frac{\xi^2}{2}\boldsymbol{\mu}_{\boldsymbol{\gamma}}^{\mathrm{T}}\widehat{\mathbf{S}}_{\boldsymbol{\gamma}}^{-1}\widehat{\mathbf{R}}\widehat{\mathbf{S}}_{\boldsymbol{\gamma}}^{-1}\boldsymbol{\mu}_{\boldsymbol{\gamma}} - \frac{\xi^2}{2}Tr(\widehat{\mathbf{S}}_{\boldsymbol{\gamma}}^{-1}\widehat{\mathbf{R}}\widehat{\mathbf{S}}_{\boldsymbol{\gamma}}^{-1}\boldsymbol{\Sigma}_{\boldsymbol{\gamma}}) + \beta_0\widehat{\boldsymbol{\Gamma}}^{\mathrm{T}}\widehat{\mathbf{S}}_{\boldsymbol{\Gamma}}^{-2}\boldsymbol{\mu}_{\boldsymbol{\gamma}} - \frac{1}{2}\beta_0^2\boldsymbol{\mu}_{\boldsymbol{\gamma}}^{\mathrm{T}}\widehat{\mathbf{S}}_{\boldsymbol{\Gamma}}^{-1}\widehat{\mathbf{R}}\widehat{\mathbf{S}}_{\boldsymbol{\Gamma}}^{-1}\boldsymbol{\mu}_{\boldsymbol{\gamma}}\\ -\ & \frac{1}{2}\beta_0^2 Tr(\widehat{\mathbf{S}}_{\boldsymbol{\Gamma}}^{-1}\widehat{\mathbf{R}}\widehat{\mathbf{S}}_{\boldsymbol{\Gamma}}^{-1}\boldsymbol{\Sigma}_{\boldsymbol{\gamma}}) - \frac{p}{2}\log\sigma_{\boldsymbol{\gamma}}^2 - \frac{1}{2\sigma_{\boldsymbol{\gamma}}^2}\boldsymbol{\mu}_{\boldsymbol{\gamma}}^{\mathrm{T}}\boldsymbol{\mu}_{\boldsymbol{\gamma}} - \frac{1}{2\sigma_{\boldsymbol{\gamma}}^2}Tr(\boldsymbol{\Sigma}_{\boldsymbol{\gamma}}) + \frac{1}{2}\sum_{j=1}^{p}\log\sigma_j^2 - p\log(2\pi)\\ +\ & \frac{p}{2} - \frac{1}{2}\left\{\log|\widehat{\mathbf{S}}_{\boldsymbol{\gamma}}\widehat{\mathbf{R}}\widehat{\mathbf{S}}_{\boldsymbol{\gamma}}| + \log|\widehat{\mathbf{S}}_{\boldsymbol{\Gamma}}\widehat{\mathbf{R}}\widehat{\mathbf{S}}_{\boldsymbol{\Gamma}}| + \widehat{\boldsymbol{\gamma}}^{\mathrm{T}}(\widehat{\mathbf{S}}_{\boldsymbol{\gamma}}\widehat{\mathbf{R}}\widehat{\mathbf{S}}_{\boldsymbol{\gamma}})^{-1}\widehat{\boldsymbol{\gamma}} + \widehat{\boldsymbol{\Gamma}}^{\mathrm{T}}(\widehat{\mathbf{S}}_{\boldsymbol{\Gamma}}\widehat{\mathbf{R}}\widehat{\mathbf{S}}_{\boldsymbol{\Gamma}})^{-1}\boldsymbol{\Gamma}\right\}.\end{aligned}$$

The reduction steps: $\boldsymbol{\mu}_{\boldsymbol{\gamma}} = \xi\boldsymbol{\mu}_{\boldsymbol{\gamma}}$, $\beta_0 = \beta_0/\xi$, $\sigma_{\boldsymbol{\gamma}}^2 = \xi^2\sigma_{\boldsymbol{\gamma}}^2$.

The corresponding PX-VBEM can be summarized as Algorithm 1.

---

**Algorithm 1:** PX-VBEM for MR-LD

---

1 *Initialization*: The parameters $(\sigma_{\gamma_j}^2, \sigma_e^2)$ are initialized using linear mixed model. Meanwhile, we set $\beta_0 = 0$, $\mu_{\boldsymbol{\gamma}} = \mathbf{0}$, $\sigma_{\boldsymbol{\gamma}} = 0.01$, $\xi = 1$.

2 **repeat**

3     **E-step:** At the $t$-th iteration, for each $j = 1, \cdots, p$, the posterior distribution $q(\gamma_j|\boldsymbol{\theta}^{(t)})$ follows Gaussian with parameters obtained using formula (S2) given $\xi = \xi^{(t)} = 1$, $\beta_0 = \beta_0^{(t)}$, $\sigma_{\boldsymbol{\gamma}}^2 = (\sigma_{\boldsymbol{\gamma}}^{(t)})^2$.

4     **M-tesp:** Update $\beta_0$, $\sigma_{\boldsymbol{\gamma}}^2$ and $\xi$ as equation(S3).

5     **Reduction-step:** Rescale the parameters $\boldsymbol{\mu}_{\boldsymbol{\gamma}}^{(t+1)} = \xi^{(t+1)}\boldsymbol{\mu}_{\boldsymbol{\gamma}}^{(t+1)}$, $\beta_0^{(t+1)} = \beta_0^{(t+1)}/\xi^{(t+1)}$, $(\sigma_j^{(t+1)})^2 = (\xi^{(t+1)})^2(\sigma_j^{(t+1)})^2$ and reset $\xi^{(t+1)} = 1$.

6 **until** *convergence or maximum iteration reached*;

---

###### 2.1.3 Statistical Inference for MR-LD

As VB searches within a factorizable family for posterior distribution, one can only obtain an approximation for the posterior distribution of latent variables. Earlier works showed that VBEM provides useful and accurate posterior mean estimates [9]. Despite its computational efficiency and accuracy for estimating posterior mean, VB suffers from under-estimating the variance of target distribution Thus, the ELBO from VB-type algorithm cannot be used directly as a proxy to log-likelihood. In this paper, we follow Yang et al. [10] and adopt the similar strategy to calibrate ELBO as well as mitigate the bias of variance. We first set up the following hypothesis test to formally examine the significance between the exposure and the outcome:

$$\mathcal{H}_0 : \beta_0 = 0 \qquad \text{v.s.} \qquad \mathcal{H}_1 : \beta_0 \neq 0.$$

A likelihood ratio test (LRT) statistic is given by

$$\Lambda \;=\; 2(\log \Pr(\widehat{\gamma}, \widehat{\boldsymbol{\Gamma}} | \widehat{\mathbf{S}}_{\boldsymbol{\gamma}}, \widehat{\mathbf{S}}_{\boldsymbol{\Gamma}}, \widehat{\mathbf{R}}; \widehat{\boldsymbol{\theta}}) - \log \Pr(\widehat{\gamma}, \widehat{\boldsymbol{\Gamma}} | \widehat{\mathbf{S}}_{\boldsymbol{\gamma}}, \widehat{\mathbf{S}}_{\boldsymbol{\Gamma}}, \widehat{\mathbf{R}}; \widehat{\boldsymbol{\theta}}_0)), \tag{S4}$$

where $\widehat{\boldsymbol{\theta}}$ and $\widehat{\boldsymbol{\theta}}_0$ are parameters obtained by maximizing the marginal likelihood, under both the null hypothesis and alternative hypothesis, respectively. As the marginal likelihood cannot be obtained but the ELBO with accurate posterior means, we calibrate the ELBO(denoted by $\widetilde{\mathcal{L}}(\boldsymbol{\theta}, \boldsymbol{\mu}_{\boldsymbol{\gamma}})$) as follows

$$\begin{aligned}
\widetilde{\mathcal{L}}(\boldsymbol{\theta}, \boldsymbol{\mu}_{\boldsymbol{\gamma}}) &= E_{q(\boldsymbol{\gamma})} \left\{ \log \Pr(\widehat{\boldsymbol{\Gamma}}, \widehat{\gamma}, \boldsymbol{\gamma} | \widehat{\mathbf{S}}_{\boldsymbol{\gamma}}, \widehat{\mathbf{S}}_{\boldsymbol{\Gamma}}, \widehat{\mathbf{R}}; \boldsymbol{\theta}) - \log q(\boldsymbol{\gamma}) \right\} \\
&= -\frac{1}{2} \boldsymbol{\mu}_{\boldsymbol{\gamma}}^{\mathrm{T}} \left( \xi^2 \widehat{\mathbf{S}}_{\boldsymbol{\gamma}}^{-1} \widehat{\mathbf{R}} \widehat{\mathbf{S}}_{\boldsymbol{\gamma}}^{-1} + \beta_0^2 \widehat{\mathbf{S}}_{\boldsymbol{\Gamma}}^{-1} \widehat{\mathbf{R}} \widehat{\mathbf{S}}_{\boldsymbol{\Gamma}}^{-2} + \sigma_{\gamma}^{-2} \mathbf{I}_p \right) \boldsymbol{\mu}_{\boldsymbol{\gamma}} \\
&+ \left( \xi \widehat{\gamma}^{\mathrm{T}} \widehat{\mathbf{S}}_{\boldsymbol{\gamma}}^{-2} + \beta_0 \widehat{\boldsymbol{\Gamma}}^{\mathrm{T}} \widehat{\mathbf{S}}_{\boldsymbol{\Gamma}}^{-2} \right) \boldsymbol{\mu}_{\boldsymbol{\gamma}} - \frac{p}{2} \log \sigma_{\gamma}^2 + \frac{1}{2} \log |\widetilde{\boldsymbol{\Sigma}}_{\boldsymbol{\gamma}}| - p \log(2\pi) \\
&- \frac{1}{2} \left\{ \log |\widehat{\mathbf{S}}_{\boldsymbol{\gamma}} \widehat{\mathbf{R}} \widehat{\mathbf{S}}_{\boldsymbol{\gamma}}| + \log |\widehat{\mathbf{S}}_{\boldsymbol{\Gamma}} \widehat{\mathbf{R}} \widehat{\mathbf{S}}_{\boldsymbol{\Gamma}}| + \widehat{\gamma}^{\mathrm{T}} (\widehat{\mathbf{S}}_{\boldsymbol{\gamma}} \widehat{\mathbf{R}} \widehat{\mathbf{S}}_{\boldsymbol{\gamma}})^{-1} \widehat{\gamma} + \widehat{\boldsymbol{\Gamma}}^{\mathrm{T}} (\widehat{\mathbf{S}}_{\boldsymbol{\Gamma}} \widehat{\mathbf{R}} \widehat{\mathbf{S}}_{\boldsymbol{\Gamma}})^{-1} \boldsymbol{\Gamma} \right\},
\end{aligned} \tag{S5}$$

where $\boldsymbol{\mu}_{\boldsymbol{\gamma}}$ and $\widetilde{\boldsymbol{\Sigma}}_{\boldsymbol{\gamma}}$ are the posterior mean and posterior variance for latent variable $\boldsymbol{\gamma}$ and the form of $\widetilde{\boldsymbol{\Sigma}}_{\boldsymbol{\gamma}}$ is from EM/PX-EM by plugging the posterior mean estimates and parameter estimates from VBEM/PX-VBEM which can be expressed as

$$\widetilde{\boldsymbol{\Sigma}}_{\gamma} = \left(\xi^2 \widehat{\mathbf{S}}_{\gamma}^{-1}\widehat{\mathbf{R}}\widehat{\mathbf{S}}_{\gamma}^{-1} + \beta_0^2 \widehat{\mathbf{S}}_{\boldsymbol{\Gamma}}^{-1}\widehat{\mathbf{R}}\widehat{\mathbf{S}}_{\boldsymbol{\Gamma}}^{-2} + \sigma_{\gamma}^{-2}\mathbf{I}_p\right)^{-1}.$$

In summary, we recalibrate the ELBO for MR-LD as follows:

1. Obtain parameters together with variational means under both $\mathcal{H}_0$ and $\mathcal{H}_1$.
2. Recalibrate ELBO using formulae (S5).
3. Calculate the test statistic (S4) using the recalibrated ELBO as a proxy to the marginal log-likelihood.

As demonstrated in Yang et al. [10], this procedure can produce the calibrated ELBO that is highly correlated with the marginal log-likelihood obtained from EM algorithm. Moreover, our empirical results in validation studies show that this procedure works well.

#### 2.2 PX-VBEM for MR-LDP

##### 2.2.1 Variational E-step:

The latent variable: $\boldsymbol{\gamma}, \boldsymbol{\alpha}$ and the parameter set $\boldsymbol{\theta} \stackrel{\text{def}}{=} \{\sigma_{\alpha}^2, \sigma_{\gamma}^2, \beta_0, \xi\}$. Therefor, the log-likelihood of complete-data can be written as follows,

$$\begin{aligned}
& \log \Pr(\widehat{\boldsymbol{\Gamma}}, \widehat{\boldsymbol{\gamma}}, \boldsymbol{\alpha}, \boldsymbol{\gamma} | \widehat{\mathbf{S}}_{\gamma}, \widehat{\mathbf{S}}_{\boldsymbol{\Gamma}}, \widehat{\mathbf{R}}; \boldsymbol{\theta}) \\
= & \log \mathcal{N}(\widehat{\boldsymbol{\Gamma}} | \widehat{\mathbf{S}}_{\boldsymbol{\Gamma}}\widehat{\mathbf{R}}\widehat{\mathbf{S}}_{\boldsymbol{\Gamma}}^{-1}(\beta_0\boldsymbol{\gamma} + \boldsymbol{\alpha}), \widehat{\mathbf{S}}_{\boldsymbol{\Gamma}}\widehat{\mathbf{R}}\widehat{\mathbf{S}}_{\boldsymbol{\Gamma}}) N(\widehat{\boldsymbol{\gamma}} | \xi\widehat{\mathbf{S}}_{\gamma}\widehat{\mathbf{R}}\widehat{\mathbf{S}}_{\gamma}^{-1}\boldsymbol{\gamma}, \widehat{\mathbf{S}}_{\gamma}\widehat{\mathbf{R}}\widehat{\mathbf{S}}_{\gamma}) \, \mathcal{N}(\mathbf{0}, \sigma_{\alpha}^{\mathbf{2}}\mathbf{I_p}) \, \mathcal{N}(\mathbf{0}, \sigma_{\gamma}^{\mathbf{2}}\mathbf{I_p}) \\
= & \left(\beta_0 \widehat{\Gamma}^{\mathrm{T}} \widehat{\mathbf{S}}_{\boldsymbol{\Gamma}}^{-2} + \xi \widehat{\boldsymbol{\gamma}} \widehat{\mathbf{S}}_{\gamma}^{-2}\right)\boldsymbol{\gamma} - \frac{1}{2}\boldsymbol{\gamma}^{\mathrm{T}} \left(\beta_0^2 \widehat{\mathbf{S}}_{\boldsymbol{\Gamma}}^{-1}\widehat{\mathbf{R}}\widehat{\mathbf{S}}_{\boldsymbol{\Gamma}}^{-1} + \xi^2 \widehat{\mathbf{S}}_{\gamma}^{-1}\widehat{\mathbf{R}}\widehat{\mathbf{S}}_{\gamma}^{-1} + \sigma_{\gamma}^{-2}\mathbf{I}\right)\boldsymbol{\gamma} + \widehat{\Gamma}\widehat{\mathbf{S}}_{\boldsymbol{\Gamma}}^{-2}\boldsymbol{\alpha} \\
- & \beta_0 \boldsymbol{\alpha}^{\mathrm{T}} \widehat{\mathbf{S}}_{\boldsymbol{\Gamma}}^{-1}\widehat{\mathbf{R}}\widehat{\mathbf{S}}_{\boldsymbol{\Gamma}}^{-1}\boldsymbol{\gamma} - \frac{p}{2}\log\sigma_{\gamma}^2 - \frac{p}{2}\log\sigma_{\boldsymbol{\alpha}}^2 - \frac{1}{2}\boldsymbol{\alpha}^{\mathrm{T}}\left(\widehat{\mathbf{S}}_{\boldsymbol{\Gamma}}^{-1}\widehat{\mathbf{R}}\widehat{\mathbf{S}}_{\boldsymbol{\Gamma}}^{-1} + \sigma_{\boldsymbol{\alpha}}^{-2}\mathbf{I}\right)\boldsymbol{\alpha} - 2p\log(2\pi) \\
- & \frac{1}{2}\left\{\log|\widehat{\mathbf{S}}_{\gamma}\widehat{\mathbf{R}}\widehat{\mathbf{S}}_{\gamma}| + \log|\widehat{\mathbf{S}}_{\boldsymbol{\Gamma}}\widehat{\mathbf{R}}\widehat{\mathbf{S}}_{\boldsymbol{\Gamma}}| + \widehat{\boldsymbol{\gamma}}^{\mathrm{T}}(\widehat{\mathbf{S}}_{\gamma}\widehat{\mathbf{R}}\widehat{\mathbf{S}}_{\gamma})^{-1}\widehat{\boldsymbol{\gamma}} + \widehat{\boldsymbol{\Gamma}}^{\mathrm{T}}(\widehat{\mathbf{S}}_{\boldsymbol{\Gamma}}\widehat{\mathbf{R}}\widehat{\mathbf{S}}_{\boldsymbol{\Gamma}})^{-1}\boldsymbol{\Gamma}\right\}. \qquad \text{(S6)}
\end{aligned}$$

We use mean field theory to approximate the posterior $q(\boldsymbol{\gamma}, \boldsymbol{\alpha}) = q(\gamma_1)\cdots q(\gamma_p)\cdots q(\alpha_1)\cdots q(\alpha_p)$.

The posterior distribution of $\gamma_j \sim \mathcal{N}(\mu_j, \sigma_j^2)$, where

$$
\begin{aligned}
-\frac{1}{2\sigma_j^2} &= -\frac{\beta_0^2}{2}\frac{\widehat{\mathbf{R}}_{jj}}{\widehat{\mathbf{s}}_{\mathbf{\Gamma}j}^2} - \frac{\xi^2\widehat{\mathbf{R}}_{jj}}{2\widehat{\mathbf{s}}_{\gamma j}^2} - \frac{1}{2\sigma_\gamma^2}, \\
\frac{\mu_j}{\sigma_j^2} &= \beta_0\frac{\widehat{\Gamma}_j}{\widehat{\mathbf{s}}_{\mathbf{\Gamma}j}^2} - \frac{\beta_0^2}{\widehat{\mathbf{s}}_{\mathbf{\Gamma}j}}\left(\sum_{k\neq j}\frac{\langle\gamma_k\rangle\widehat{\mathbf{R}}_{jk}}{\widehat{\mathbf{s}}_{\mathbf{\Gamma}k}}\right) - \frac{\beta_0}{\widehat{\mathbf{s}}_{\mathbf{\Gamma}j}}\left(\sum_{i=1}^{p}\frac{\alpha_i\widehat{\mathbf{R}}_{ij}}{\widehat{\mathbf{s}}_{\mathbf{\Gamma}i}}\right) + \frac{\xi\widehat{\gamma}_j}{\widehat{\mathbf{s}}_{\gamma j}^2} - \frac{\xi^2}{\widehat{\mathbf{s}}_{\gamma j}}\left(\sum_{k\neq j}\frac{\langle\gamma_k\rangle\widehat{\mathbf{R}}_{jk}}{\widehat{\mathbf{s}}_{\gamma k}}\right). \quad \text{(S7)}
\end{aligned}
$$

where $\langle\gamma_k\rangle \stackrel{\text{def}}{=} E_q(\gamma_k)$.

Therefore, we can denote the posterior of $\boldsymbol{\gamma}$ by $\mathcal{N}(\boldsymbol{\gamma}|\mu_{\boldsymbol{\gamma}}, \boldsymbol{\Sigma}_{\boldsymbol{\gamma}})$, where $\mu_{\boldsymbol{\gamma}} = (\mu_1, \cdots, \mu_p)^{\mathrm{T}}$ and $\boldsymbol{\Sigma}_{\boldsymbol{\gamma}} = \mathrm{diag}((\sigma_1, \cdots, \sigma_p))$

The posterior distribution of $\alpha_k \sim \mathcal{N}(\widetilde{\mu}_k, \widetilde{\sigma}_k^2)$, where

$$
\begin{aligned}
-\frac{1}{2\widetilde{\sigma}_k^2} &= -\frac{1}{2}\frac{\widehat{\mathbf{R}}_{kk}}{\widehat{\mathbf{s}}_{\mathbf{\Gamma}k}^2} - \frac{1}{2\sigma_\alpha^2}, \\
\frac{\widetilde{\mu}_k}{\widetilde{\sigma}_k^2} &= \frac{\widehat{\Gamma}_k}{\widehat{\mathbf{s}}_{\mathbf{\Gamma}k}^2} - \frac{\beta_0}{\widehat{\mathbf{s}}_{\mathbf{\Gamma}k}}\sum_{i=1}^{p}\frac{\widehat{\mathbf{R}}_{ik}\langle\gamma_i\rangle}{\widehat{\mathbf{s}}_{\mathbf{\Gamma}i}} - \frac{1}{\widehat{\mathbf{s}}_{\mathbf{\Gamma}k}}\sum_{j\neq k}\frac{\langle\alpha_j\rangle\widehat{\mathbf{R}}_{jk}}{\widehat{\mathbf{s}}_{\mathbf{\Gamma}j}}, \qquad \text{(S8)}
\end{aligned}
$$

where, $\langle\alpha_j\rangle \stackrel{\text{def}}{=} E_q(\alpha_j)$.

Similarly, $\boldsymbol{\alpha} \sim \mathcal{N}(\boldsymbol{\alpha}|\mu_{\boldsymbol{\alpha}}, \boldsymbol{\Sigma}_{\boldsymbol{\alpha}})$, where $\mu_{\boldsymbol{\alpha}} = (\widetilde{\mu}_1, \cdots, \widetilde{\mu}_p)^{\mathrm{T}}$ and $\boldsymbol{\Sigma}_{\boldsymbol{\alpha}} = \mathrm{diag}((\widetilde{\sigma}_1, \cdots, \widetilde{\sigma}_p))$

##### 2.2.2 Variational M-step:

Given the current estimates of all parameters $\boldsymbol{\theta}$, the ELBO of the marginal log-likelihood can be written as follows

$$
\mathcal{L}(\boldsymbol{\theta}) = E_{q(\boldsymbol{\gamma},\boldsymbol{\alpha})}\left\{\log\Pr(\widehat{\mathbf{\Gamma}}, \widehat{\boldsymbol{\gamma}}, \boldsymbol{\alpha}, \boldsymbol{\gamma}|\widehat{\mathbf{S}}_{\boldsymbol{\gamma}}, \widehat{\mathbf{S}}_{\mathbf{\Gamma}}, \widehat{\mathbf{R}}; \boldsymbol{\theta})\right\} - E_{q(\boldsymbol{\gamma})}\left\{\log q(\boldsymbol{\gamma})\right\} - E_{q(\boldsymbol{\alpha})}\left\{\log q(\boldsymbol{\alpha})\right\}.
$$

Only the first term contains the parameters $\boldsymbol{\theta}$, which is the expectation of log-complete-data likelihood, in details

$$
\begin{aligned}
&E_{q(\boldsymbol{\gamma},\boldsymbol{\alpha})}\left\{\log\Pr(\widehat{\mathbf{\Gamma}}, \widehat{\boldsymbol{\gamma}}, \boldsymbol{\alpha}, \boldsymbol{\gamma}|\widehat{\mathbf{S}}_{\boldsymbol{\gamma}}, \widehat{\mathbf{S}}_{\mathbf{\Gamma}}, \widehat{\mathbf{R}}; \boldsymbol{\theta})\right\} \\
= &-\frac{1}{2}\beta_0^2\left\{\boldsymbol{\mu}_{\boldsymbol{\gamma}}^{\mathrm{T}}\widehat{\mathbf{S}}_{\mathbf{\Gamma}}^{-1}\widehat{\mathbf{R}}\widehat{\mathbf{S}}_{\mathbf{\Gamma}}^{-1}\boldsymbol{\mu}_{\boldsymbol{\gamma}} + Tr(\widehat{\mathbf{S}}_{\mathbf{\Gamma}}^{-1}\widehat{\mathbf{R}}\widehat{\mathbf{S}}_{\mathbf{\Gamma}}^{-1}\boldsymbol{\Sigma}_{\boldsymbol{\gamma}})\right\} - \frac{1}{2}\left\{\boldsymbol{\mu}_{\boldsymbol{\alpha}}^{\mathrm{T}}\widehat{\mathbf{S}}_{\mathbf{\Gamma}}^{-1}\widehat{\mathbf{R}}\widehat{\mathbf{S}}_{\mathbf{\Gamma}}^{-1}\boldsymbol{\mu}_{\boldsymbol{\alpha}} + Tr(\widehat{\mathbf{S}}_{\mathbf{\Gamma}}^{-1}\widehat{\mathbf{R}}\widehat{\mathbf{S}}_{\mathbf{\Gamma}}^{-1}\boldsymbol{\Sigma}_{\boldsymbol{\alpha}})\right\}
\end{aligned}
$$

$$
\begin{aligned}
- & \frac{1}{2}\left\{\xi^2\boldsymbol{\mu}_{\gamma}^{\mathrm{T}}\widehat{\mathbf{S}}_{\gamma}^{-1}\widehat{\mathbf{R}}\widehat{\mathbf{S}}_{\gamma}^{-1}\boldsymbol{\mu}_{\gamma} + Tr(\xi^2\widehat{\mathbf{S}}_{\gamma}^{-1}\widehat{\mathbf{R}}\widehat{\mathbf{S}}_{\gamma}^{-1}\boldsymbol{\Sigma}_{\gamma})\right\} + \beta_0\widehat{\boldsymbol{\Gamma}}^{\mathrm{T}}\widehat{\mathbf{S}}_{\boldsymbol{\Gamma}}^{-2}\boldsymbol{\mu}_{\gamma} - \beta_0\boldsymbol{\mu}_{\boldsymbol{\alpha}}^{\mathrm{T}}\widehat{\mathbf{S}}_{\boldsymbol{\Gamma}}^{-1}\widehat{\mathbf{R}}\widehat{\mathbf{S}}_{\boldsymbol{\Gamma}}^{-1}\boldsymbol{\mu}_{\gamma} \\
+ & \widehat{\boldsymbol{\Gamma}}^{\mathrm{T}}\widehat{\mathbf{S}}_{\boldsymbol{\Gamma}}^{-2}\boldsymbol{\mu}_{\boldsymbol{\alpha}} + \xi\widehat{\boldsymbol{\gamma}}^{\mathrm{T}}\widehat{\mathbf{S}}_{\gamma}^{-2}\boldsymbol{\mu}_{\gamma} - \frac{1}{2\sigma_{\boldsymbol{\gamma}}^2}\left\{\boldsymbol{\mu}_{\gamma}^{\mathrm{T}}\boldsymbol{\mu}_{\gamma} + Tr(\boldsymbol{\Sigma}_{\gamma})\right\} - \frac{1}{2\sigma_{\boldsymbol{\alpha}}^2}\left\{\boldsymbol{\mu}_{\boldsymbol{\alpha}}^{\mathrm{T}}\boldsymbol{\mu}_{\boldsymbol{\alpha}} + Tr(\boldsymbol{\Sigma}_{\boldsymbol{\alpha}})\right\} \\
- & \frac{p}{2}\ln\sigma_{\boldsymbol{\gamma}}^2 - \frac{p}{2}\ln\sigma_{\boldsymbol{\alpha}}^2 + const(\boldsymbol{\theta}). \qquad \text{(S9)}
\end{aligned}
$$

In the M-step, we obtain the new updates for all the parameters $\boldsymbol{\theta}$ by setting the derivative of $E_{q(\boldsymbol{\gamma},\boldsymbol{\alpha})}\left\{\log\Pr(\widehat{\boldsymbol{\Gamma}}, \widehat{\boldsymbol{\gamma}}, \boldsymbol{\alpha}, \boldsymbol{\gamma}|\widehat{\mathbf{S}}_{\gamma}, \widehat{\mathbf{S}}_{\boldsymbol{\Gamma}}, \widehat{\mathbf{R}}; \boldsymbol{\theta})\right\}$ to zeros. Thus, we have the following update equation

$$
\begin{aligned}
\beta_0 &= \left\{\boldsymbol{\mu}_{\gamma}^{\mathrm{T}}\widehat{\mathbf{S}}_{\boldsymbol{\Gamma}}^{-1}\widehat{\mathbf{R}}\widehat{\mathbf{S}}_{\boldsymbol{\Gamma}}^{-1}\boldsymbol{\mu}_{\gamma} + Tr(\widehat{\mathbf{S}}_{\boldsymbol{\Gamma}}^{-1}\widehat{\mathbf{R}}\widehat{\mathbf{S}}_{\boldsymbol{\Gamma}}^{-1}\boldsymbol{\Sigma}_{\gamma})\right\}^{-1}\left(\widehat{\boldsymbol{\Gamma}}^{\mathrm{T}}\widehat{\mathbf{S}}_{\boldsymbol{\Gamma}}^{-2}\boldsymbol{\mu}_{\gamma} - \boldsymbol{\mu}_{\boldsymbol{\alpha}}^{\mathrm{T}}\widehat{\mathbf{S}}_{\boldsymbol{\Gamma}}^{-1}\widehat{\mathbf{R}}\widehat{\mathbf{S}}_{\boldsymbol{\Gamma}}^{-1}\boldsymbol{\mu}_{\gamma}\right), \\
\sigma_{\boldsymbol{\gamma}}^2 &= \left\{\boldsymbol{\mu}_{\gamma}^{\mathrm{T}}\boldsymbol{\mu}_{\gamma} + Tr(\boldsymbol{\Sigma}_{\gamma})\right\}/p, \\
\sigma_{\boldsymbol{\alpha}}^2 &= \left\{\boldsymbol{\mu}_{\boldsymbol{\alpha}}^{\mathrm{T}}\boldsymbol{\mu}_{\boldsymbol{\alpha}} + Tr(\boldsymbol{\Sigma}_{\boldsymbol{\alpha}})\right\}/p, \\
\xi &= \left\{\boldsymbol{\mu}_{\gamma}^{\mathrm{T}}\widehat{\mathbf{S}}_{\gamma}^{-1}\widehat{\mathbf{R}}\widehat{\mathbf{S}}_{\gamma}^{-1}\boldsymbol{\mu}_{\gamma} + Tr(\widehat{\mathbf{S}}_{\gamma}^{-1}\widehat{\mathbf{R}}\widehat{\mathbf{S}}_{\gamma}^{-1}\boldsymbol{\Sigma}_{\gamma})\right\}^{-1}\left(\widehat{\boldsymbol{\gamma}}^{\mathrm{T}}\widehat{\mathbf{S}}_{\gamma}^{-2}\boldsymbol{\mu}_{\gamma}\right), \qquad \text{(S10)}
\end{aligned}
$$

where $\boldsymbol{\mu}_{\boldsymbol{\gamma}} = (\mu_1, \cdots, \mu_p)^{\mathrm{T}}, \boldsymbol{\Sigma}_{\boldsymbol{\gamma}} = \mathrm{diag}((\sigma_1^2, \cdots, \sigma_p^2)^{\mathrm{T}}), \boldsymbol{\mu}_{\boldsymbol{\alpha}} = (\widetilde{\mu}_1, \cdots, \widetilde{\mu}_p)^{\mathrm{T}}, \boldsymbol{\Sigma}_{\boldsymbol{\alpha}} = \mathrm{diag}((\widetilde{\sigma}_1^2, \cdots, \widetilde{\sigma}_p^2)^{\mathrm{T}})$. Since the entropy of $q(\boldsymbol{\gamma})$ equals $-\frac{p}{2}\ln(2\pi) - \frac{1}{2}\sum_{j=1}^{p}\ln\sigma_j^2 - \frac{p}{2}$ and the entropy of $q(\boldsymbol{\alpha})$ equals $-\frac{p}{2}\ln(2\pi) - \frac{1}{2}\sum_{k=1}^{p}\ln\widetilde{\sigma}_k^2 - \frac{p}{2}$, the ELBO can be written by

$$
\begin{aligned}
& \mathcal{L}(\boldsymbol{\theta}) \\
= & E_{q(\boldsymbol{\gamma},\boldsymbol{\alpha})}\left\{\log\Pr(\widehat{\boldsymbol{\Gamma}}, \widehat{\boldsymbol{\gamma}}, \boldsymbol{\alpha}, \boldsymbol{\gamma}|\widehat{\mathbf{S}}_{\gamma}, \widehat{\mathbf{S}}_{\boldsymbol{\Gamma}}, \widehat{\mathbf{R}}; \boldsymbol{\theta})\right\} - E_{q(\boldsymbol{\gamma})}\left\{\log q(\boldsymbol{\gamma})\right\} - E_{q(\boldsymbol{\alpha})}\left\{\log q(\boldsymbol{\alpha})\right\} \\
= & -\frac{1}{2}\beta_0^2\left\{\boldsymbol{\mu}_{\gamma}^{\mathrm{T}}\widehat{\mathbf{S}}_{\boldsymbol{\Gamma}}^{-1}\widehat{\mathbf{R}}\widehat{\mathbf{S}}_{\boldsymbol{\Gamma}}^{-1}\boldsymbol{\mu}_{\gamma} + Tr(\widehat{\mathbf{S}}_{\boldsymbol{\Gamma}}^{-1}\widehat{\mathbf{R}}\widehat{\mathbf{S}}_{\boldsymbol{\Gamma}}^{-1}\boldsymbol{\Sigma}_{\gamma})\right\} - \frac{1}{2}\left\{\boldsymbol{\mu}_{\boldsymbol{\alpha}}^{\mathrm{T}}\widehat{\mathbf{S}}_{\boldsymbol{\Gamma}}^{-1}\widehat{\mathbf{R}}\widehat{\mathbf{S}}_{\boldsymbol{\Gamma}}^{-1}\boldsymbol{\mu}_{\boldsymbol{\alpha}} + Tr(\widehat{\mathbf{S}}_{\boldsymbol{\Gamma}}^{-1}\widehat{\mathbf{R}}\widehat{\mathbf{S}}_{\boldsymbol{\Gamma}}^{-1}\boldsymbol{\Sigma}_{\boldsymbol{\alpha}})\right\} \\
- & \frac{1}{2}\left\{\xi^2\boldsymbol{\mu}_{\gamma}^{\mathrm{T}}\widehat{\mathbf{S}}_{\gamma}^{-1}\widehat{\mathbf{R}}\widehat{\mathbf{S}}_{\gamma}^{-1}\boldsymbol{\mu}_{\gamma} + Tr(\xi^2\widehat{\mathbf{S}}_{\gamma}^{-1}\widehat{\mathbf{R}}\widehat{\mathbf{S}}_{\gamma}^{-1}\boldsymbol{\Sigma}_{\gamma})\right\} + \beta_0\widehat{\boldsymbol{\Gamma}}^{\mathrm{T}}\widehat{\mathbf{S}}_{\boldsymbol{\Gamma}}^{-2}\boldsymbol{\mu}_{\gamma} - \beta_0\boldsymbol{\mu}_{\boldsymbol{\alpha}}^{\mathrm{T}}\widehat{\mathbf{S}}_{\boldsymbol{\Gamma}}^{-1}\widehat{\mathbf{R}}\widehat{\mathbf{S}}_{\boldsymbol{\Gamma}}^{-1}\boldsymbol{\mu}_{\gamma} \\
+ & \widehat{\boldsymbol{\Gamma}}^{\mathrm{T}}\widehat{\mathbf{S}}_{\boldsymbol{\Gamma}}^{-2}\boldsymbol{\mu}_{\boldsymbol{\alpha}} + \xi\widehat{\boldsymbol{\gamma}}^{\mathrm{T}}\widehat{\mathbf{S}}_{\gamma}^{-2}\boldsymbol{\mu}_{\gamma} - \frac{1}{2\sigma_{\boldsymbol{\gamma}}^2}\left\{\boldsymbol{\mu}_{\gamma}^{\mathrm{T}}\boldsymbol{\mu}_{\gamma} + Tr(\boldsymbol{\Sigma}_{\gamma})\right\} - \frac{1}{2\sigma_{\boldsymbol{\alpha}}^2}\left\{\boldsymbol{\mu}_{\boldsymbol{\alpha}}^{\mathrm{T}}\boldsymbol{\mu}_{\boldsymbol{\alpha}} + Tr(\boldsymbol{\Sigma}_{\boldsymbol{\alpha}})\right\} \\
- & \frac{p}{2}\ln\sigma_{\boldsymbol{\gamma}}^2 - \frac{p}{2}\ln\sigma_{\boldsymbol{\alpha}}^2 + \frac{1}{2}\sum_{j=1}^{p}\ln\sigma_j^2 + \frac{1}{2}\sum_{k=1}^{p}\ln\widetilde{\sigma}_k^2 + const(\boldsymbol{\theta}).
\end{aligned}
$$

The reduction steps: $\boldsymbol{\mu}_{\boldsymbol{\gamma}} = \xi\boldsymbol{\mu}_{\boldsymbol{\gamma}},\ \beta_0 = \beta_0/\xi,\ \sigma_{\boldsymbol{\gamma}}^2 = \xi^2\sigma_{\boldsymbol{\gamma}}^2.$

The corresponding PX-VBEM for MR-LDP can be summarized as Algorithm 2.

**Algorithm 2:** PX-VBEM for MR-LDP

*Initialization*: $\beta_0 = 0, \boldsymbol{\mu}_{\boldsymbol{\gamma}} = \boldsymbol{\mu}_{\boldsymbol{\alpha}} = \mathbf{0}$, $\sigma^2_{\boldsymbol{\alpha}} = \sigma^2_{\boldsymbol{\gamma}} = 0.01$ and $\xi = 1$.

**repeat**

  **E-step:** At the $t$-th iteration, for $j = 1, \cdots, p$ and $k = 1, \cdots, p$, both the posterior distribution of $q(\gamma_j|\boldsymbol{\theta}^{(t)})$ and $q(\alpha_k|\boldsymbol{\theta}^{(t)})$ are Gaussian with expressions (S7) and (S8). The parameters are given as $\xi = \xi^{(t)} = 1$, $\beta_0 = \beta_0^{(t)}$, $\sigma^2_{\boldsymbol{\gamma}} = (\sigma^{(t)}_{\boldsymbol{\gamma}})^2$ and $\sigma^2_{\boldsymbol{\alpha}} = (\sigma^{(t)}_{\boldsymbol{\alpha}})^2$.

  **M-tesp:** Update $\beta_0$, $\sigma^2_{\boldsymbol{\gamma}}$ , $\sigma^2_{\boldsymbol{\alpha}}$ and $\xi$ as equation(S10).

  **Reduction-step:** Rescale the parameters $\boldsymbol{\mu}_{\boldsymbol{\gamma}}^{(t+1)} = \xi^{(t+1)}\boldsymbol{\mu}_{\boldsymbol{\gamma}}^{(t+1)}$, $\beta_0^{(t+1)} = \beta_0^{(t+1)}/\xi^{(t+1)}$, $(\sigma_j^{(t+1)})^2 = (\xi^{(t+1)})^2(\sigma_j^{(t+1)})^2$ and reset $\xi^{(t+1)} = 1$.

**until** *convergence or maximum iteration reached*;

##### 2.2.3 Statistical Inference for MR-LDP

The procedure of statistical inference for MR-LDP is similar with MR-LD, we first calibrate the ELBO(denoted by $\widetilde{\mathcal{L}}(\boldsymbol{\theta}, \boldsymbol{\mu}_\gamma, \boldsymbol{\mu}_{\boldsymbol{\alpha}})$) as follows

$$
\begin{aligned}
& \widetilde{\mathcal{L}}(\boldsymbol{\theta}, \boldsymbol{\mu}_\gamma, \boldsymbol{\mu}_{\boldsymbol{\alpha}}) \\
= \;& E_{q(\boldsymbol{\gamma},\boldsymbol{\alpha})}\left\{\log \Pr(\widehat{\boldsymbol{\Gamma}}, \widehat{\boldsymbol{\gamma}}, \boldsymbol{\alpha}, \boldsymbol{\gamma}|\widehat{\mathbf{S}}_\gamma, \widehat{\mathbf{S}}_{\boldsymbol{\Gamma}}, \widehat{\mathbf{R}}; \boldsymbol{\theta})\right\} - E_{q(\boldsymbol{\gamma})}\left\{\log q(\boldsymbol{\gamma})\right\} - E_{q(\boldsymbol{\alpha})}\left\{\log q(\boldsymbol{\alpha})\right\} \\
= \;& \left(\beta_0 \widehat{\boldsymbol{\Gamma}}^{\mathrm{T}} \widehat{\mathbf{S}}_{\boldsymbol{\Gamma}}^{-2} + \xi \widehat{\boldsymbol{\gamma}} \widehat{\mathbf{S}}_\gamma^{-2}\right)\boldsymbol{\mu}_\gamma - \frac{1}{2}\boldsymbol{\mu}_\gamma^{\mathrm{T}} \widetilde{\boldsymbol{\Sigma}}_\gamma^{-1}\boldsymbol{\mu}_\gamma + \widehat{\boldsymbol{\Gamma}} \widehat{\mathbf{S}}_{\boldsymbol{\Gamma}}^{-2}\boldsymbol{\mu}_{\boldsymbol{\alpha}} - \frac{1}{2}\boldsymbol{\mu}_{\boldsymbol{\alpha}}^{\mathrm{T}} \widetilde{\boldsymbol{\Sigma}}_{\boldsymbol{\alpha}}^{-1}\boldsymbol{\mu}_{\boldsymbol{\alpha}} - \beta_0 \boldsymbol{\mu}_{\boldsymbol{\alpha}}^{\mathrm{T}} \widehat{\mathbf{S}}_{\boldsymbol{\Gamma}}^{-1}\widehat{\mathbf{R}}\widehat{\mathbf{S}}_{\boldsymbol{\Gamma}}^{-1}\boldsymbol{\mu}_\gamma \\
- \;& \frac{p}{2}(\log\sigma^2_{\boldsymbol{\gamma}} + 1) - \frac{p}{2}(\log\sigma^2_{\boldsymbol{\alpha}} + 1) + \frac{1}{2}\log|\widetilde{\boldsymbol{\Sigma}}_\gamma| + \frac{1}{2}\log|\widetilde{\boldsymbol{\Sigma}}_{\boldsymbol{\alpha}}| + p - p\log(2\pi) \\
- \;& \frac{1}{2}\left\{\log|\widehat{\mathbf{S}}_\gamma\widehat{\mathbf{R}}\widehat{\mathbf{S}}_\gamma| + \log|\widehat{\mathbf{S}}_{\boldsymbol{\Gamma}}\widehat{\mathbf{R}}\widehat{\mathbf{S}}_{\boldsymbol{\Gamma}}| + \widehat{\boldsymbol{\gamma}}^{\mathrm{T}}(\widehat{\mathbf{S}}_\gamma\widehat{\mathbf{R}}\widehat{\mathbf{S}}_\gamma)^{-1}\widehat{\boldsymbol{\gamma}} + \widehat{\boldsymbol{\Gamma}}^{\mathrm{T}}(\widehat{\mathbf{S}}_{\boldsymbol{\Gamma}}\widehat{\mathbf{R}}\widehat{\mathbf{S}}_{\boldsymbol{\Gamma}})^{-1}\boldsymbol{\Gamma}\right\}, \qquad \text{(S11)}
\end{aligned}
$$

where $\boldsymbol{\mu}_\gamma$ and $\widetilde{\boldsymbol{\Sigma}}_\gamma$ are the posterior mean and posterior variance for latent variable $\boldsymbol{\gamma}$, $\boldsymbol{\mu}_{\boldsymbol{\alpha}}$ and $\widetilde{\boldsymbol{\Sigma}}_{\boldsymbol{\alpha}}$ are the posterior mean and posterior variance for latent variable $\boldsymbol{\alpha}$, the forms of $\widetilde{\boldsymbol{\Sigma}}_\gamma$) and $\widetilde{\boldsymbol{\Sigma}}_{\boldsymbol{\alpha}}$) are from EM/PX-EM by plugging the posterior mean estimates and parameter estimates from VBEM/PX-VBEM which can be expressed as

$$
\widetilde{\boldsymbol{\Sigma}}_\gamma = \left(\beta_0^2 \widehat{\mathbf{S}}_{\boldsymbol{\Gamma}}^{-1}\widehat{\mathbf{R}}\widehat{\mathbf{S}}_{\boldsymbol{\Gamma}}^{-1} + \xi^2 \widehat{\mathbf{S}}_\gamma^{-1}\widehat{\mathbf{R}}\widehat{\mathbf{S}}_\gamma^{-1} + \sigma_{\boldsymbol{\gamma}}^{-2}\mathbf{I}\right)^{-1},
$$

and

$$\widetilde{\boldsymbol{\Sigma}}_{\boldsymbol{\alpha}} = \left(\widehat{\mathbf{S}}_{\boldsymbol{\Gamma}}^{-1}\widehat{\mathbf{R}}\widehat{\mathbf{S}}_{\boldsymbol{\Gamma}}^{-1} + \sigma_{\boldsymbol{\alpha}}^{-2}\mathbf{I}\right)^{-1}.$$

Similarly, we recalibrate the ELBO for MR-LDP as follows:

1. Obtain parameters together with variational means under both $\mathcal{H}_0$ and $\mathcal{H}_1$.

2. Calibrate ELBO using formulae (S11).

3. Calculate the test statistic (S4) using the calibrated ELBO as a proxy to the marginal log-likelihood.

### 3 More simulation results for different settings

The results of type-I error and point estimates for the dense pleiotropy with $n_3 = 2,500; 4,000$ are shown in Figure S2 - S3, respectively. For sparse horizontal pleiotropy, we consider the sparsity level at 0.2 and 0.4. Figure S4 and S5 show the results of type-I error and point estimates for the sparse pleiotropy at 0.2 with $n_3 = 2,500; 4,000$, respectively, and Figures-S6 - S8 show the results of type-I error and point estimates with $n_3 = 500; 2,500; 4,000$, respectively, for the sparse pleiotropy at 0.4.

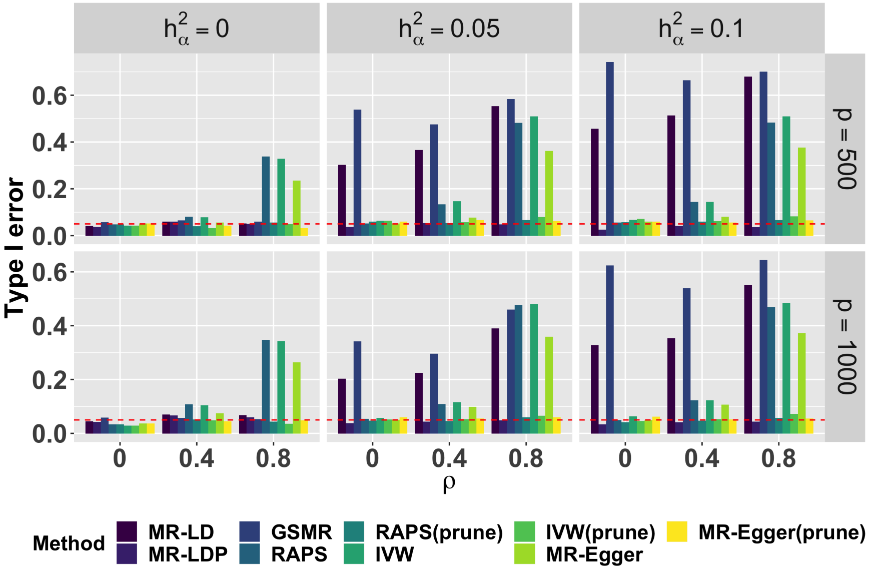

(A) Type-I error

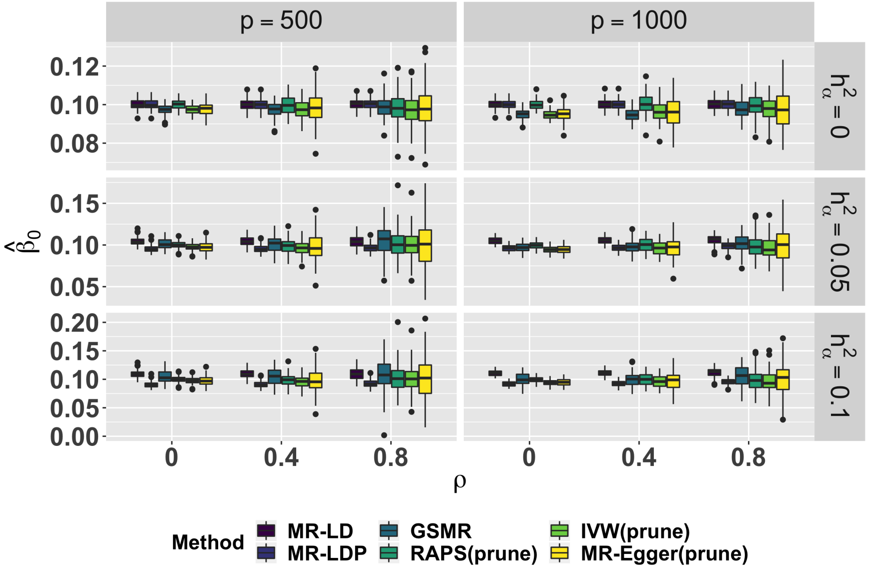

(B) Point estimates for $\beta_0$

Figure S2: Simulation of type-I error control and point estimates under the dense horizontal pleiotropy. $n_1 = n_2 = 20,000, n_3 = 2,500$, and the number of replication is 1000.

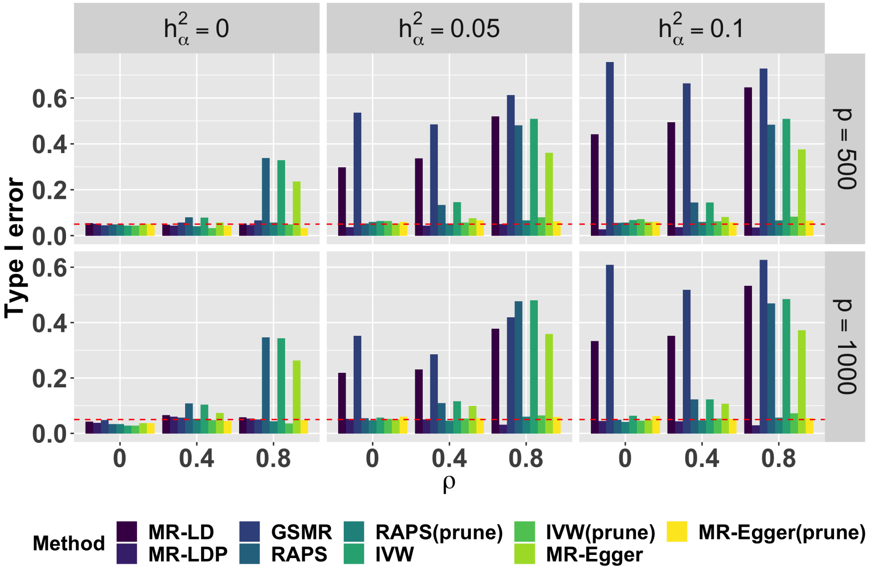

(A) Type-I error

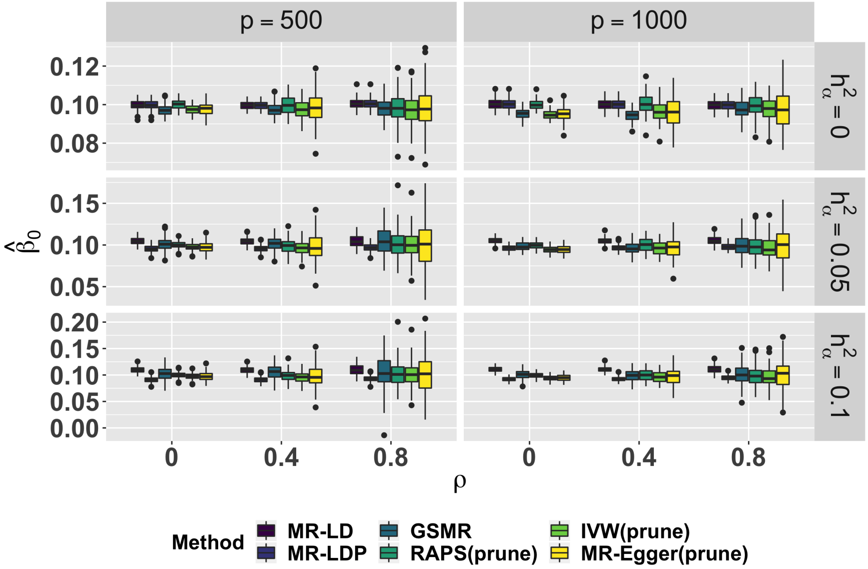

(B) Point estimates for $\beta_0$

Figure S3: Simulation of type-I error control and point estimates under the dense horizontal pleiotropy. $n_1 = n_2 = 20,000, n_3 = 4,000$, and the number of replication is 1000.

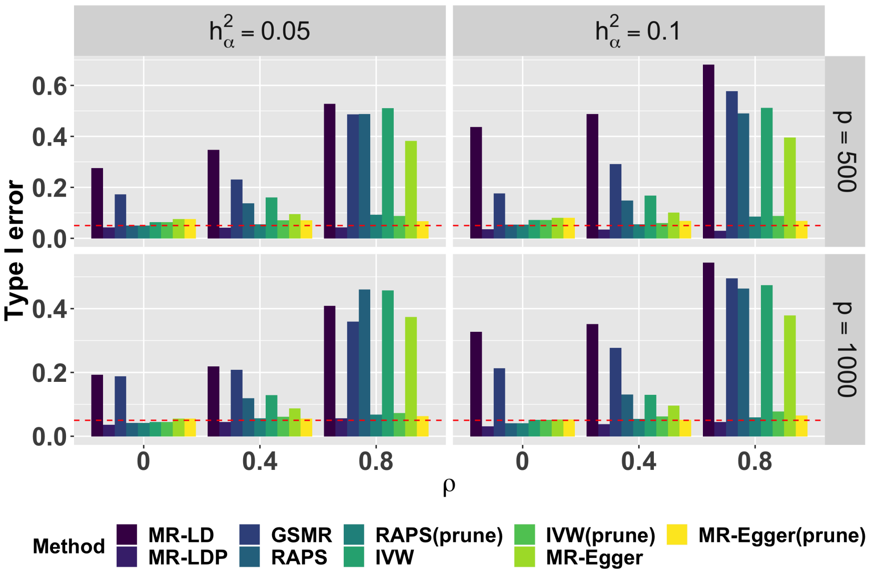

(A) Type-I error

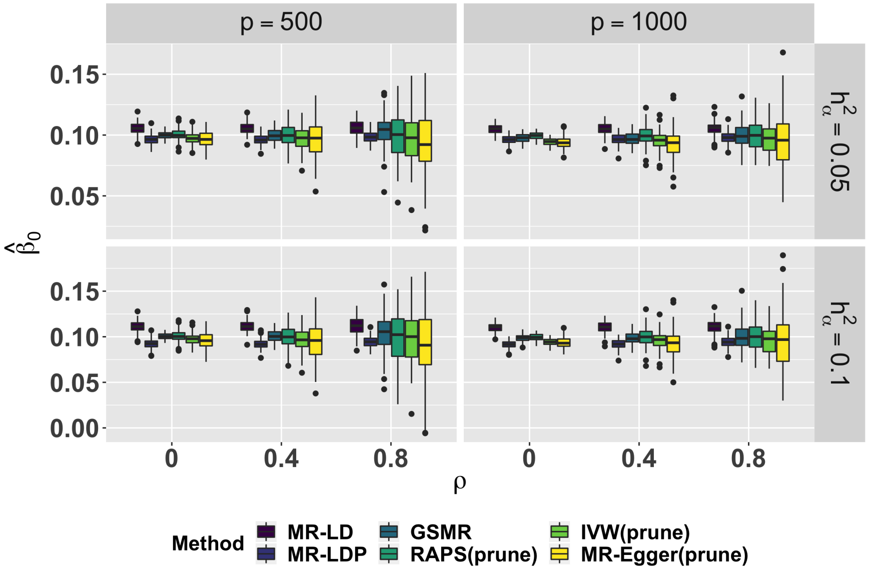

(B) Point estimates for $\beta_0$

Figure S4: Simulation of type-I error control and point estimates under the sparse horizontal pleiotropy, sparsity = 0.2. $n_1 = n_2 = 20,000, n_3 = 2,500.$, and the number of replication is 1000.

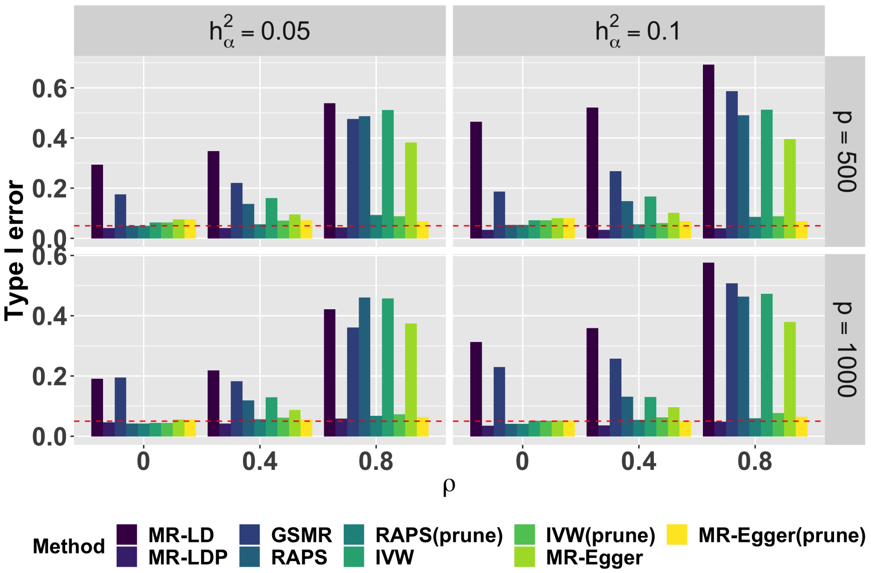

(A) Type-I error

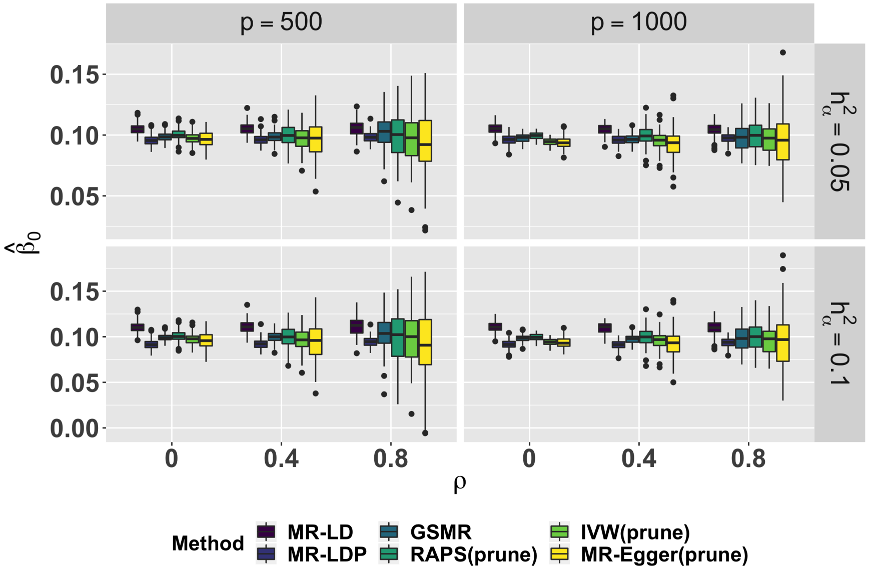

(B) Point estimates for $\beta_0$

Figure S5: Simulation of type-I error control and point estimates under the sparse horizontal pleiotropy, sparsity $= 0.2$. $n_1 = n_2 = 20,000, n_3 = 4,000.$, and the number of replication is 1000.

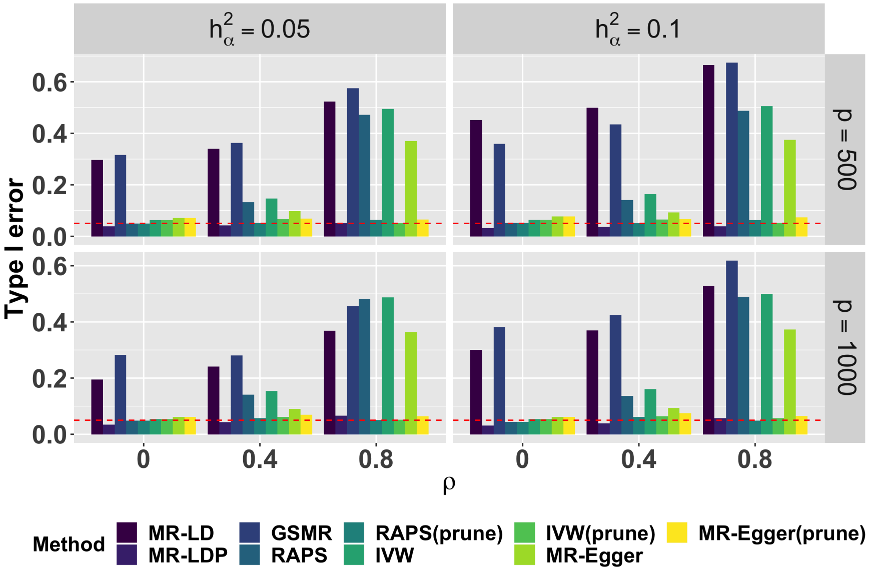

(A) Type-I error

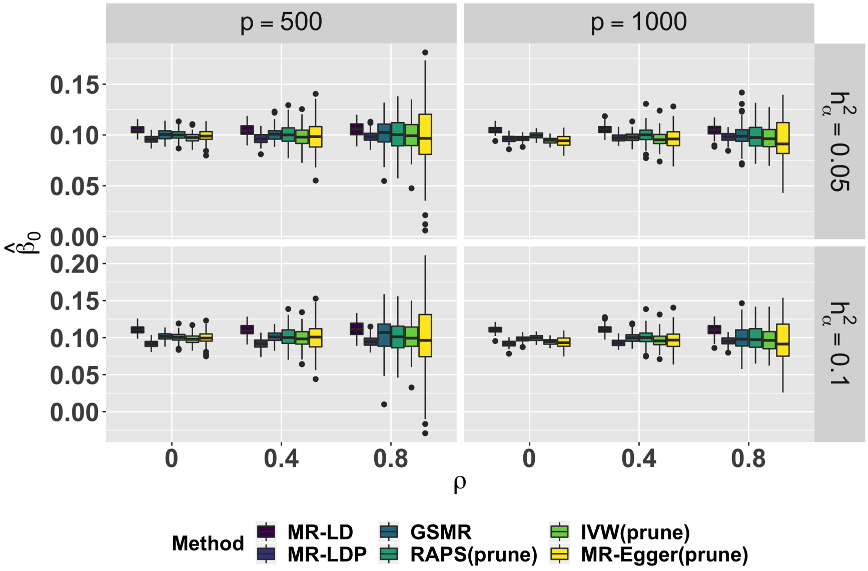

(B) Point estimates for $\beta_0$

Figure S6: Simulation of type-I error control and point estimates under the sparse horizontal pleiotropy, sparsity $= 0.4$. $n_1 = n_2 = 20,000, n_3 = 500.$, and the number of replication is 1000.

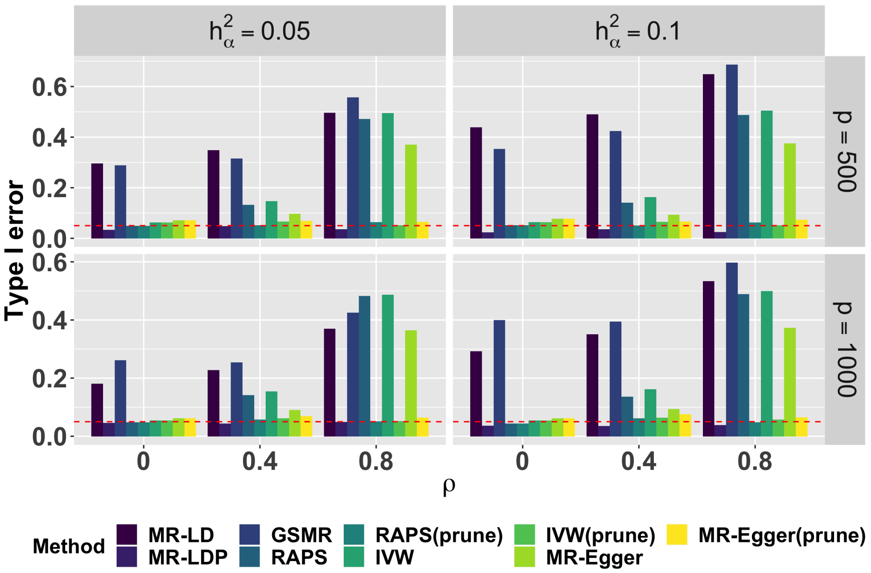

(A) Type-I error

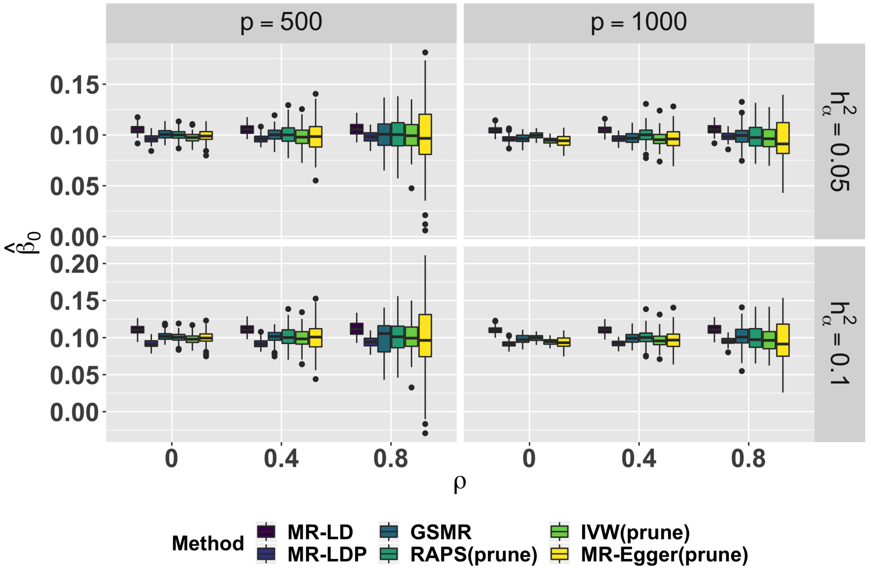

(B) Point estimates for $\beta_0$

Figure S7: Simulation of type-I error control and point estimates under the sparse horizontal pleiotropy, sparsity = 0.4. $n_1 = n_2 = 20,000, n_3 = 2,500.$, and the number of replication is 1000.

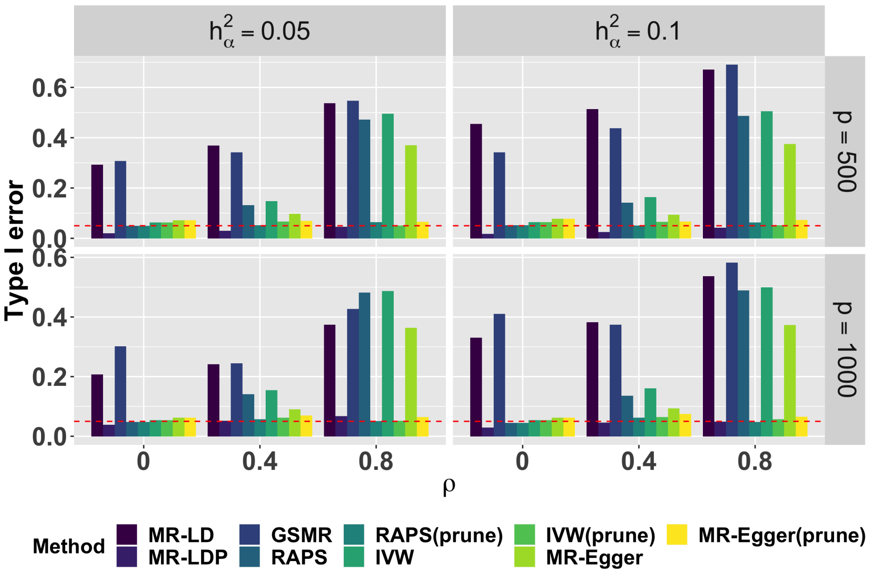

(A) Type-I error

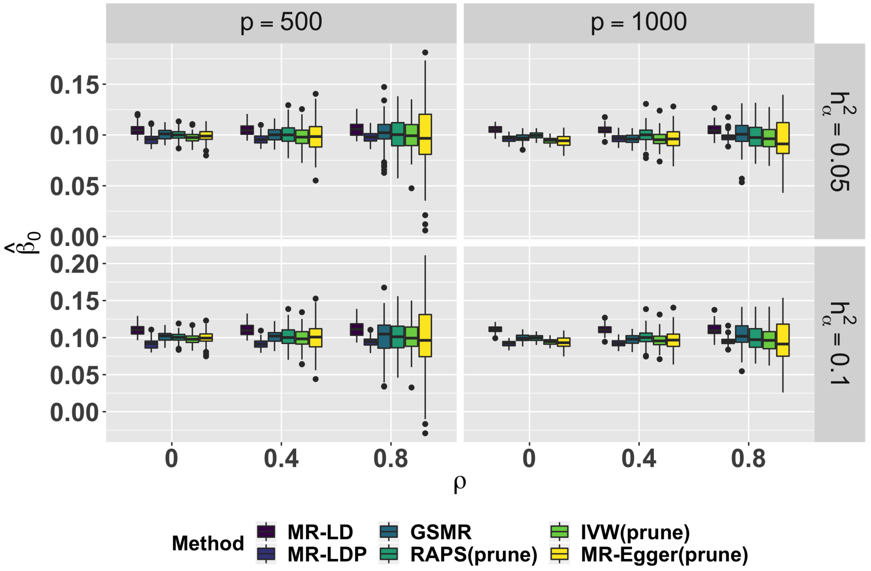

(B) Point estimates for $\beta_0$

Figure S8: Simulation of type-I error control and point estimates under the sparse horizontal pleiotropy, sparsity $= 0.4$. $n_1 = n_2 = 20,000, n_3 = 4,000.$, and the number of replication is 1000.

### 4 Real Data Analysis

#### 4.1 Two validation studies

##### 4.1.1 CAD-CAD study

We display the scatter plot of $\widehat{\boldsymbol{\gamma}}$ (C4D) against $\widehat{\boldsymbol{\Gamma}}$ (CAD1) in Figure S9, each point is augmented by the standard error of $\widehat{\gamma}_i$ and $\widehat{\Gamma}_i$ on the vertical and horizontal sides. In Figure S10, we report detailed results of CAD-CAD study with shrinkage parameters $\lambda = 0.15$.

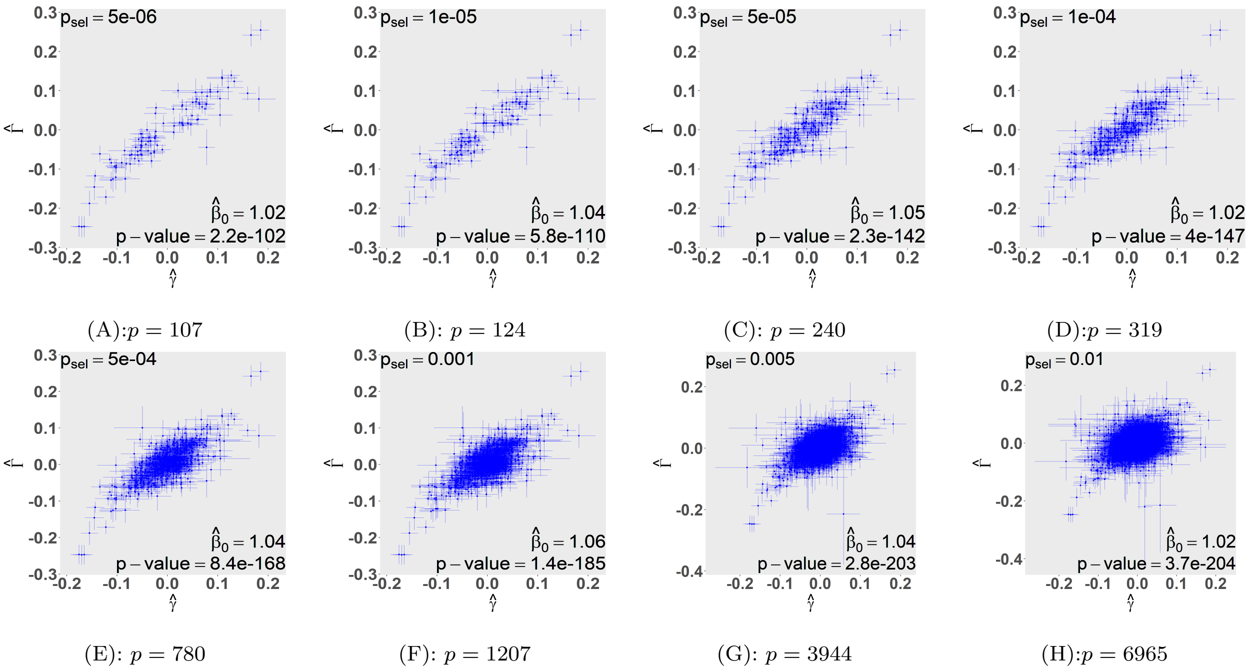

(A):$p = 107$ (B): $p = 124$ (C): $p = 240$ (D):$p = 319$

(E): $p = 780$ (F): $p = 1207$ (G): $p = 3944$ (H):$p = 6965$

Figure S9: Scatter plot for CAD-CAD study, where $\widehat{\beta}_0$ and the corresponding p-value were estimated by MR-LD using UK10K as the reference panel with $\lambda = 0.1$.

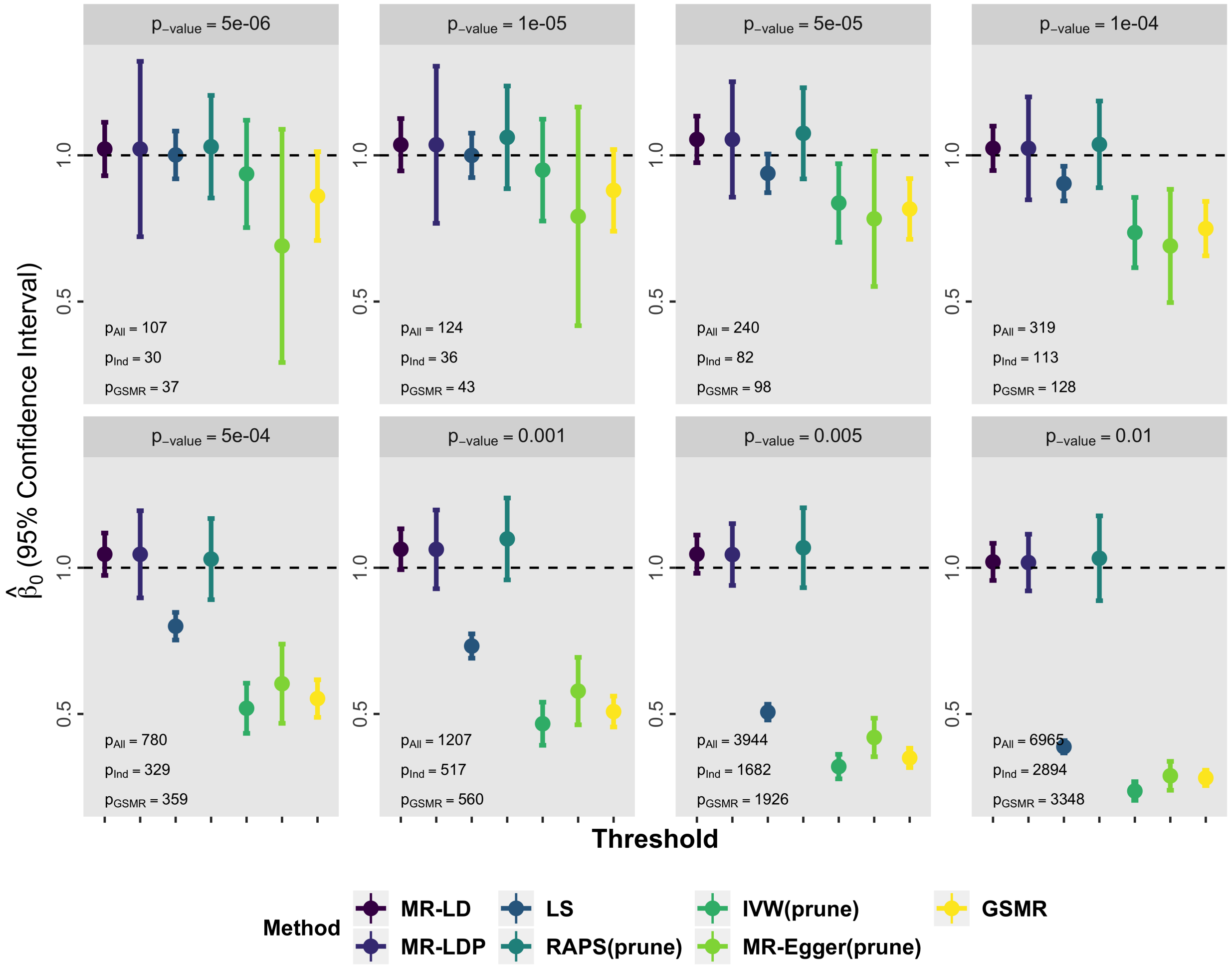

Figure S10: The result of CAD-CAD using UK10K as the reference penal with shrinkage parameter $\lambda = 0.15$. MR-LD, MR-LDP and LS methods use all SNPs selected by the screening dataset. The default value is used for thresholding $r^2$ in GSMR and the other three methods are 0.001.

##### 4.1.2 Height-Height study

We display the scatter plot of $\widehat{\boldsymbol{\gamma}}$ (height for males) against $\widehat{\boldsymbol{\Gamma}}$ (height for females) in Figure S11, each point is augmented by the standard error of $\widehat{\gamma}_i$ and $\widehat{\Gamma}_i$ on the vertical and horizontal sides. In Figures S12 and S13, we report detailed results of Height-Height study with shrinkage parameters $\lambda = 0.1$ and $\lambda = 0.15$, respectively.

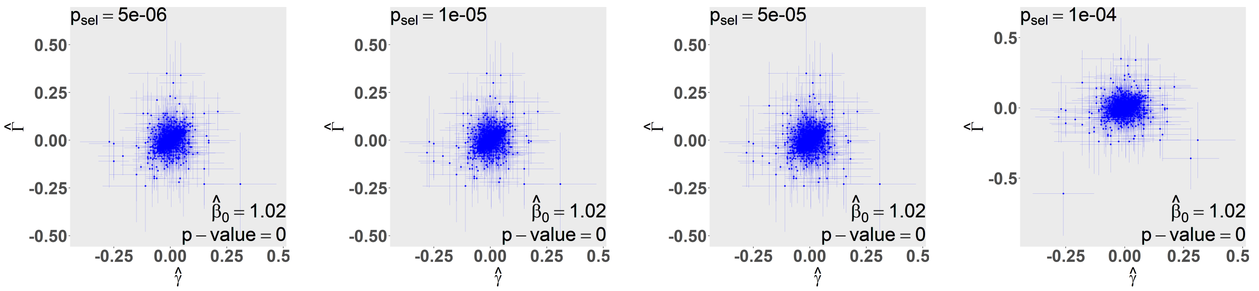

(A):$p = 7075$ (B): $p = 7557$ (C): $p = 8969$ (D):$p = 9734$

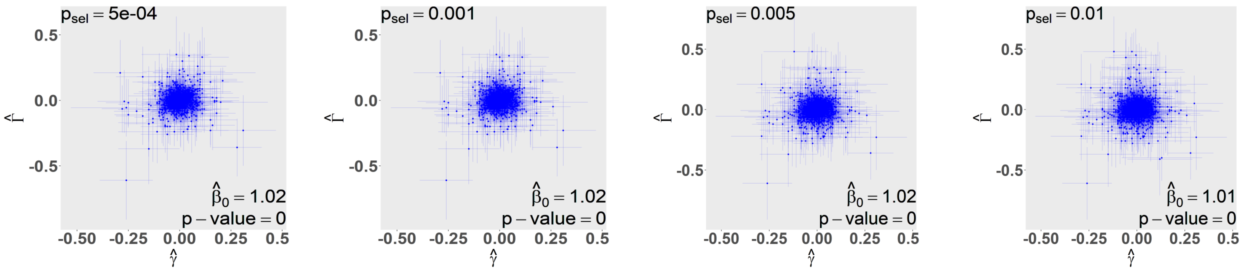

(E): $p = 12165$ (F): $p = 13539$ (G): $p = 18059$ (H):$p = 20947$

Figure S11: Scatter plot for Height-Height study, where $\widehat{\beta}_0$ and the corresponding p-value were estimated by MR-LD using UK10K as the reference panel with $\lambda = 0.1$.

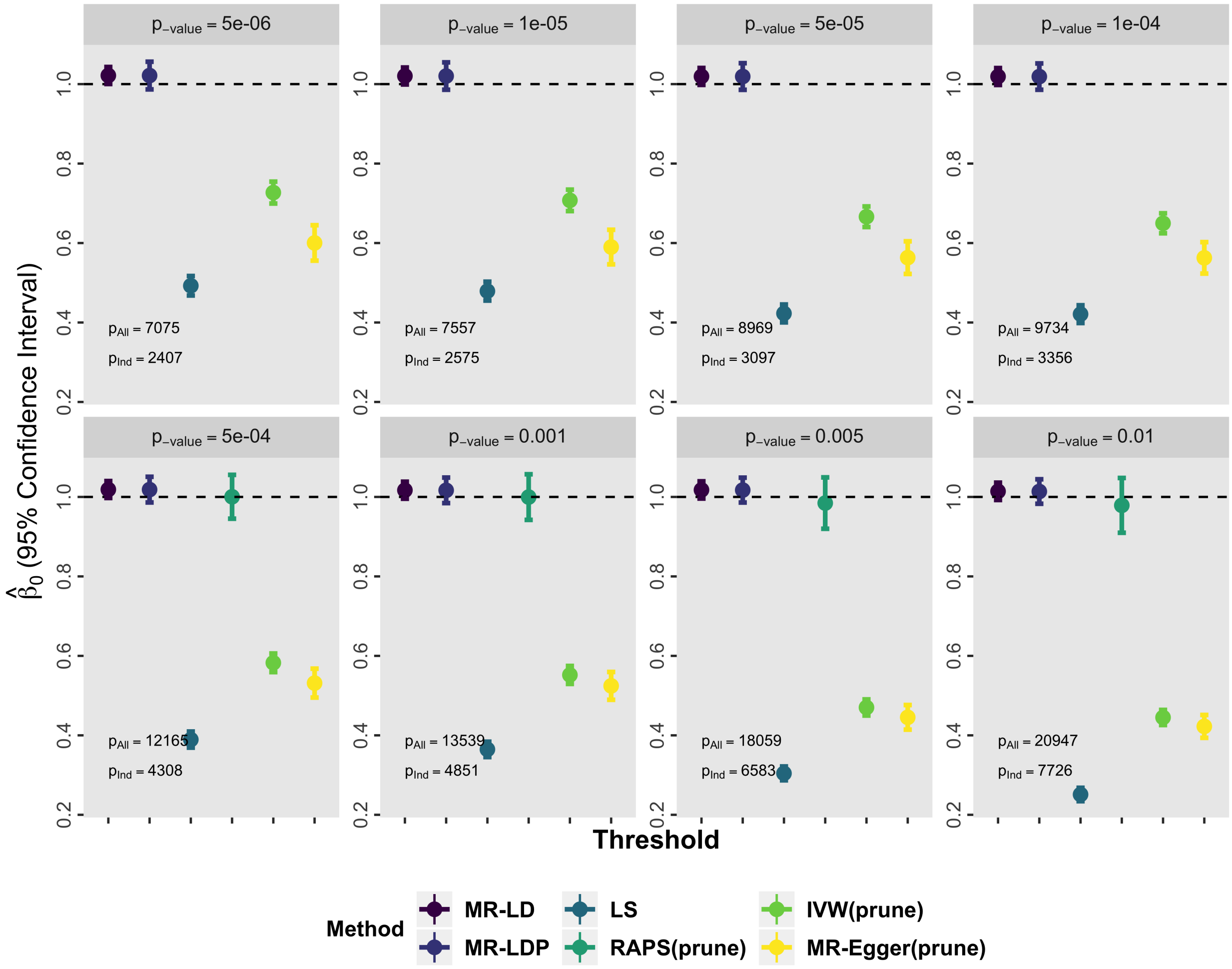

Figure S12: The result of Height-Height using UK10K as the reference penal with shrinkage parameter $\lambda = 0.1$. MR-LD, MR-LDP and LS methods use all SNPs selected by the screening dataset. The default value is used for thresholding $r^2$ in GSMR and the other three methods are 0.001.

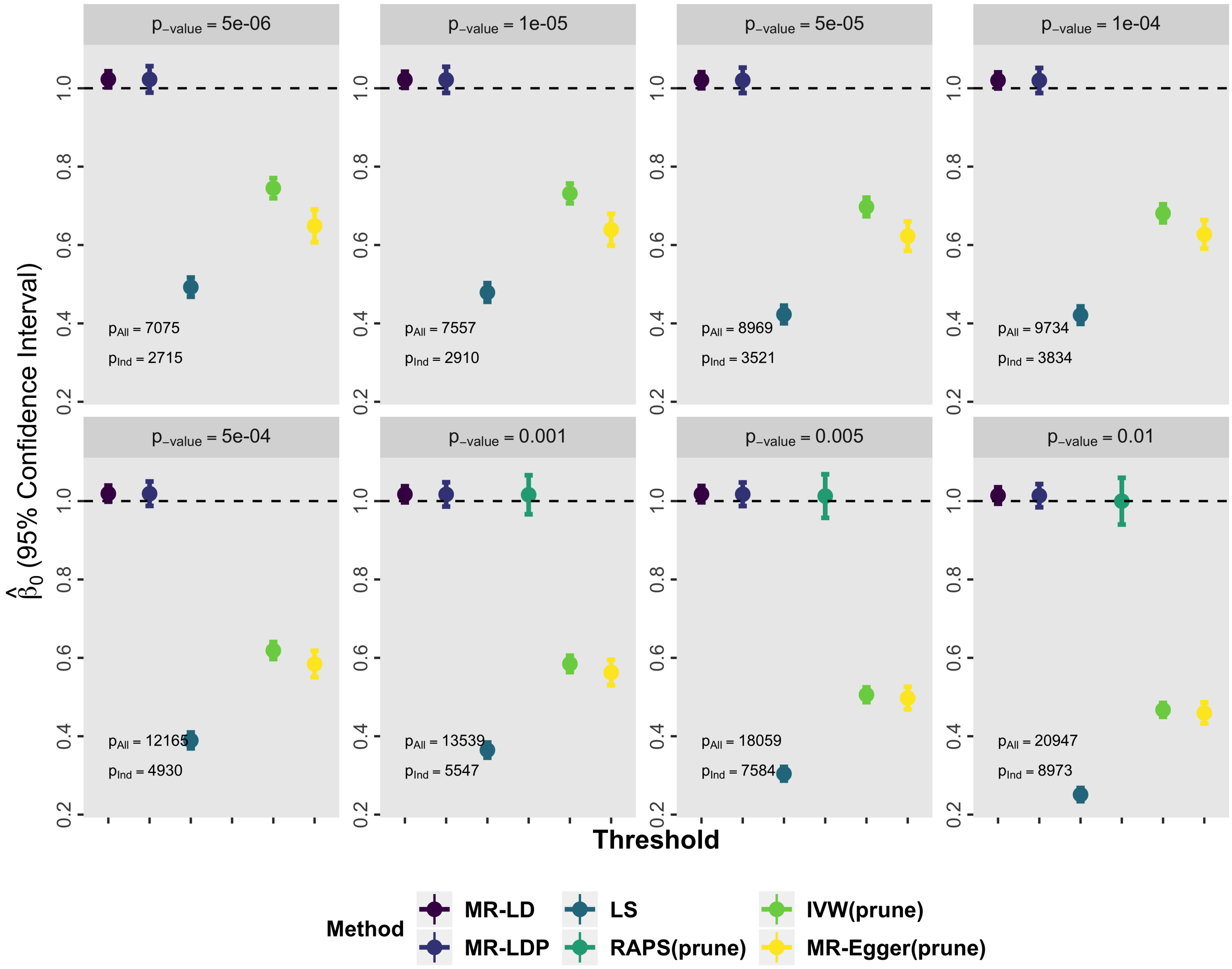

Figure S13: The result of Height-Height using UK10K as the reference penal with shrinkage parameter $\lambda = 0.15$. MR-LD, MR-LDP and LS methods use all SNPs selected by the screening dataset. The default value is used for thresholding $r^2$ in GSMR and the other three methods are 0.001.

#### 4.2 Applications to the effect of lipids and BMI on common diseases

##### 4.2.1 Lipids-disease outcome

The association result of lipids on common human diseases with shrinkage parameter $\lambda = 0.15$ is presented in Table S1, where the threshold for selecting instrumental variants in the selection dataset is set to $1 \times 10^{-4}$. Furthermore, we analysis the association of HDL-C on CAD1, CAD2, and PVD using a sequence of thresholds with different shrinkage parameters, the results are illustrated in Figures S14 and S18.

| Lipids | Outcome | $\#SNP_{ALL}$ | MRLDP | $\#SNP_{GSMR}$ | GSMR | $\#SNP_{LD}$ | Raps | IVW | MREGGER |
|---|---|---|---|---|---|---|---|---|---|
| HDL-C | CAD1 | 2104 | -0.16(0.032) | 299 | -0.27(0.036) | 222 | -0.38(0.065) | -0.36(0.064) | -0.26(0.139) |
| | CAD2 | 2071 | -0.09(0.023) | 307 | -0.05(0.029) | 226 | -0.11(0.045) | -0.12(0.043) | -0.01(0.088) |
| | MDD | 2071 | -0.06(0.031) | 310 | -0.14(0.038) | 226 | -0.13(0.056) | -0.13(0.053) | -0.11(0.109) |
| | T2D | 2071 | -0.14(0.036) | 304 | -0.19(0.042) | 226 | -0.3(0.073) | -0.31(0.074) | 0.07(0.149) |
| | Dyslid | 2071 | -0.17(0.027) | 283 | -0.1(0.029) | 226 | -0.27(0.082) | -0.27(0.072) | -0.23(0.148) |
| | Hyper | 2071 | -0.09(0.02) | 301 | -0.16(0.022) | 226 | -0.2(0.035) | -0.2(0.035) | -0.08(0.07) |
| | PVD | 2071 | -0.13(0.055) | 308 | -0.13(0.075) | 226 | -0.18(0.1) | -0.17(0.097) | 0.08(0.199) |
| | DC | 2071 | -0.06(0.011) | 300 | -0.08(0.013) | 226 | -0.08(0.023) | -0.09(0.023) | -0.02(0.046) |
| LDL-C | CAD1 | 1863 | 0.38(0.031) | 283 | 0.45(0.034) | 211 | 0.4(0.051) | 0.38(0.051) | 0.43(0.09) |
| | CAD2 | 1816 | 0.16(0.023) | 294 | 0.2(0.026) | 215 | 0.16(0.036) | 0.15(0.037) | 0.21(0.058) |
| | T2D | 1816 | -0.07(0.032) | 290 | -0.1(0.037) | 215 | -0.05(0.054) | -0.05(0.057) | -0.06(0.092) |
| | Dyslid | 1816 | 0.75(0.031) | 286 | 0.96(0.026) | 215 | 0.93(0.042) | 0.9(0.041) | 0.98(0.064) |
| | Hyper | 1816 | 0.05(0.018) | 289 | 0.06(0.019) | 215 | 0.04(0.028) | 0.04(0.029) | 0.05(0.045) |
| | Osteoa | 1816 | -0.05(0.025) | 295 | -0.05(0.028) | 215 | -0.08(0.036) | -0.07(0.036) | -0.07(0.057) |
| | DC | 1816 | 0.11(0.01) | 294 | 0.14(0.011) | 215 | 0.13(0.016) | 0.12(0.016) | 0.16(0.026) |
| TC | CAD1 | 2516 | 0.37(0.031) | 334 | 0.49(0.035) | 239 | 0.48(0.048) | 0.46(0.05) | 0.47(0.104) |
| | CAD2 | 2454 | 0.15(0.023) | 346 | 0.2(0.027) | 240 | 0.19(0.034) | 0.18(0.034) | 0.21(0.063) |
| | Dyslid | 2454 | 0.79(0.033) | 332 | 1.1(0.028) | 240 | 1.01(0.046) | 0.98(0.042) | 1.1(0.075) |
| | Osteoa | 2454 | -0.06(0.026) | 347 | -0.05(0.03) | 240 | -0.17(0.04) | -0.16(0.04) | -0.14(0.073) |
| | DC | 2454 | 0.1(0.011) | 345 | 0.14(0.012) | 240 | 0.12(0.017) | 0.12(0.017) | 0.14(0.031) |

Table S1: The causal associations of lipids on common diseases using UK10K as the reference penal with shrinkage parameter $\lambda = 0.15$. MRLDP and LS methods use all SNPs selected by the screening dataset. The thresholds of $r^2$ for GSMR and the other three methods are 0.05 and 0.001, respectively. Statistically significant results are shown in blue color.

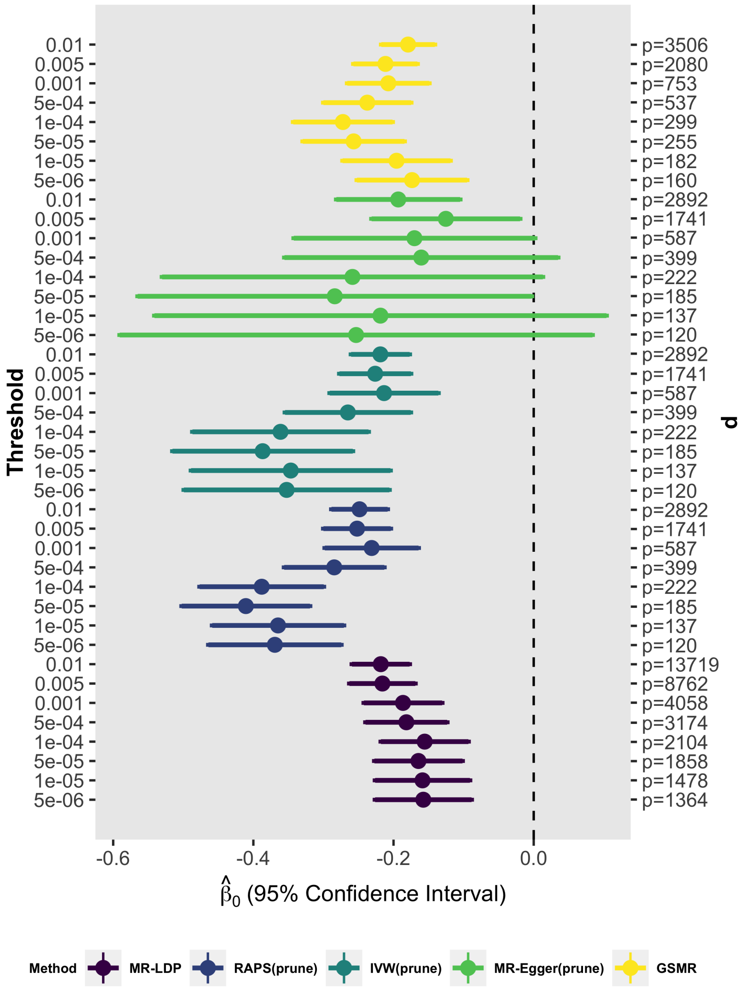

Figure S14: The causal associations of HDL-C on CAD1 under different thresholds using UK10K as the reference panel with $\lambda = 0.15$.

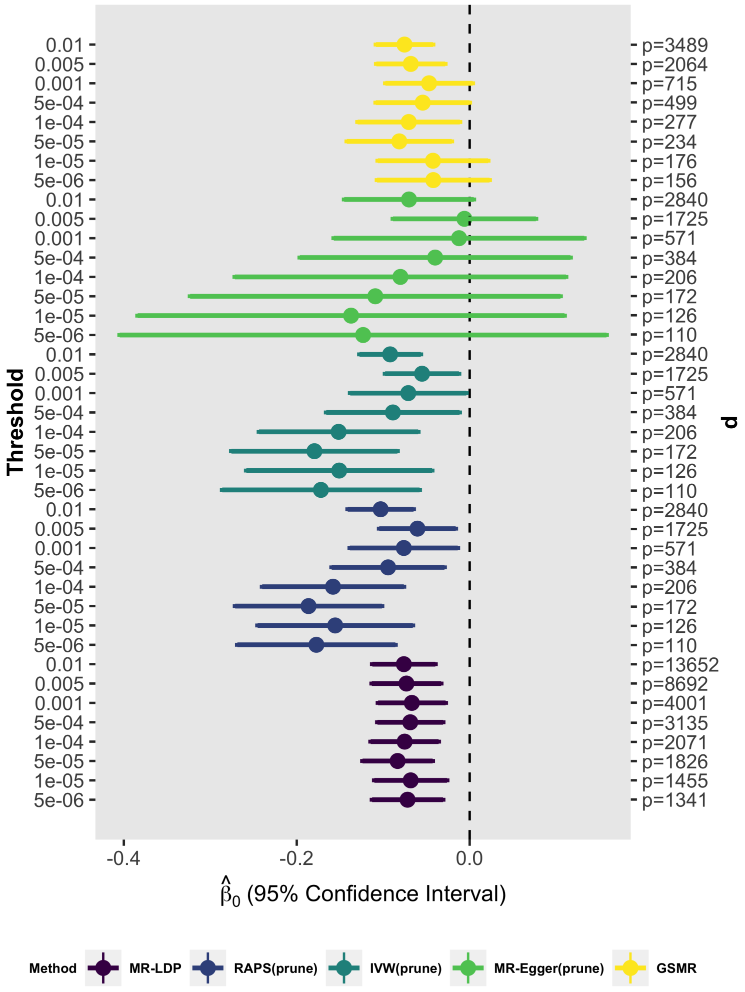

Figure S15: The causal associations of HDL-C on CAD2 under different thresholds using UK10K as the reference panel with $\lambda = 0.1$.

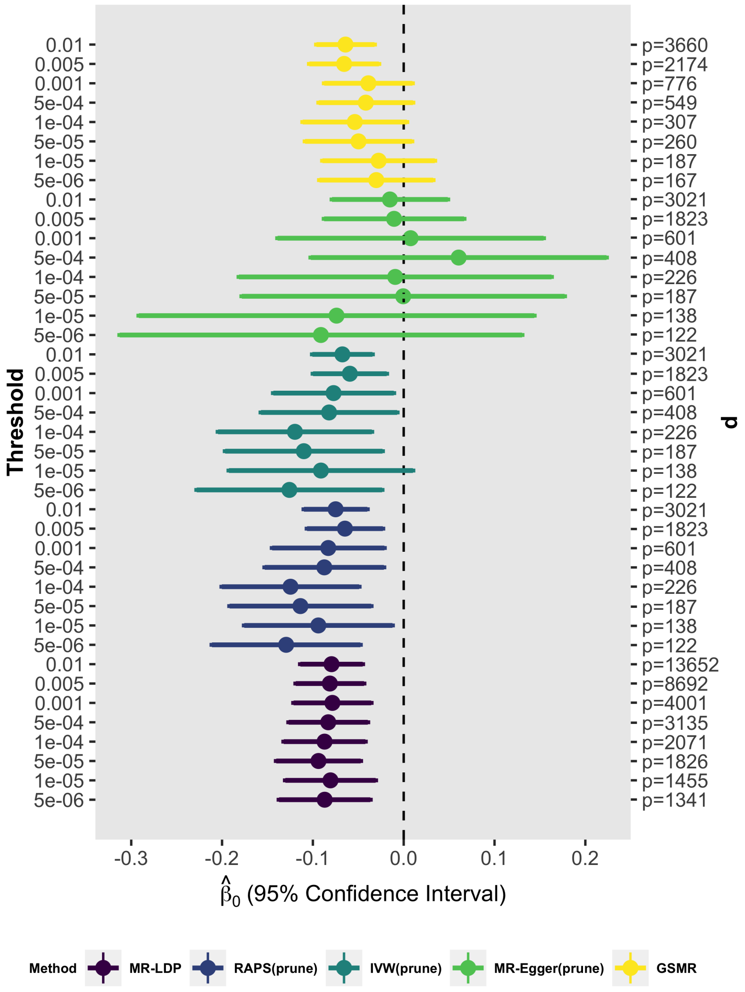

Figure S16: The causal associations of HDL-C on CAD2 under different thresholds using UK10K as the reference panel with $\lambda = 0.15$.

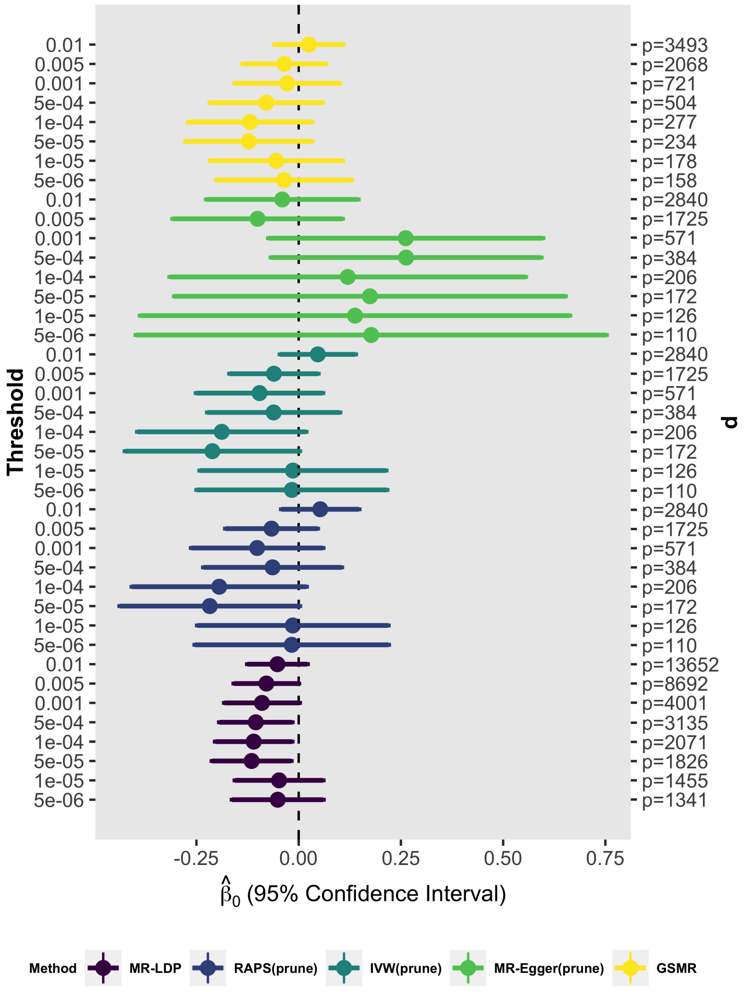

Figure S17: The causal associations of HDL-C on PVD under different thresholds using UK10K as the reference panel with $\lambda = 0.1$.

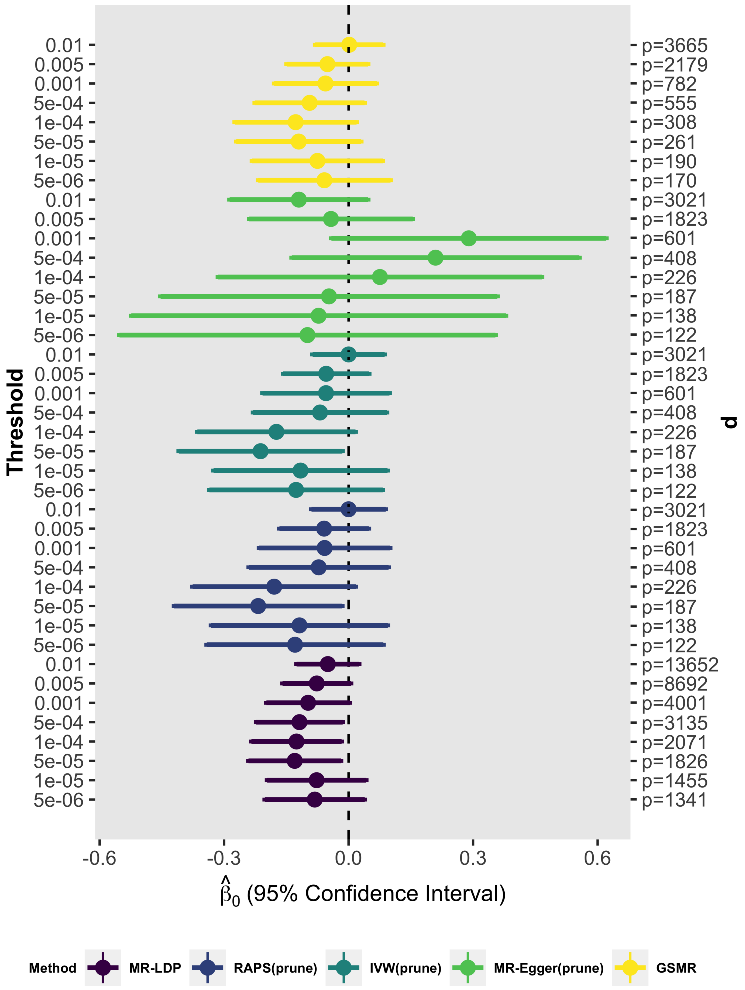

Figure S18: The causal associations of HDL-C on PVD under different thresholds using UK10K as the reference panel with $\lambda = 0.15$.

##### 4.2.2 BMI-disease outcome

Similarly, the association result of BMI on common human diseases with shrinkage parameter $\lambda = 0.15$ is showed in Table S2, where the threshold for selecting instrumental variants in the selection dataset is set to $1 \times 10^{-4}$. We also analysis the association of BMI on hemorrhoids and PVD using a sequence of thresholds with different shrinkage parameters, the results are illustrated in Figures S19 and S22.

| Outcome | $\#SNP_{ALL}$ | MR-LDP | $\#SNP_{GSMR}$ | GSMR | $\#SNP_{LD}$ | RAPS(prune) | IVW(prune) | MR-Egger(prune) |
|---|---|---|---|---|---|---|---|---|
| CAD1 | 4403 | 0.2(0.08) | 745 | 0.32(0.068) | 594 | 0.25(0.116) | 0.21(0.088) | 0.19(0.128) |
| Asthma | 4426 | 0.28(0.069) | 748 | 0.24(0.059) | 596 | 0.23(0.104) | 0.19(0.077) | 0.18(0.113) |
| CAD2 | 4426 | 0.22(0.064) | 749 | 0.22(0.06) | 596 | 0.19(0.101) | 0.15(0.076) | 0.08(0.111) |
| T2D | 4426 | 0.86(0.133) | 748 | 0.72(0.093) | 596 | 1.12(0.158) | 0.85(0.12) | 1.18(0.175) |
| Dyslid | 4426 | 0.22(0.072) | 745 | 0.28(0.057) | 596 | 0.12(0.125) | 0.12(0.083) | 0.21(0.121) |
| Hemorrhoids | 4426 | 0.31(0.128) | 750 | 0.24(0.108) | 596 | 0.19(0.165) | 0.15(0.124) | 0(0.182) |
| Hyper | 4426 | 0.47(0.063) | 744 | 0.47(0.045) | 596 | 0.58(0.092) | 0.45(0.066) | 0.53(0.097) |
| Insomnia | 4426 | 0.78(0.226) | 749 | 0.87(0.21) | 596 | 1.16(0.312) | 0.9(0.238) | 0.62(0.348) |
| Osteoa | 4426 | 0.27(0.075) | 750 | 0.28(0.067) | 596 | 0.29(0.106) | 0.22(0.079) | 0.4(0.115) |
| Osteop | 4426 | -0.44(0.17) | 750 | -0.39(0.146) | 596 | -0.53(0.232) | -0.41(0.173) | -0.52(0.253) |
| PVD | 4426 | 0.35(0.161) | 750 | 0.36(0.155) | 596 | 0.25(0.231) | 0.19(0.179) | 0.29(0.261) |
| DC | 4426 | 0.27(0.033) | 741 | 0.29(0.027) | 596 | 0.28(0.049) | 0.22(0.036) | 0.23(0.052) |

Table S2: The causal associations of BMI on common diseases using UK10K as the reference penal with shrinkage parameter $\lambda = 0.15$. MR-LDP uses all SNPs selected by the screening dataset. The thresholds of $r^2$ for GSMR and the other three methods are 0.05 and 0.001, respectively. Statistically significant results are shown in blue color.

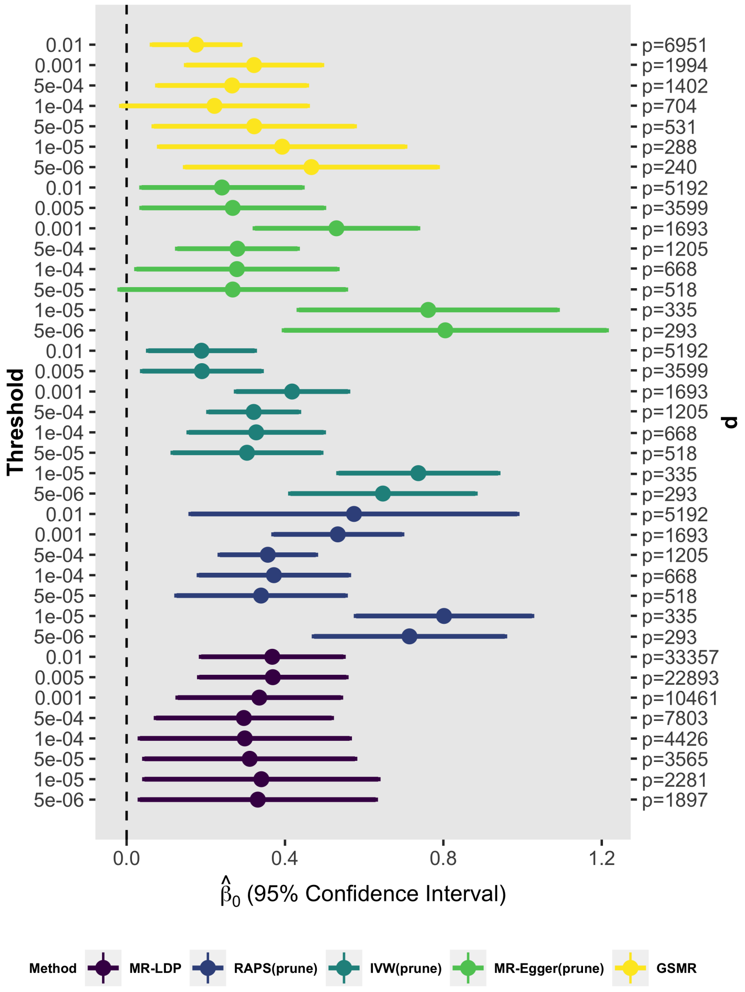

Figure S19: The causal associations of BMI on Hemorrhoids under different thresholds using UK10K as the reference panel with $\lambda = 0.1$.

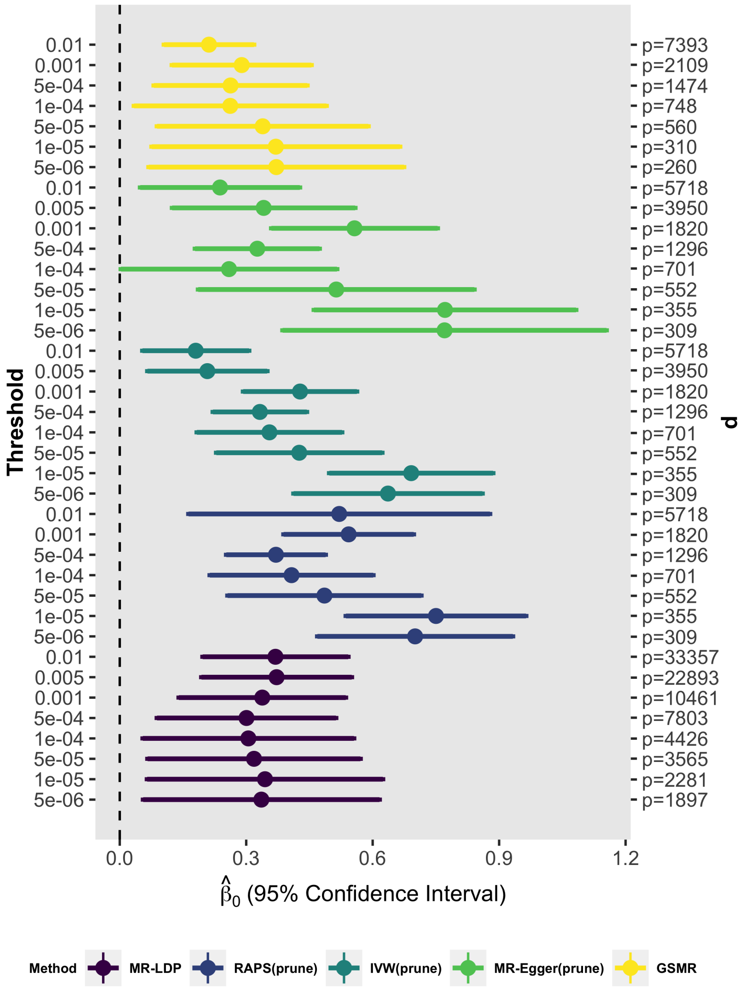

Figure S20: The causal associations of BMI on Hemorrhoids under different thresholds using UK10K as the reference panel with $\lambda = 0.15$.

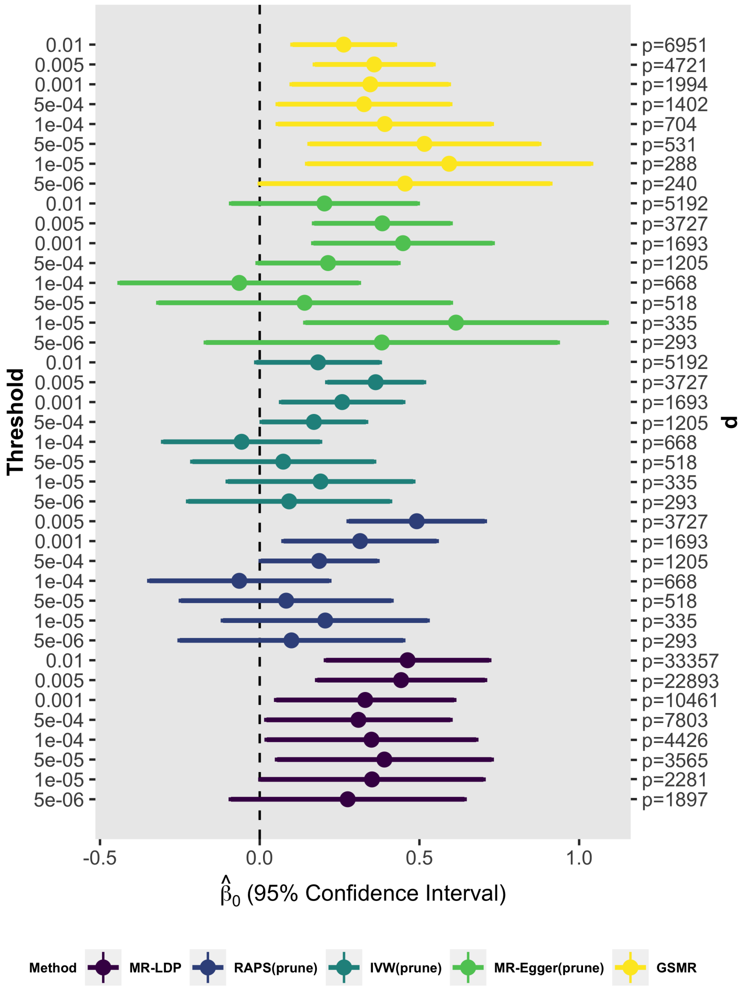

Figure S21: The causal associations of BMI on PVD under different thresholds using UK10K as the reference panel with $\lambda = 0.1$.

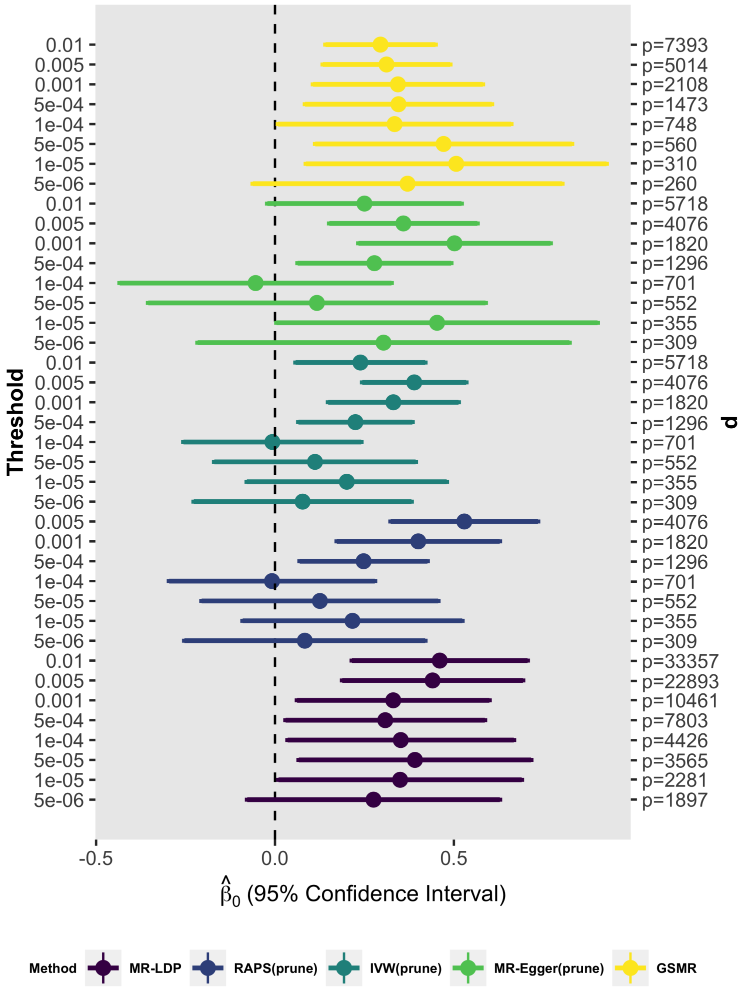

Figure S22: The causal associations of BMI on PVD under different thresholds using UK10K as the reference panel with $\lambda = 0.15$.

### 5 The detail information of GWAS Datasets

Tables S3 and S4 summarize the total number of SNPs and sample sizes for each trait in each health risk factor or disease outcome and the details for the sources of these GWAS summary statistics.

| ID | Year | Website |
|---|---|---|
| MI (screen) | 2018 | http://geneatlas.roslin.ed.ac.uk/downloads/ |
| C4D | 2011 | http://www.cardiogramplusc4d.org/data-downloads |
| CAD1 | 2011 | http://www.cardiogramplusc4d.org/data-downloads |
| CAD2 | 2018 | http://cnsgenomics.com/data.html |
| Height(screen) | 2018 | http://geneatlas.roslin.ed.ac.uk/downloads/ |
| Height-M | 2013 | https://portals.broadinstitute.org/collaboration/giant/index.php/Main_Page |
| Height-F | 2013 | https://portals.broadinstitute.org/collaboration/giant/index.php/Main_Page |
| BMI-JAP(screen) | 2017 | ftp://ftp.ebi.ac.uk/pub/databases/gwas/summary_statistics/AkiyamaM_28892062_GCST004904 |
| BMI | 2015 | https://portals.broadinstitute.org/collaboration/giant/index.php/Main_Page |
| HDL-c(screen) | 2010 | http://csg.sph.umich.edu/willer/public/lipids2010/ |
| LDL-c(screen) | 2010 | http://csg.sph.umich.edu/willer/public/lipids2010/ |
| TC(screen) | 2010 | http://csg.sph.umich.edu/willer/public/lipids2010/ |
| HDL-c | 2013 | http://csg.sph.umich.edu/willer/public/lipids2013/ |
| LDL-c | 2013 | http://csg.sph.umich.edu/willer/public/lipids2013/ |
| TC | 2013 | http://csg.sph.umich.edu/willer/public/lipids2013/ |
| Asthma | 2018 | http://cnsgenomics.com/data.html |
| AR | 2018 | http://cnsgenomics.com/data.html |
| Cancer | 2018 | http://cnsgenomics.com/data.html |
| MDD | 2018 | http://cnsgenomics.com/data.html |
| T2D | 2018 | http://cnsgenomics.com/data.html |
| Dyslid | 2018 | http://cnsgenomics.com/data.html |
| Hyper | 2018 | http://cnsgenomics.com/data.html |
| Hemorrhoids | 2018 | http://cnsgenomics.com/data.html |
| Hernia abdominopelvic cavity | 2018 | http://cnsgenomics.com/data.html |
| Insomnia | 2018 | http://cnsgenomics.com/data.html |
| IDA | 2018 | http://cnsgenomics.com/data.html |
| IBS | 2018 | http://cnsgenomics.com/data.html |
| Macular degeneration | 2018 | http://cnsgenomics.com/data.html |
| Osteoa | 2018 | http://cnsgenomics.com/data.html |
| Osteop | 2018 | http://cnsgenomics.com/data.html |
| PVD | 2018 | http://cnsgenomics.com/data.html |
| PU | 2018 | http://cnsgenomics.com/data.html |
| Pychiatric disorder | 2018 | http://cnsgenomics.com/data.html |
| Stress | 2018 | http://cnsgenomics.com/data.html |
| VV | 2018 | http://cnsgenomics.com/data.html |
| DC | 2018 | http://cnsgenomics.com/data.html |

Table S3: The website of publicly available summary datasets used in this paper.

| ID | Trait | Consortium | sample siz |
|---|---|---|---|
| MI (screen) | Myocardial infarction | UK BioBank | 30,358 |
| C4D | Coronary artery disease [11] | C4D Genetics Consortium | 30,442 |
| CAD1 | Coronary artery disease[12] | CARDIoGRAM | 86,995 |
| CAD2 | Coronary artery disease | UK BioBank | 108, 039 |
| WHRadjBMI-F(screen) | waist-to-hip ratio adjusted Body Mass Index in female in European group[13] | GIANT | 224,459 |
| Height(screen) | Height | UK BioBank | 500,062 |
| Height-M | Height in male in European group | [14] | 270,000 |
| Height-F | Height in female in European group | [14] | 270,000 |
| BMI-JAP(screen) | Body Mass Index[15] | [15] | 173,430 |
| BMI | Body Mass Index[16] in European group | Genetic Investigation of ANthropometric Traits(GIANT) | 322,154 |
| HDL-c(screen) | High-density lipoprotein cholesterol[17] | [17] | ∼100,000 |
| LDL-c(screen) | Low-density lipoprotein cholesterol[17] | [17] | ∼100,000 |
| TC(screen) | Total cholesterol[17] | [17] | ∼100,000 |
| HDL-C | High-density lipoprotein cholesterol[18] | Global Lipids Genetics Consortium(GLGC) | 188,577 |
| LDL-C | Low-density lipoprotein cholesterol[18] | Global Lipids Genetics Consortium(GLGC) | 188,577 |
| TC | Total cholesterol[18] | Global Lipids Genetics Consortium(GLGC) | 188,577 |
| Asthma | Asthma | UK BioBank | 108, 039 |
| AR | Allergic rhinitis | UK BioBank | 108, 039 |
| Cancer | Cancer | UK BioBank | 108,039 |
| MDD | Major depression disorder | UK BioBank | 108,039 |
| T2D | type 2 diabetes | UK BioBank | 108,039 |
| Dyslid | Dyslipidemia | UK BioBank | 108,039 |
| Hyper | Hypertension disease | UK BioBank | 108,039 |
| Hemorrhoids | Hemorrhoids | UK BioBank | 108,039 |
| Hernia abdominopelvic cavity | hernia abdominopelvic cavity | UK BioBank | 108,039 |
| Insomnia | Insomnia | UK BioBank | 108,039 |
| IDA | Iron deficiency anemias | UK BioBank | 108,039 |
| IBS | Irritable bowel syndrome | UK BioBank | 108,039 |
| Macular degeneration | Macular degeneration | UK BioBank | 108,039 |
| Osteoa | Osteoarthritis | UK BioBank | 108,039 |
| Osteop | Osteoporosis | UK BioBank | 108,039 |
| PVD | Peripheral vascular disease | UK BioBank | 108,039 |
| PU | Peptic ulcer | UK BioBank | 108,039 |
| Pychiatric disorder | Psychiatric disorder | UK BioBank | 108,039 |
| Stress | acute reaction to stress | UK BioBank | 108,039 |
| VV | varicose veins | UK BioBank | 108,039 |
| DC | disease count | UK BioBank | 108,039 |

Table S4: The Consortia of publicly available summary datasets used in this paper.
